# Capturing and Recreating Diverse Antibody Repertoires as Multivalent Recombinant Polyclonal Antibody Drugs

**DOI:** 10.1101/2020.08.05.232975

**Authors:** Sheila M. Keating, Rena A. Mizrahi, Matthew S. Adams, Michael A. Asensio, Emily Benzie, Kyle P. Carter, Yao Chiang, Robert C. Edgar, Bishal K. Gautam, Ashley Gras, Jackson Leong, Renee Leong, Yoong Wearn Lim, Vishal A. Manickam, Angelica V. Medina-Cucurella, Ariel R. Niedecken, Jasmeen Saini, Jan Fredrik Simons, Matthew J. Spindler, Kacy Stadtmiller, Brendan Tinsley, Ellen K. Wagner, Nicholas Wayham, LaRee Tracy, Carina Vingsbo Lundberg, Dirk Büscher, Jose Vicente Terencio, Lucy Roalfe, Emma Pearce, Hayley Richardson, David Goldblatt, Anushka T. Ramjag, Christine V.F. Carrington, Graham Simmons, Marcus O. Muench, Steven M. Chamow, Bryan Monroe, Charles Olson, Thomas H. Oguin, Heather Lynch, Robert Jeanfreau, Everett H. Meyer, Adam S. Adler, David S. Johnson

## Abstract

Plasma-derived polyclonal antibodies are polyvalent drugs used for many important clinical indications that require modulation of multiple drug targets simultaneously, including emerging infectious disease and transplantation. However, plasma-derived drugs suffer many problems, including low potency, impurities, constraints on supply, and batch-to-batch variation. In this study, we demonstrated proofs-of-concept for a technology that uses microfluidics and molecular genomics to capture diverse mammalian antibody repertoires as multivalent recombinant drugs. These “recombinant hyperimmune” drugs comprised thousands to tens of thousands of antibodies and were derived from convalescent human donors, or vaccinated human donors or immunized mice. Here we used our technology to build a highly potent recombinant hyperimmune for Severe Acute Respiratory Syndrome Coronavirus-2 (SARS CoV-2) in less than three months. We also validated a recombinant hyperimmune for Zika virus disease that abrogates antibody-dependent enhancement (ADE) through Fc engineering. For patients with primary immune deficiency (PID), we built high potency polyvalent recombinant hyperimmunes against pathogens that commonly cause serious lung infections. Finally, to address the limitations of rabbit-derived anti-thymocyte globulin (ATG), we generated a recombinant human version and demonstrated in vivo function against graft-versus-host disease (GVHD). Recombinant hyperimmunes are a novel class of drugs that could be used to target a wide variety of other clinical applications, including cancer and autoimmunity.

## INTRODUCTION

Many diseases are best treated by drugs that target multiple epitopes, for example, infectious viruses or bacteria with many variants or serotypes. Such multi-target diseases are treated with polyspecific (polyvalent) antibodies derived from human or animal plasma, such as intravenous immunoglobulin (IVIG).^1^ Higher-potency polyclonal antibody drugs known as “hyperimmunes” are often derived from the plasma of recently vaccinated human donors, for example, HepaGam B against Hepatitis B Virus (HBV)^2^ and BabyBIG against infant botulism.^3^ In diseases for which human vaccination is not possible, hyperimmunes can be generated by immunizing animals, for example, rabbit-derived Thymoglobulin against human thymocytes for transplant tolerance.^4^ For rapid response to emerging pathogens with poorly characterized neutralizing epitopes, many groups have developed hyperimmunes derived from immunized animal plasma or convalescent human serum, for example, Zika virus hyperimmune^5^ or Severe Acute Respiratory Syndrome Coronavirus-2 (SARS CoV-2).^6–7^

Plasma-derived antibody therapeutics have significant drawbacks. First, demand for normal and convalescent donor plasma often outstrips supply.^8^ Plasma-derived drugs have in the past suffered from impurities, including infectious viruses and clotting factors, that have resulted in serious adverse events.^9–10^ Antibody drugs derived from animal plasma occasionally cause allergic reactions,^11^ lead to anti-drug antibodies, and have suboptimal effector properties.^12^ Because they are derived from naturally occurring proteins, plasma-derived drugs are not easily engineered, for example it is not possible to modify Fc sequences to improve mechanism of action or drug half-life. Finally, each batch of plasma-derived drug is usually derived from a different cohort of human donors or animals, resulting in batch-to-batch variation.^13–15^

Generating multivalent hyperimmunes using recombinant DNA technology could solve many of the problems inherent to plasma-derived hyperimmunes. However, manufacturing recombinant polyclonal antibodies presents substantial technical hurdles. Most importantly, a recombinant hyperimmune technology would have to isolate significant numbers of B cells from donors or animals, natively pair heavy and light chain immunoglobulin (Ig) at a single cell level, and then clone the sequences into recombinant expression libraries for manufacturing. Conventionally, most production cell lines for recombinant antibody drugs are generated by random integration of expression constructs into mammalian cell genomes.^16^ To prevent mispairing between heavy and light chain Ig, a recombinant polyclonal hyperimmune technology would require a single genome integration site. Pioneering work used 96-well plates to capture antibody sequences from B cells isolated from human donors immunized with Rho(D)+ erythrocytes and then engineer polyvalent recombinant antibodies,^17^ but this approach produced drug candidates with fewer than 30 antibodies, complicating broad application and reducing potential for polyvalence.

Here we describe a novel technology that has the potential to solve unmet clinical problems in the field by generating diverse recombinant hyperimmunes through modern high-throughput microfluidics, genomics, and mammalian cell engineering. B cells from human donors or mice were run through a microfluidic platform, heavy and light chain Ig nucleic acid sequences were fused on a single-cell level to create antibody repertoires,^18^ antibody repertoires were engineered into full-length expression constructs *en masse*, and then the full-length antibody expression constructs were stably introduced *en masse* into Chinese hamster ovary (CHO) cells in a site-directed manner. We applied our technology to develop 10^3^- to 10^4^-diverse polyvalent recombinant hyperimmune drug candidates that address unmet clinical needs for the COVID-19 pandemic, Zika virus disease, primary immune deficiency (PID), and transplant tolerance. We showed *in vivo* and/or *in vitro* validation of a drug candidate for each of the four clinical applications and discuss how this novel class of drugs might be broadly applicable for other emerging and existing clinical problems.

## RESULTS

### Capturing Diverse Antibody Repertoires as CHO Libraries

Mammalian antibody repertoires are extremely diverse, comprising as many as 10^7^ antibody clonotypes.^19^ Advanced molecular technology is required to capture a substantial fraction of a mammalian donor’s diverse antibody repertoire. Previously, we reported methods for generating millions-diverse libraries of natively paired heavy and light chain Ig sequences in yeast.^18^ That method used microfluidics to isolate millions of single B cells per hour into picoliter droplets for lysis, followed by overlap extension reverse transcriptase polymerase chain reaction (OE-RT-PCR), to generate libraries of natively paired single chain variable fragments (scFv).

Because antibody repertoires often comprise many antibodies not directed against the target(s) of interest, we employed a variety of enrichment methods (**Figure 1**). For ATG, Zika virus, *Haemophilus influenzae* b (Hib), and *Streptococcus pneumoniae* (pneumococcus), we administered immunogens to human donors or humanized mice prior to sampling antibody-producing cells. For SARS CoV-2, we recruited convalescent donors who recently tested positive for COVID-19, made yeast display scFv libraries from donor B cells, and sorted the libraries derived from these donors to enrich for antibodies directed against SARS CoV-2 antigen. In all cases, the output was a library of thousands to tens of thousands of natively paired scFv DNAs, enriched for activity against their respective target(s).

**Figure 1.**
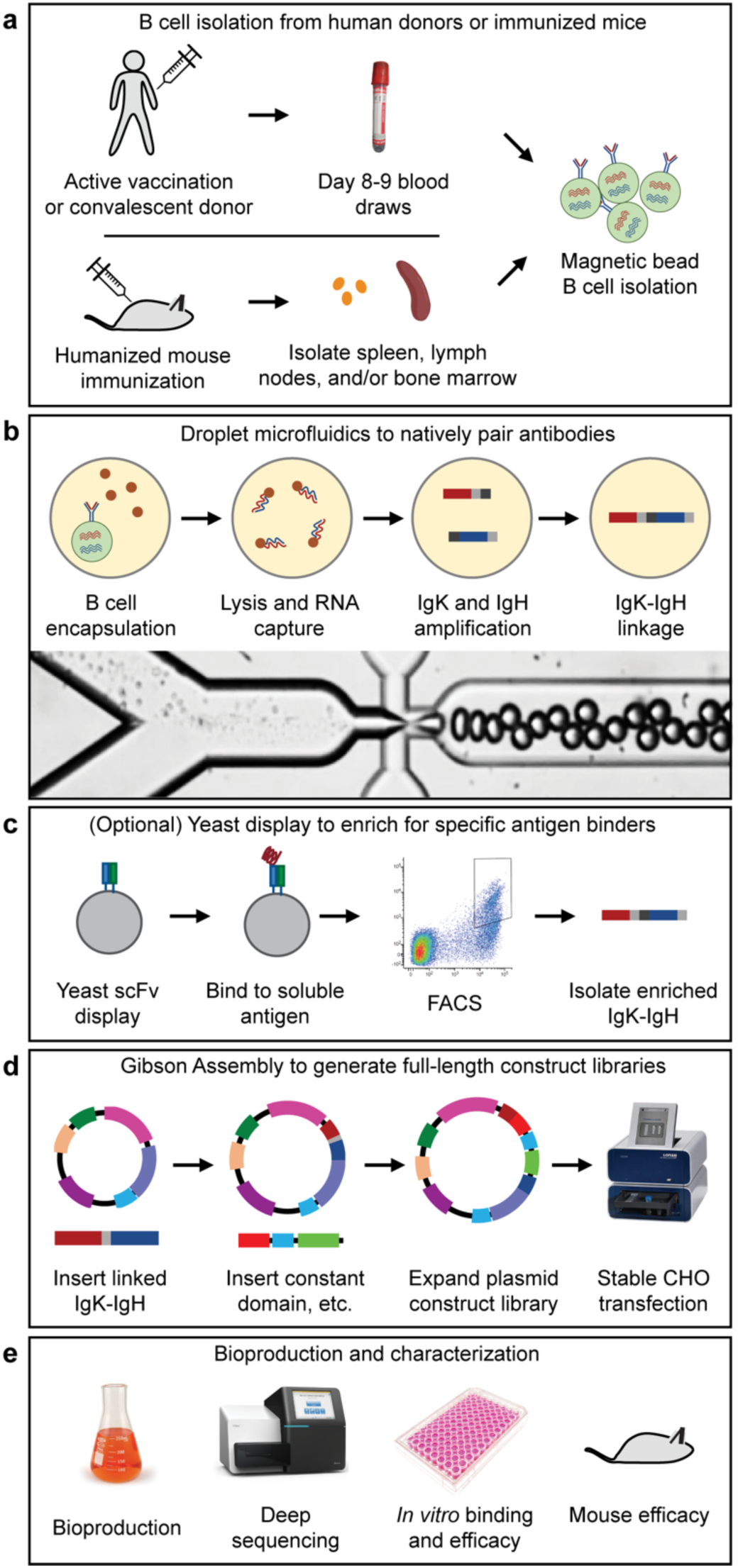
(**a**) B cells were isolated from human donors (vaccinated or convalescent) or immunized humanized mice. (**b**) Droplet microfluidics were used to capture natively paired antibody sequences from millions of single cells. (**c**) An optional yeast scFv display system was used to enrich for binders to a soluble antigen. (**d**) A two-step Gibson Assembly process converted the scFv fragment to full-length antibody expression constructs, which were then stably integrated into CHO cells following electroporation and selection. (**e**) After bioproduction, the libraries were characterized in many ways including deep sequencing, *in vitro* binding and efficacy assays, and *in vivo* mouse efficacy studies.

Next, we used each library of scFv DNAs to produce natively paired full-length antibody expression constructs, which were then engineered into mammalian cells for bioproduction of recombinant hyperimmunes (**Figure 1**). Cloning into full-length antibody expression constructs was performed *en masse*, i.e., we performed all molecular steps on full libraries rather than individual clones. Briefly, the protocol involved a series of two Gibson Assemblies,^20^ which we termed Gibson Assembly 1 (GA1) and Gibson Assembly 2 (GA2) (**Supplementary Figure S1**). In GA1, the scFv library was inserted into a vector backbone that contained a promoter, a fragment of the IgG1 constant domain, and a poly(A) signal. In GA2, we linearized the GA1 plasmid, and subcloned it into a DNA fragment that contained a fragment of the IgK constant domain, a second poly(A) signal, and a second promoter.

Production cell lines for monoclonal antibodies are typically produced by randomly inserting expression constructs into the CHO genome.^16^ This method produces cell lines with genomic insertion of multiple copies of the expression construct. If we randomly inserted our polyclonal antibody construct libraries into the CHO genome, because each cell might contain several inserted transgenes, many clones would express multiple antibodies, which would result in frequent non-native pairing between heavy and light chain Ig. Additionally, different genome locations have different transcriptional activity levels,^21^ which could result in heterogeneous, inconsistent and/or unstable bioproduction. We therefore used CHO cell lines engineered with a Flp recombinase recognition target (FRT) landing pad. We then used these cell lines for stable expression of recombinant hyperimmunes in polyclonal cell banks.

### Recombinant Hyperimmune for Rapid Response to SARS CoV-2

Emerging viruses are a constant and unpredictable threat to human health. In the past two decades alone, the world has seen outbreaks of Ebola virus,^22^ SARS,^23^ Middle East Respiratory Syndrome (MERS),^24^ 2009 H1N1 swine flu,^25^ Zika Virus,^26^ and SARS CoV-2,^27^ among others. Prophylactic vaccines often require long development timelines, for example, the recently approved Ebola virus vaccine required over 15 years of pre-clinical and clinical development before showing efficacy.^28^ Development of broadly neutralizing monoclonal antibodies is confounded by the difficulty of identifying broadly neutralizing epitopes,^22^ which often requires years of research. Because of such issues, convalescent COVID-19 plasma has emerged as a promising approach to address the COVID-19 pandemic.^6–7^ However, convalescent plasma is difficult to manufacture at scale because convalescent plasma supply is constrained and each plasma donor supplies enough therapeutic for only 1-2 patients.

To address the urgent unmet clinical need of the COVID-19 pandemic, we used our technology to build a recombinant hyperimmune against SARS CoV-2, which we called recombinant Coronavirus 2 Immune Globulin, or rCIG. In March 2020, we recruited 50 human donors from a single clinic in Louisiana (USA), who either had tested positive for SARS CoV-2 by nasal swab PCR testing or had shown symptoms of COVID-19 around the time of a major local outbreak. First, we assessed anti-SARS CoV-2 plasma titer for each of the donors using the S1 and receptor binding domain (RBD) regions of SARS CoV-2 Spike glycoprotein (**Figure 2a**; **Supplementary Table S1**). We observed a wide range of EC50s among patients who tested positive for COVID-19 (range: 0.0056-9.94 mg/ml). We selected 16 donors with high plasma antibody titers and used our technology to build yeast scFv display libraries from pools of 2 donors, for a total of 8 libraries. The libraries comprised a median of 70,940 antibodies (range: 54,986-156,592; **Supplementary Table S2**).

**Figure 2.**
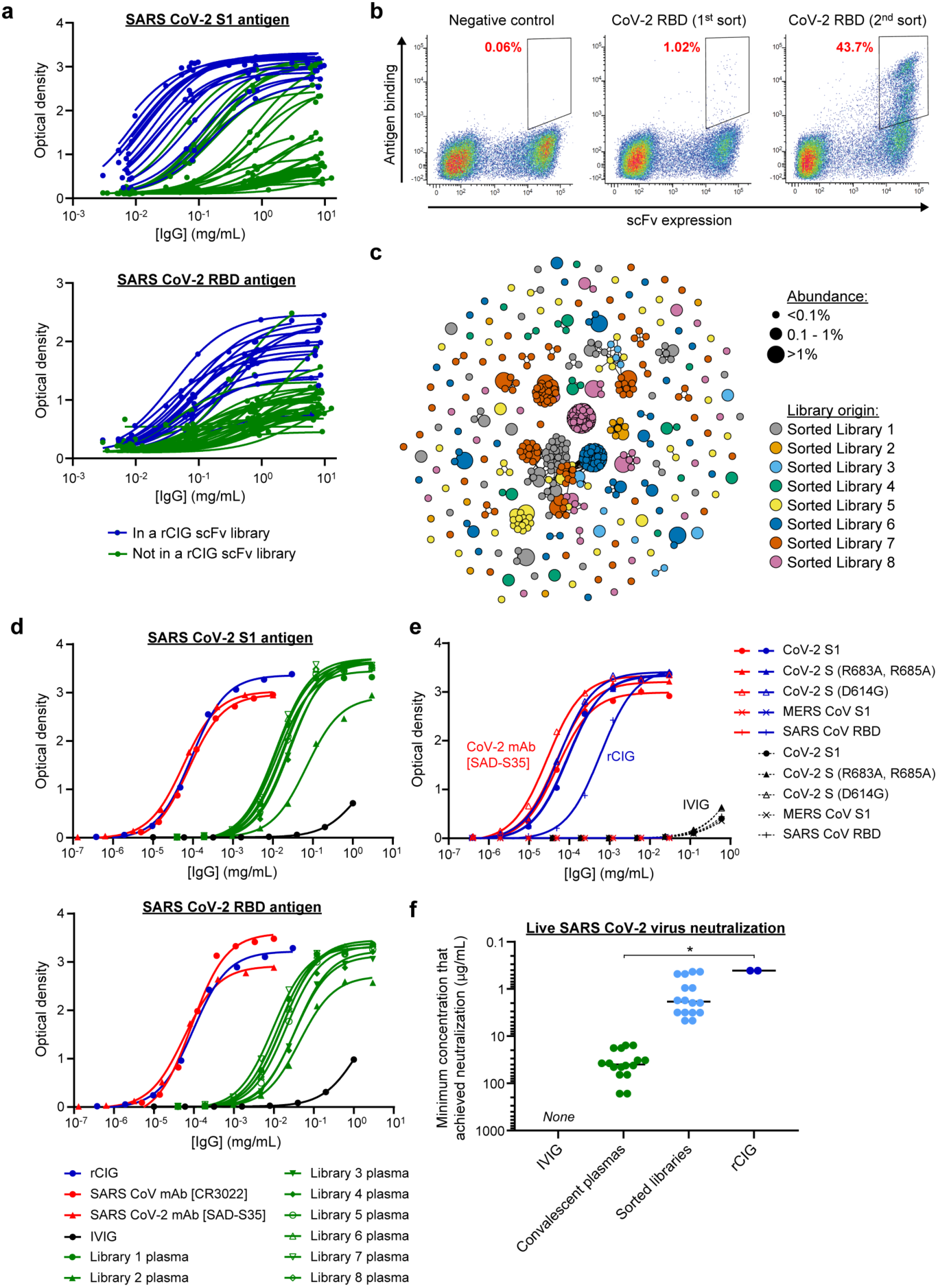
(**a**) ELISA of individual human plasma donors against SARS CoV-2 S1 antigen (top) or RBD antigen (bottom). Dark blue indicates donors used in rCIG. (**b**) Example FACS enrichment of scFv against CoV-2 RBD from library 1 using yeast display. The x-axis measures presence of a C-terminal c-Myc tag, indicating expression of an scFv on the surface of the cell. The y-axis measures binding of antigen to the scFv-expressing cells. The gates used for yeast selection (double positive) are indicated, with the percentage of scFv-expressed antigen binders in red. (**c**) Clonal cluster analysis of rCIG antibodies. Each node represents an antibody clone (full-length heavy chain). The color of the nodes indicates the sorted scFv library from which the scFv clones were derived. The size of the nodes reflects the frequency of the scFv clones in the final library (only clones ≥0.01% are plotted). We computed the total number of amino acid differences between each pairwise alignment, and edges indicate ≤5 amino acid differences. (**d**) ELISA of the indicated samples against SARS CoV-2 S1 antigen (top) or RBD antigen (bottom). (**e**) ELISA of the indicated samples (indicated by the color) against the indicated antigens (different shapes). For rCIG, no binding was observed against MERS CoV S1. For the CoV-2 mAb [SAD-S35], no binding was observed against MERS CoV S1 and SARS CoV RBD. (**f**) Live virus neutralization. Individual dots are separate libraries or samples that represent the minimum antibody concentration that achieved neutralization. Each sample was run in duplicate with the same result observed for each replicate. No neutralization was seen for IVIG. * denotes p<0.05.

We used flow sorting to enrich for anti-SARS CoV-2 antibodies in the 8 yeast scFv libraries (**Figure 2b**; **Supplementary Figure S2**; **Supplementary Table S2**). One round of flow sorting suggested that a median of 0.99% of antibodies (range: 0.42-2.29%) were directed against SARS CoV-2. After two rounds of sorting, a median of 62.7% of unsorted antibody sequences were human IgG1 isotype (range: 51.5-83.4%), whereas in the sorted libraries a median of 82.4% of antibody sequences were human IgG1 isotype (range: 63.6-92.2%), suggesting that the COVID-19 antibody response was generally dominated by IgG1 antibodies. Next, we used our technology to make full-length polyclonal antibody preparations from each of the 8 scFv libraries. We used anti-SARS CoV-2 ELISA, Spike:ACE2 blocking assays, and pseudotype and live virus neutralization assays to assess the relative activity of each of the 8 antibody libraries (**Figure 2f**; **Supplementary Figures S3-S5**; **Supplementary Table S2**). We pooled the 8 scFv-sorted CHO cell banks in a way that sought to balance high antibody diversity with high anti-SARS CoV-2 pseudotype neutralization titer (**Supplementary Table S3**) and used the combined cell bank to generate rCIG protein product (**Supplementary Figure S6**). We completed this entire process, from delivery of the first donor sample to lab-scale generation of the rCIG protein product, in less than three months.

Antibody RNA sequencing of the final CHO cell bank indicated that the rCIG drug candidate comprised a diverse set of 12,500 antibodies (**Figure 2c**; **Supplementary Table S4**). Anti-SARS CoV-2 ELISA suggested that the binding titer of rCIG was between 99- and 747-fold higher than corresponding plasma (**Figure 2d**; **Supplementary Figures S3**; **Supplementary Tables S2, S4**). ELISAs with several natural variants of SARS CoV-2 and antigens from related viruses, including SARS CoV and MERS CoV, showed that rCIG bound a broader variety of antigen targets than IVIG or a neutralizing CoV-2 monoclonal antibody (mAb; **Figure 2e**; **Supplementary Figure S7**; **Supplementary Table S4**). Finally, Spike:ACE2 blocking assays, pseudotype virus neutralization assays, and live SARS CoV-2 neutralization assays suggested that the neutralizing titer of rCIG was between 44- and 1,767-fold higher than corresponding convalescent plasma (**Figure 2f**; **Supplementary Figures S4-S5; Supplementary Tables S2, S4**). We concluded that rCIG was a promising alternative to COVID-19 convalescent plasma due to significantly higher potency against live virus (Wilcoxon rank sum test, p=0.02869) and the ability to scale good manufacturing practice (GMP) production without the need to recruit more donors.

### Recombinant Hyperimmune for Zika Virus

In 2015-2016, a major outbreak of the mosquito-borne flavivirus Zika spread across Latin America.^26^ Though Zika virus has been less widespread and less deadly than SARS CoV-2, Zika can spread from mother to fetus *in utero*, resulting in birth defects such as microcephaly.^29^ As of July 2020, there was no FDA-approved vaccine or therapy for Zika virus. Zika virus disease is complicated by antibody-dependent enhancement (ADE),^30–31^ a phenomenon in which poorly neutralizing antibodies enhance viral infection by bringing virus particles to cells that express Fc receptor (FcR). This problem is particularly troublesome for individuals who have been previously infected with Dengue, a related flavivirus, since many anti-Dengue antibodies are poor neutralizers against Zika virus, and vice versa.^32^ ADE is a safety concern in the development of plasma-derived hyperimmunes and vaccines.^33^

To address the Zika pandemic, we used our technology to build a recombinant hyperimmune against Zika virus, which we termed recombinant Zika Immune Globulin, or rZIG. Though convalescent Zika-infected donors may have been available internationally, we decided to use Zika as a test case to show how a recombinant hyperimmune could be built against an emerging pathogen in the absence of any human donors. Therefore, to create rZIG, we used human-transgenic mice (Trianni) that expressed a complete repertoire of human antibody sequences. The mice were immunized with Zika virus antigens (**Supplementary Figure S8**). To explore our ability to engineer an rZIG that would not exhibit ADE, we additionally boosted with four inactivated Dengue virus serotypes.

We used B cells from the immunized animals and our microfluidics technology to create an scFv library of natively paired IgGs. The resulting scFv library comprised approximately 119,700 IgG-IgK clonotypes (**Supplementary Table S5**). Next, we used the scFv library and our CHO engineering technology to create rZIG CHO cell banks with a wild type human IgG1 isotype (rZIG-IgG1) or a mutated human IgG1 with abrogated FcR binding (rZIG-LALA).^34^ Antibody RNA sequencing of IgG sequences in the rZIG cell banks suggested that the rZIG-IgG1 comprised 33,642 antibodies and rZIG-LALA comprised 26,708 antibodies (**Figure 3a**; **Supplementary Table S6**). A Morisita overlap of 86% and a Jaccard overlap of 58% between the rZIG-IgG1 and rZIG-LALA libraries suggested that the cell banks comprised substantially similar antibody repertoires. We used these CHO cell banks to produce rZIG-IgG1 and rZIG-LALA hyperimmunes at laboratory scale (**Supplementary Figures S9-S10**).

**Figure 3.**
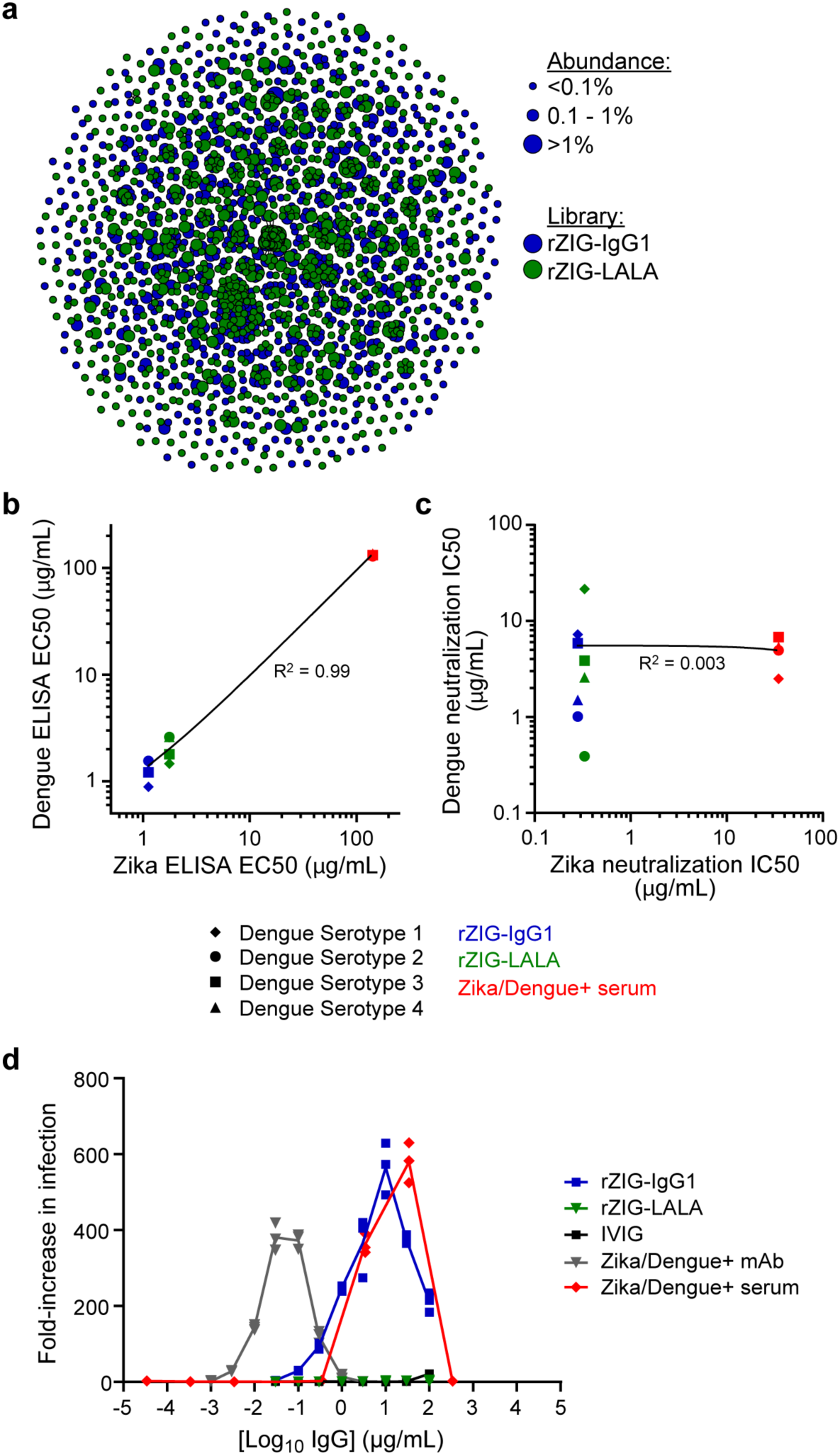
(**a**) Clonal cluster analysis of rZIG-IgG1 (blue) and rZIG-LALA (green) antibodies. Each node represents an antibody clone (full-length heavy chain). The size of the nodes reflects the frequency of the scFv clones in the final library (only clones ≥0.01% are plotted). We computed the total number of amino acid differences between each pairwise alignment after combining both libraries together, and edges indicate ≤5 amino acid differences. (**b**) ELISA of rZIG-IgG1 (blue), rZIG-LALA (green), and Zika/Dengue+ serum control (red) for Dengue serotypes 1-4 (y-axis; indicated by shape) and Zika virus antigen (x-axis). Linear regression trendline is indicated in black. (**c**) Pseudotype neutralization by rZIG-IgG1 (blue), rZIG-LALA (green), and Zika/Dengue+ serum control (red) for Dengue serotypes 1-4 (y-axis; indicated by shape) and Zika virus antigen (x-axis). Linear regression trendline is indicated in black. (**d**) Zika pseudotype virus ADE assay for rZIG-IgG1 (blue), rZIG-LALA (green), and positive and negative controls. Test article concentration is on the x-axis. Fold-increase infection is on the y-axis, which was the infection-induced luciferase signal observed in the presence of antibody divided by the luciferase signal observed with a no antibody control.

Anti-Zika virus ELISA showed that both rZIG-LALA and rZIG-IgG1 had >75-fold higher titers against Zika virus than a human Zika positive serum sample (**Supplementary Figure S11**; **Supplementary Table S6**). Both rZIG-LALA and rZIG-IgG1 had anti-Dengue binding activity across four serotypes, with pooled EC50s showing strong correlation with anti-Zika EC50s (linear regression, R^2^=0.9993, F-statistic p<0.001; **Figure 3b**; **Supplementary Figure S12**; **Supplementary Table S5**). In contrast, though both rZIG-LALA and rZIG-IgG1 had strong activity in a Zika pseudotype neutralization assay (**Supplementary Figure S13**), there was no correlation between Zika and pooled Dengue neutralization (linear regression, R^2^=0.00271, F-statistic p>0.05; **Figure 3c**; **Supplementary Figures S14**). We investigated whether the abrogated Fc function of rZIG-LALA could decrease ADE in a Zika pseudotype virus assay (**Supplementary Figure S15**). Both Zika+ human serum and rZIG-IgG1 showed considerable ADE, whereas rZIG-LALA showed no detectable ADE (**Figure 3d**). We concluded that rZIG-LALA was a promising alternative to plasma-derived drugs, due to high neutralizing potency against Zika virus and complete abrogation of the risk of ADE.

### IVIG Spike-in for Patients with Primary Immune Deficiency

Plasma-derived IVIG acts as antibody replacement for patients with humoral primary immune deficiency (PID), who have low serum IgG titers. Though plasma-derived IVIG reduces rates of serious infections in PID, many patients still suffer frequent serious infections that require hospitalization.^35^ In particular, about 78% of serious lung infections are caused by pneumococcus and Hib bacteria.^36^ Clinicians have improved outcomes by further increasing IVIG doses,^37^ suggesting that plasma-derived IVIG has insufficient anti-pathogen activity for certain at-risk PID patients. However, bacterial species are often incredibly diverse, for example, there are 90 known pneumococcus serotypes,^38^ complicating therapeutic development. To address this unmet clinical need, we manufactured recombinant hyperimmunes directed against pneumococcus and Hib bacteria, designed as polyvalent “spike-ins” for plasma-derived IVIG, i.e., recombinant Haemophilus Immune Globulin (rHIG) and recombinant Pneumococcus Immune Globulin (rPIG).

We recruited healthy human donors and administered vaccines directed against pneumococcus or Hib. Eight to nine days after vaccination, PBMCs were collected and shipped to our microfluidics processing facility. We selected B cells from the PBMCs, ran millions of cells through our microfluidics platform (**Supplementary Table S7**), and then used the scFv libraries and our CHO engineering technology to create IgG1 CHO cell banks for rHIG and rPIG. Heavy chain antibody RNA sequencing of the cell banks indicated that rHIG comprised 49,206 IgG sequences and rPIG comprised 17,938 IgG sequences (**Figure 4a**; **Supplementary Tables S8-S9**). We used these CHO cell banks to produce rHIG and rPIG hyperimmunes at laboratory scale (**Supplementary Figures S16-S17**).

**Figure 4.**
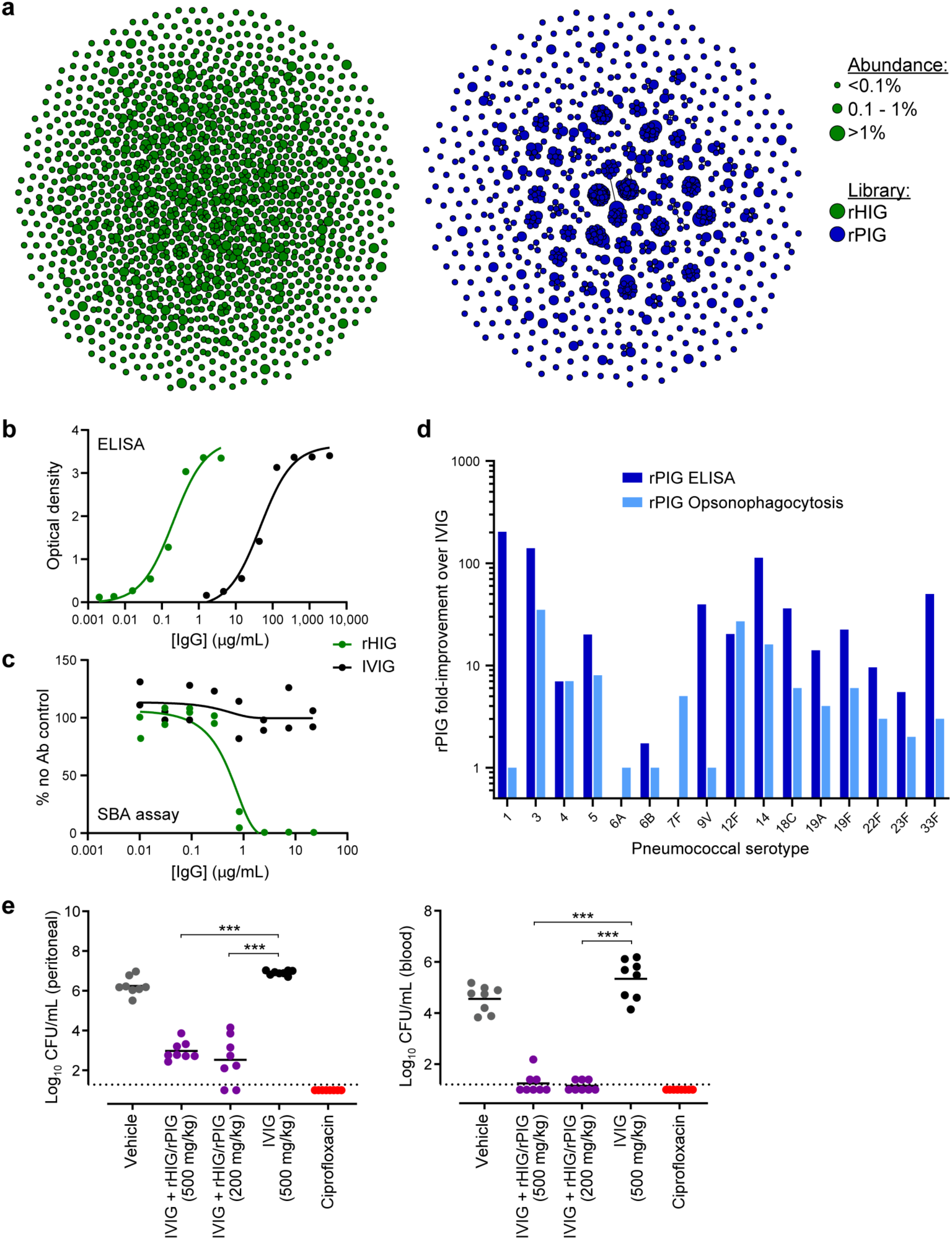
(**a**) Clonal cluster analysis of rHIG (green) and rPIG (blue) antibodies. Each node represents an antibody clone (full-length heavy chain). The size of the nodes reflects the frequency of the scFv clones in the final library (only clones ≥0.01% are plotted). We computed the total number of amino acid differences between each pairwise alignment, and edges indicate ≤5 amino acid differences. (**b**) Anti-Hib ELISA for rHIG (green) and IVIG (black). (**c**) Serum bactericidal assay (SBA) for rHIG (green) and IVIG (black) with the ATCC 10211 Hib strain. % no Ab control (y-axis) was computed as the number of bacterial colonies in the test sample divided by the number of bacterial colonies in a no antibody control sample. (**d**) ELISA binding to (dark blue) or opsonophagocytosis of (light blue) the indicated pneumococcal serotype. Fold-improvement in binding/activity over IVIG was computed as a mean of duplicate measurements for rPIG divided by a mean of duplicate measurements for IVIG (based on the binding concentration for ELISA and the number of bacterial colonies for opsonophagocytosis). (**e**) *In vivo* assay with ATCC 10211 Hib strain. Each circle represents CFU Hib per mL (y-axis) from either peritoneal fluid or blood from a single mouse in a given test group. Black bars represent mean of the CFU Hib per mL. Dotted lines represent the lower limit of detection for CFU quantification. *** denotes p<0.001.

Anti-Hib ELISA indicated that rHIG had 233-fold higher titer than plasma-derived IVIG (**Figure 4b**; **Supplementary Table S8**). A serum bactericidal assay demonstrated that rHIG was strongly active against two different Hib strains, whereas no bactericidal activity was observed for plasma-derived IVIG (**Figure 4c**; **Supplementary Figure S18; Supplementary Table S8**). An ELISA against a combination of 23 pneumococcus serotypes showed that rPIG has 85-fold higher titer than plasma-derived IVIG (**Supplementary Figure S19**; **Supplementary Table S9**). ELISA for individual pneumococcus serotypes showed that rPIG was at least 5-fold higher titer than plasma-derived IVIG for 13 out of 16 serotypes measured, indicating broadly enriched polyvalent reactivity, and significantly higher than IVIG overall across all separate ELISAs combined (Wilcoxon signed rank test, p=0.00123; **Figure 4d**; **Supplementary Table S9**). Finally, semi-quantitative serotype-specific opsonophagocytosis assays suggested that rPIG was as effective or more effective than plasma-derived IVIG at cell killing for 15 out of 16 serotypes tested, and had significantly higher activity than IVIG across all separate opsonophagocytosis assays combined (Wilcoxon signed rank test, p=0.00251; **Figure 4d**; **Supplementary Table S9**).

To simulate the potential clinical application, rHIG and rPIG were mixed in with plasma-derived IVIG (IVIG + rHIG/rPIG) at a ratio of 1:1:8 (rHIG:rPIG:IVIG), producing a product with 18.3-fold higher titer than plasma IVIG for Hib and 8.3-fold higher titer than plasma IVIG for a pool of 23 pneumococcus serotypes (**Supplementary Figure S20**; **Supplementary Table S10**). A Hib mouse challenge model using IVIG + rHIG/rPIG as prophylactic treatment showed significantly lower bacterial loads in the blood (Welch t-test, p<0.001) and peritoneal fluid (Welch t-test, p<0.001) as compared to plasma IVIG alone (**Figure 4e**). We concluded that a polyvalent IVIG + rHIG/rPIG product had strong potential to address unmet clinical needs in PID patients through increased potency against key pathogens.

### Recombinant Human ATG for Transplant Tolerance

In 2019, nearly 40,000 solid organ transplants were performed in the United States alone (www.unos.org). Transplant generally introduces at least some mismatch between the human leukocyte antigen (HLA) genotypes of the donor and host. This frequently results in some host-versus-graft (HVG) effects, leading to loss of the graft and other serious complications. To encourage tolerance of grafts, transplant physicians use a variety immunosuppressive drugs at the time of the graft and thereafter.^39^ One such drug is anti-thymocyte globulin (ATG), which is manufactured by injecting rabbits with human thymocytes and isolating antibodies from the rabbit serum.^40^ However, rabbit Ig can cause allergic reactions and other complications in humans,^11^ and the drug shows significant variation in potency across lots.^15^

To improve on rabbit-ATG, we made a recombinant human ATG, or rhATG, derived from transgenic mice that express human antibodies (Trianni). The mice were immunized with either human T cells or human fetal thymocytes (**Supplementary Figure S21**). We used B cells from the immunized animals and our microfluidics technology to create four scFv libraries of natively paired IgGs: bone marrow cells from T cell immunized mice, lymph node cells from T cell immunized mice, lymph node cells from thymocyte immunized mice, and spleen cells from thymocyte immunized mice. The resulting scFv libraries comprised a range of 13,314 to 34,324 IgG-IgK clonotypes (**Supplementary Table S11**). We then used our CHO engineering technology to make cell banks from each of the four libraries.

We produced protein from each of the CHO cell banks, and then pooled the proteins in equal mass equivalents to create rhATG (**Supplementary Figure S22**). Sequencing of individual libraries suggests that the pool comprised 49,885 antibodies (**Figure 5a**; **Supplementary Table S12**). We then performed ELISA for a panel of known cell surface antigen targets for rabbit-ATG^41^ and observed that rhATG bound several immune cell surface targets, but only a subset of the targets bound by rabbit-ATG (**Supplementary Figure S23**). To investigate further, we performed *in vitro* cell killing assays with human peripheral blood mononuclear cells (PBMCs), and showed that rhATG and rabbit-ATG were not significantly different in cell killing potency against cytotoxic T cells and helper T cells (linear mixed effects model, p>0.05), whereas rhATG is significantly stronger than rabbit-ATG at killing B cells (linear mixed effects model, p<0.01) but significantly weaker than rabbit-ATG at killing NK cells (linear mixed effects model, p<0.01; **Figure 5b**). We also performed anti-erythrocyte binding assays, which suggested that rhATG has less off-target activity than rabbit-ATG (**Supplementary Figure S24**).

**Figure 5.**
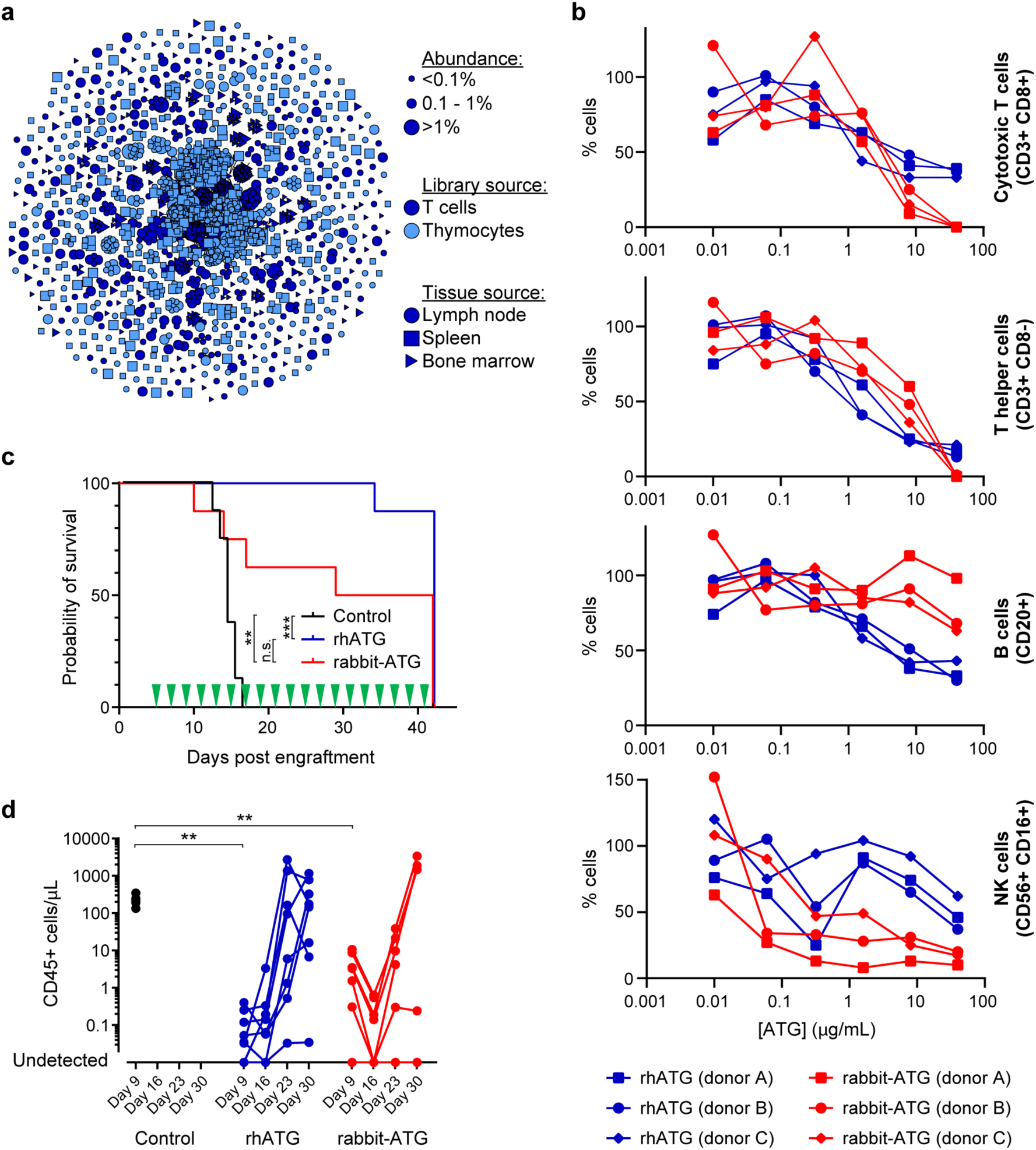
(**a**) Clonal cluster analysis of rhATG antibodies. Each node represents an antibody clone (full-length heavy chain). The color of the nodes indicates the immunized library source. The shape of the nodes indicates the mouse tissue origin. The size of the nodes reflects the frequency of the scFv clones in the final library (only clones ≥0.01% are plotted). We computed the total number of amino acid differences between each pairwise alignment, and edges indicate ≤5 amino acid differences. (**b**) Cell killing assays of a dilution series of rabbit-ATG (red) and rhATG (blue) with three PBMC donors. The y-axis (% cells) was determined by dividing the number of cells of the indicated cell type present after overnight incubation with the indicated amount of antibody by the number of cells of that cell type present in a no antibody control. (**c**) Survival of mice in the GVH study using PBMC donor 1 treated every other day with a negative vehicle control (black), rabbit-ATG (red), or rhATG (blue). Treatment days are indicated by green triangles. ** denotes p<0.01, *** denotes p<0.001, n.s. denotes not significant. (**d**) Flow cytometry was used to determine the concentration of CD45+ cells from each alive mouse on Days 9, 16, 23, and 30 of the GVH study from (c) for negative vehicle control (black circles), rhATG (blue circles), or rabbit-ATG (red circles). Lines connect measurements from each mouse. No CD45+ cells were observed where circles intercept the x-axis. ** denotes p<0.01, *** denotes p<0.001.

We studied the efficacy of rhATG *in vivo*, using a graft-versus-host (GVH) model in which human PBMCs were grafted onto immune-incompetent mice.^42^ We dosed animals (n=8 per PBMC donor) with rhATG, rabbit-ATG, or vehicle control, either every other day for 5 weeks starting 5 days after the PBMC graft, or only on days 5, 6, and 7 after the graft. Two different PBMC donors were tested for each dosing regimen. After 42 days, rhATG was not significantly different than rabbit-ATG for survival (log-rank pairwise tests, p>0.05) and was superior to vehicle control for survival (log-rank pairwise tests, p<0.001), in both dosing schemes across multiple PBMC donors (**Figure 5c**; **Supplementary Figure S25**). In both dosing regimens across both PBMC donors, immune cell (CD45+) expansion was not significantly different between rhATG and rabbit-ATG (linear mixed effects model, p>0.05), whereas for the vehicle control immune cell counts were significantly higher than rhATG at Day 9 (Wilcoxon rank sum tests, p<0.01; **Figure 5d**; **Supplementary Figure S26**). We concluded that though rhATG and rabbit-ATG did not share identical antigen targets, the drugs had similar efficacy *in vivo*, suggesting that rhATG could one day address unmet clinical needs in transplant tolerance.

## DISCUSSION

We have generated the first proofs-of-concept for polyvalent, 10^3^- to 10^4^-diverse recombinant hyperimmune antibody drugs generated from convalescent human blood donors, vaccinated human blood donors, and humanized mouse repertoires. We extensively validated the drug candidates using *in vitro* and *in vivo* methods, highlighting advantages of our drug candidates over plasma-derived incumbents. Our technology for generating the recombinant hyperimmune antibody drugs uniquely combined cutting-edge methods in microfluidics, genomics, and mammalian cell engineering. Contrasted against prior recombinant polyclonal antibody generation methods,^17^ our new class of drugs comprised hundreds-fold higher antibody diversity, and therefore represented substantive fractions of antigen-reactive repertoires. Our recombinant hyperimmunes had significant advantages over plasma-derived products, including higher potency, the ability to scale production without collecting further donors, consistency of production, and the ability to modulate pharmacologic problems such as ADE. The technology is fast, producing a master cell bank against a poorly characterized virus (SARS CoV-2) in less than three months. The rCIG product (GIGA-2050) is currently being manufactured at two good manufacturing practice (GMP) facilities, with the goal of achieving First-in-Human studies in early 2021.

In the future, there are many opportunities to improve our manufacturing processes and further improve upon our drug candidates. First, input linked scFv repertoires typically comprised approximately 2- to 4-fold more antibodies than the final CHO cell banks. Our prior work to clone T cell receptor (TCR) repertoires into Jurkat cells was much more efficient,^43^ but that work used lentivirus rather than Flp-In site directed integration, and therefore many Jurkat clones expressed multiple TCRs. In the future, we will work to improve the efficiency of CHO engineering. Another problem for consideration is that we are currently unable to characterize the protein diversity present in the drug candidates; rather, antibody RNA sequencing is used as a proxy for protein diversity. Advances in proteomics may help to solve this problem.^44^ Our method relies on PCR amplification of RNA from single cells, which may suffer from amplification bias.^45^ Though our goal was not necessarily to precisely recapitulate the input repertoires, PCR bias can result in undesired antibody ratios. Improvements in throughput and cost of DNA synthesis^46^ may abrogate the need for PCR amplification and allow for precise engineering of complex antibody mixtures. Finally, the drug candidates described in this study could be improved in many ways. For example, rPIG did not bind equivalently to different serotypes. To improve this product, we could sort yeast scFv libraries for binders to specific serotypes, and then mix in ratios that might be considered more clinically appropriate. Also, though rhATG functioned similarly to rabbit-ATG in our *in vivo* GVH model, and showed less off-target binding, further *in vivo* work may reveal that a different polyvalent mixture would be more efficacious. Such a mixture could be made by further optimizing mouse immunizations, and/or implementing a yeast scFv sorting protocol that selects for on-target specificity using cell lysate, as we have reported previously.^47^

With this novel technology for recombinant hyperimmunes in hand, we are finally able to consider drug development approaches that routinely combine the advantages of recombinant antibodies (purity, consistency, potency) with the advantages of plasma-derived antibodies (proven efficacy, diversity, polyvalence, *in vivo* affinity maturation). In this current study, we have shown how our technology could be used to improve existing plasma-derived products such as IVIG and rabbit-ATG. In the future, we envision that our technology could be used to develop drugs with novel mechanisms of action, for example, anti-tumor antibody mixtures, or anti-plasma cell mixtures to cure humoral-driven autoimmune disease. Though we demonstrated that our technology could be used to generate a pandemic response very quickly, there needs to be further investment by public health authorities to build out emergency GMP production facilities to quickly generate clinical material during a pandemic. Further, a relatively small budget of <$50 million could be used to pre-emptively generate recombinant hyperimmune cell banks against the 50 most pressing biodefense threats. Thus, when the next pandemic hits, we would be ready with the drug candidates in hand and spare GMP production facilities standing ready – possibly saving hundreds of thousands of lives and trillions of dollars of lost economic activity.

## ACKNOWLEDGEMENTS

This work was partially funded by NIAID grants R44AI115892 and R44AI124901, and NSF grant 1230150, to D.S.J. Cell line CSS-1286 was financed by grant R44AI124901.

The staff at MedPharmics (Metairie, LA, USA) and Access Biologicals (Vista, CA, USA) were instrumental in acquiring convalescent COVID-19 donor samples under IRB-approved protocols. Ms. Jennifer Keller (GigaGen, Inc., South San Francisco, CA, USA) helped to organize collection of COVID-19, Hib, and pneumococcus donor samples. Antibody Solutions (Santa Clara, CA, USA) provided helpful advice in design of mouse immunization work for rZIG and rhATG and performed the immunizations. Jackson Labs (Sacramento, CA, USA) provided useful advice in design of GVH studies and performed the studies. Dr. Erica Stone (GigaGen, Inc., South San Francisco, CA, USA) provided useful assistance in the field of immunology and in critical reviews of the scientific direction. Mr. Carter Keller (GigaGen, Inc., South San Francisco, CA, USA) and Dr. Charles Wilson provided useful strategic advice. The following reagents were obtained through BEI Resources, NIAID, NIH: Vector pCAGGS Containing the SARS-CoV-2, Wuhan-Hu-1 Spike Glycoprotein Gene RBD with C-Terminal Hexa-Histidine Tag, NR-52309; Vector pCAGGS Containing the SARS-Related Coronavirus 2, Wuhan-Hu-1 Spike Glycoprotein Gene (soluble, stabilized), NR-52394 produced under HHSN272201400008C; Spike Glycoprotein Receptor Binding Domain (RBD) from SARS-Related Coronavirus 2, Wuhan-Hu-1, Recombinant from HEK-293T Cells, NR-52306.

## AUTHOR CONTRIBUTIONS

Conceptualization, S.M.K., R.A.M., E.H.M., A.S.A., and D.S.J.; methodology, S.M.K., R.A.M., M.A.A., K.P.C., M.J.S., J.F.S., E.K.W., N.W., C.V.L., S.M.C., B.M., C.O., A.S.A., and D.S.J.; software, R.C.E. and Y.W.L.; investigation, S.M.K., R.A.A., M.S.A., M.A.A., E.B., K.P.C., Y.C., B.K.G., A.G., J.L., R.L., E.P., V.A.M., A.V.M-C., A.R.N., J.S., J.F.S., M.J.S., K.S., B.T., E.K.W., N.W., L.R., H.R., C.V.F.C., T.H.O., and A.T.R.; data curation, S.M.K., R.A.M., M.A.A., Y.W.L., L.T., A.S.A., and D.S.J.; writing—original draft preparation, D.S.J.; writing—review and editing, S.M.K., R.A.M, A.S.A., and D.S.J.; visualization, S.M.K., Y.W.L., A.S.A., and D.S.J.; supervision, M.J.S., A.S.A., and D.S.J.; project administration, S.M.K., R.A.M., C.V.L., D.B., J.V.T., D.G., G.S., M.O.M., H.L., R.J., A.S.A., and D.S.J.; funding acquisition, D.S.J.

## ONLINE METHODS

### Sourcing Human Materials

Local ethical regulations were followed and informed consent was obtained for all human sample collection.

#### rCIG

A contract research organization (CRO; Access Biologics, New Orleans, LA, USA) recruited under sample collection protocol #PRO00026464 (Advarra, Columbia, MD, USA) approved by Institutional Review Board (IRB) and included if donors were 12-46 days (average 24 days +/− 14 days) from the onset of two or more COVID-19 symptoms (fever, cough, shortness of breath, sore throat, and pneumonia). Sixty mL of whole blood was collected in ACD tubes, de-identified, and transported overnight to GigaGen for processing. 16 donors with high SARS CoV-2 Spike antigen-specific antibodies by ELISA (as described below) were included in rCIG and were predominantly Caucasian (87.5%), female (75%), aged 49 (+/− 17 years), collected 21 days (+/− 6 days) from onset of symptoms, and were approximately equally distributed, with 6 donors with 2 symptoms, 6 donors with 3 symptoms, and 4 donors with 4 symptoms.

#### rHIG

A contract research organization (BloodCenter Wisconsin, Milwaukee, WI, USA) vaccinated two donors (Donor 1, a 26-year-old Caucasian female, and Donor 2, a 21-year-old Asian male) with PedvaxHIB vaccine (Merck, Kenilworth, NJ, USA). Leukapheresis was performed eight or nine days later to obtain PBMCs. In parallel, plasma was isolated from separate blood draws on the day of leukapheresis and prior to vaccination. We performed ELISA against Hib (Alpha Diagnostics, San Antonio, TX, USA; see methods below) on the plasma samples to confirm a response to the vaccine as compared to plasma from the same donors prior to vaccination. Sample collection protocols were approved by IRB protocol #PRO00028063 (Medical College of Wisconsin/Froedtert Hospital IRB) to GigaGen. Informed consent was obtained from all participants and samples were shipped to GigaGen de-identified.

#### rPIG

A contract research organization (AllCells, Alameda, CA, USA) vaccinated three donors (Donor 1, 57-year-old Caucasian male; Donor 2, 44-year-old Caucasian male; Donor 3, 35-year-old Caucasian/Asian male) with Pneumovax®23 vaccine (Merck, Kenilworth, NJ, USA). We performed a 60 mL blood draw eight days later. Plasma and pan-B cells were isolated from whole blood (see methods below). We performed ELISA against a mixture of all 23 pneumococcal polysaccharides (Alpha Diagnostics, San Antonio, TX, USA; see methods below) on the plasma samples to confirm response to the vaccine. Sample collection protocols were approved by IRB protocol #7000-SOP-045 (Alpha IRB, San Clemente, CA, USA) to AllCells. Informed consent was obtained from all participants and samples were shipped to GigaGen de-identified.

### Processing Human Materials

For whole blood, PBMCs and plasma were isolated using density gradient centrifugation SepMate tubes with Lymphoprep medium (StemCell Technologies, Vancouver, BC, Canada). To isolate pan-B cells from PBMCs (from either whole blood or a leukopak), we used the Human EasySep Pan-B Cell Enrichment Kit (StemCell, Vancouver, BC, Canada). After isolation, the cells were cryopreserved using CryoStor® CS10 (StemCell Technologies, Vancouver, BC, Canada). Immediately prior to generating paired heavy and light chain libraries, cells were thawed, washed in cold DPBS+0.5% BSA, assessed for viability with Trypan blue on a disposable hemocytometer (Bulldog Bio, Portsmouth, NH, USA) or with AOPI on a Cellometer K2 (Nexcelom Bioscience, Lawrence, MA, USA), and then re-suspended in 12% OptiPrep™ Density Gradient Medium (Sigma, St. Louis, MO, USA) at 5,000-10,000 cells per μl. This cell mixture was used for microfluidic encapslation as described below.

### Immunization of Trianni Mouse® Mice

Humanized mice were obtained from Trianni (San Francisco, CA, USA). Local ethical regulations were followed for mouse immunizations, by Antibody Solutions IACUC (Santa Clara, CA, USA).

#### rZIG

Two Trianni humanized mice were immunized consecutively weekly with Zika VLP, inactivated Dengue 1, inactivated Dengue 4, inactivated Dengue 3, then inactivated Dengue 2 with alhydrogel/muramyl dipeptide (ALD/MDP) adjuvant. Animals were checked for antibody titer and boosted with Zika VLPs without adjuvant 5 days before harvest (Antibody Solutions, Santa Clara, CA, USA).

#### rhATG

Two Trianni humanized mice were immunized weekly with human thymocytes from 5 de-identified specimens acquired from a CRO (Vitalant Research Institute, San Francisco, CA, USA) for 5 weeks with ALD/MDP adjuvant and boosted on week 6 without adjuvant. Three Trianni mice were immunized weekly for 5 weeks with Pan T cells (StemCell, Vancouver, Canada) in ALD/MDP isolated from PBMCs from 1 de-identified donor (StemCell), checked for an elevated antigen-specific antibody titer, and boosted with the same cells 5 days before harvest without adjuvant (Antibody Solutions, Santa Clara, CA, USA).

After sacrifice, spleen, lymph nodes, and/or bone marrow were harvested and processed into a single cell suspension. Samples from multiple mice were pooled together by tissue and pan-B cells were isolated from spleen and lymph node tissue using the EasySep Mouse Pan-B Cell Enrichment Kit (StemCell Technologies, Vancouver, BC, Canada). CD138+ cells were isolated from bone marrow using Miltenyi CD138+ mouse microbeads (Miltenyi, Bergisch Gladbach, Germany). After isolation, the cells were cryopreserved using CryoStor® CS10 (StemCell Technologies, Vancouver, BC, Canada).

### Generating Paired Heavy and Light Chain Libraries

Generation of scFv libraries from antibody-producing cells^18^ comprises three steps: (i) poly(A)+ mRNA capture, (ii) multiplexed overlap extension reverse transcriptase polymerase chain reaction (OE-RT-PCR), and (iii) nested PCR. Briefly, a microfluidic device captures single cells in droplets with a mixture of lysis buffer and oligo dT beads (NEB, Ipswich, MA, USA). After the cell is lysed and mRNA is bound to the bead, the emulsion is broken, and the mRNA-containing beads are purified. Next, an emulsion is created using OE-RT-PCR reagents and the beads as template. The emulsion is subjected to thermal cycling which creates cDNA, amplifies the IgK and IgG variable regions, and links them together in an scFv format. Then the emulsion is broken and the linked scFv DNA product is extracted and purified. The purified scFv product is then amplified using nested PCR to remove artifacts and add adapter sequences. Depending on the adapter sequences, the product can be used for deep sequencing, yeast display libraries, or full-length CHO expression.

To convert the scFv libraries into full-length CHO expression libraries, we first used nested outer PCR primers to add adapters with overhangs for Gibson assembly to the 5’ and 3’ ends of the scFv library (for rCIG, this was done after yeast scFv display enrichment, as described in the next section). Then NEBuilder HiFi DNA Assembly Master Mix (NEB, Ipswich, MA, USA) was used to insert the scFv library into a vector containing a single promoter, a secretory leader sequence for light chain Ig and the remainder of the IgG1 constant region, creating a cloned scFv library. This intermediate library was transformed into *E. coli* and plasmids were purified by either (a) spreading onto LB-ampicillin plates, scraping 0.5-1 million colonies and pooling or (b) inoculating directly into LB-ampicillin broth and growing overnight. Plasmid purification was performed using ZymoPURE II Plasmid Maxiprep Kits (Zymo Research, Irvine, CA, USA). To create the full-length antibody library, we performed a second Gibson assembly by linearizing the product of GA1 with BamHI-HF (rHIG) or NheI-HF (rCIG, rPIG, rhATG, and rZIG) (NEB, Ipswich, MA, USA) and using it as a vector to insert a synthetic amplicon containing a portion of the light chain Ig constant region, a poly(A) signal for light chain Ig, a promoter for the IgG gene and a secretory leader sequence for the IgG gene. The full-length library was then transformed into *E. coli* and spread on LB-ampicillin plates. We typically combined >0.5 million colonies and purified plasmid with a ZymoPURE II Plasmid Maxiprep Kits (Zymo Research, Irvine, CA, USA) to make the full-length recombinant hyperimmune maxiprep library for transfection. When the transformed *E. coli* were inoculated directly into LB-ampicillin broth, a small volume of cells was plated to calculate the total number of transformants. In some cases, ampicillin was used for both plates and broth, whereas in other cases carbenicillin was used instead of ampicillin. Paired heavy and light chain libraries were made only once from each sample.

### Enrichment for Antigen Binders by Yeast scFv Display

Polyclonal COVID-19 scFv libraries were sorted^18^ to enrich for relevant sequences. Briefly, yeast surface display scFv libraries were generated using COVID-19 scFv DNA libraries and a custom yeast surface display vector transformed by electroporation into EBY100 yeast strain (MYA-4941; ATCC, Manassas, VA, USA). Surface displayed scFv sequences include a C-terminal myc tag to identify scFv expression with an anti-myc primary (A21281; Thermo Fisher Scientific, Waltham, MA, USA) and AF488 secondary antibody (A11039; Thermo Fisher Scientific, Waltham, MA, USA). Binding to antigen was identified by staining with soluble biotinylated SARS CoV-2 receptor binding domain antigen (SPD-C82E9; Acro Biosystems, Newark, DE, USA) at 1200 nM and APC-streptavidin (SA1005; Thermo Fisher Scientific, Waltham, MA, USA). Stained yeast libraries were sorted on a FACSMelody (BD Biosciences, San Jose, CA, USA) and double positive (AF488+/APC+) cells were collected. The gating strategy is outlined in **Supplementary Figure S2**. The collected cells were expanded and sorted again to further enrich the libraries. After the second round of sorting, cells were expanded a third time prior to plasmid isolation with a Zymoprep Yeast Plasmid Miniprep kit (Zymo Research, Irvine, CA, USA). The plasmid libraries were then used as template for barcoding PCR and subsequent analysis by deep sequencing (Illumina, San Diego, CA, USA). Plasmid from twice-sorted libraries was used as template for PCR towards full-length CHO antibody expression. Yeast scFv sorting was performed only once from each yeast scFv library.

### Cell Line Used for rHIG and rhATG

We adapted the adherent Flp-In™-CHO cell line with a genetically integrated FRT site (Thermo Fisher Scientific, Waltham, MA, USA) to suspension culture. For all steps in the adaptation process, “Ham’s F-12” refers to Ham’s F-12 (with L-glutamine, Thermo Fisher Scientific, Waltham, MA, USA) plus 10% FBS (Thermo Fisher Scientific, Waltham, MA, USA), and “BalanCD” refers to BalanCD CHO Growth A (Irvine Scientific, Santa Ana, CA, USA) with 4 mM Glutamax (Thermo Fisher Scientific, Waltham, MA, USA). To adapt this cell line to suspension, we first passaged the cells into a mixture of 50% Ham’s F-12 plus 50% BalanCD in T-flasks. Cells were next passaged into 25% Ham’s F-12 plus 75% BalanCD and switched to shaking Erlenmeyer flasks. Cells were then passaged into 10% Ham’s F-12, 90% BalanCD + 0.2% anti-clumping agent (Irvine Scientific, Santa Ana, CA, USA) and banked for future use.

Approximately 100 million of the adapted Flp-In CHO cells were transfected per recombinant hyperimmune library using an Amaxa Nucleofector 4D (SG buffer, pulse DU133; Lonza, Basel, Switzerland). These cells were plated into shaking Erlenmeyer flasks and recovered in an incubator at 37°C, 5% CO_2_, 125 rpm. After 48 hours, the cells were counted to determine viability, cells were seeded at 1 million cells/mL, and selection was started using 600 μg/mL Hygromycin-B (Gemini Bio, West Sacramento, CA, USA) in fresh media. Cells were counted and media was changed every 2-3 days during the 7-day selection. The libraries were kept on 600 μg/mL Hygromycin-B (Gemini Bio, West Sacramento, CA, USA) during expansion until viability exceeded 95%. When cells were >95% viable and doubling every 24 hours, the adapted Flp-In™-CHO cell line was banked for liquid nitrogen storage. Before banking, cells were sampled from each library, RNA was purified, and antibody RNA-seq (Illumina, San Diego, CA, USA) was performed to assess the diversity of the libraries (**Supplementary Tables S8, S12**).

### Cell Line Used for rPIG, rZIG, and rCIG

A landing pad construct (PMD-4681) was designed and cloned at GigaGen. PMD-4681 was based on pFRT-lacZeo (Thermo Fisher Scientific, Waltham, MA, USA), with some modifications. In place of the LacZ expression construct a cassette was inserted coding for expression of CD34 and GFP. The CD34, GFP, and downstream Zeocin resistance genes (ZeoR present in pFRT-lacZeo) were separated by 2A motifs (T2A or P2A) to allow for translation of three separate polypeptide chains. The CD34 sequence was sourced as a gBlock from IDT (Coralville, IA, USA). The GFP sequence was sourced from ATUM (DasherGFP; Newark, CA, USA).

The GMP suspension CHO line CHOZN® GS-/- was obtained from MilliporeSigma (St. Louis, MO, USA). PMD-4681 was linearized using ScaI-HF and purified via ethanol precipitation. Cells were transfected with the linearized DNA using Amaxa Nucleofector 4D, SE kit, pulse CM-150 (Lonza, Basel, Switzerland). Cells recovered overnight in an incubator and were plated the next day into minipools at approximately 5,000 cells per well, across ten 96-well plates in selective media. The remaining cells were plated and selected together as a bulk pool control. Wells were topped off with fresh media every seven days until at least 80% confluency was reached.

A total of 236 minipools grew out and were screened in parallel for high GFP expression via flow cytometry and low copy number with a quantitative PCR Copy Number Variation (CNV) assay. Minipools with a copy number less than 2.5 and GFP expression at least 50% of the bulk pool were expanded into shaking adaptation. Expanded pools were re-tested for GFP expression via flow cytometry.

Cells were then adapted to BalanCD CHO Growth A in preparation for plating into semi-solid media. Minipools were deemed fully adapted when cells showed consistent doubling times and high viability (>90%). Adapted cells were plated into semisolid media for the Molecular Devices (Fremont, CA, USA) ClonePix3 single cell cloning platform. Single cell imaging was obtained on day 0 of cell plating in semisolid media to confirm monoclonality. After 14 days, clonal cell colonies were picked and deposited as one colony per well of a 96-well plate. Each clone was then expanded, re-adapted to selection media, and cryopreserved. Doubling times were calculated and clones with less than a 30-hour doubling time were chosen for further development. Expanded clones were retested for GFP expression and copy number.

Remaining clones were transfected in duplicate using the Gene Pulser Xcell Total System (BioRad, Hercules, CA, USA) per guidelines from MilliporeSigma (St. Louis, MO, USA) for use with CHOZN GS with a mAb-cyan fluorescent protein (FrostyCFP, ATUM, Newark, CA, USA) construct to test expression titer. CFP expression was evaluated via flow cytometry 3 days post transfection to confirm transfection efficiency >35%. After full selection and recovery, cell lines were tested in a 10-day fed batch TPP shaking production run in duplicate. Titers for candidate cell lines ranged from 50–100 mg/L. A single clone (CSS-1286) was selected to use for recombinant hyperimmune expression.

For transfection of recombinant hyperimmunes into CSS-1286, approximately 50 million cells were transfected per recombinant hyperimmune library using the BioRad Gene Pulser Xcell Total System (Hercules, CA, USA), per guidelines from MilliporeSigma (St. Louis, MO, USA) for use with CHOZN GS. The cells were plated into T-75 flasks (approximately 10 million cells per flask) and recovered in an incubator at 37°C, 5% CO_2_ for 72 hours. After 72 hours, the cells were counted to determine viability and then seeded into 100 mL fresh media without glutamine (EX-CELL CD CHO Fusion, MilliporeSigma, St. Louis, MO, USA) in a 500 mL Erlenmeyer flask. Cells were counted and media was changed every 2-3 days during the ∼14-day selection. When cells were >95% viable and doubling every 24 hours, the cell line was banked for liquid nitrogen storage. Before banking, cells were sampled from each library, RNA was purified, and antibody RNA-seq (Illumina, San Diego, CA, USA) was performed to assess the diversity of the libraries (**Supplementary Tables S4, S6, S9**).

### Bioproduction of rHIG and rhATG

Adapted Flp-In™-CHO cells stably expressing antibody libraries were grown in media consisting of 90% BalanCD CHO Growth A Medium (Irvine Scientific, Santa Ana, CA, USA), 9% Ham’s F-12 (Thermo Fisher Scientific, Waltham, MA, USA), 1% FBS (Thermo Fisher Scientific, Waltham, MA, USA), 4 mM Glutamax (Thermo Fisher Scientific, Waltham, MA, USA), 0.2% anti-clumping agent (Irvine Scientific, Santa Ana, CA, USA), 600 μg/mL Hygromycin-B (Gemini Bio, West Sacramento, CA, USA). Protein production was performed at either small (250 mL) or medium (5 L) scale. For small-scale production, cells were seeded at 1*×*10^6^ cells/mL into 50 mL media in a 250 mL Erlenmeyer flask and grown at 37°C, 5% CO_2_, 125 rpm. Cells were continually grown under these conditions and supplemented with 7.5 mL CHO Feed 1 (Irvine Scientific, Santa Ana, CA, USA) on Days 2, 4 and 7 of the production run. Supernatant was harvested on Day 8 or 9 by centrifugation followed by filtration through a 0.22 μm 250 mL filter bottle (MilliporeSigma, St. Louis, MO, USA) with 1 μm pre-filter (MilliporeSigma, St. Louis, MO, USA). Harvested cell culture fluid (HCCF) was stored at 4°C (if less than 1 week) or at −80°C (if more than one week) until Protein A purification. For medium-scale production, cells were grown in the same media. Cells were then seeded at 1*×*10^6^ cells/mL in 2.3 L in a 5 L flask (in duplicate; Day 0). Each flask was fed with 345 mL CHO Feed 1 (Irvine Scientific, Santa Ana, CA, USA) on Days 2 and 4 of the culture. Cultures were harvested on Day 8 or 9. Each of the four rhATG protein libraries were produced separately.

### Bioproduction of rPIG, rZIG, and rCIG

CSS-1286 CHO cells stably expressing antibody libraries were grown in media without glutamine (EX-CELL CHOZN Advanced; MilliporeSigma, St. Louis, MO, USA). Protein production was performed at either small (500 mL flask) or medium (5 L flask) scale. For small-scale production, cells were seeded at 0.5*×*10^6^ cells/mL into 100 mL media in a 500 mL Erlenmeyer flask and grown at 37°C, 5% CO_2_, 125 rpm. Cells were continually grown under these conditions and supplemented with 15 mL CHO Feed 1 (MilliporeSigma, St. Louis, MO, USA) on Day 3, and 10 mL CHO Feed 1 (MilliporeSigma, St. Louis, MO, USA) on Days 6 and 8 of the production run. Starting on Day 3, glucose was measured each day and supplemented to 6 g/L if below 4 g/L. Supernatant was harvested after cell viability peak and before dropping below 70% viability between Days 9-11, centrifuged and filtered through a 0.22 μm 250 mL filter bottle (MilliporeSigma, St. Louis, MO, USA) with 1 μm pre-filter (MilliporeSigma, St. Louis, MO, USA). HCCF was stored at 4°C (if less than 1 week) or at −80°C (if more than one week) until Protein A purification. For medium-scale production, cells were grown in the same media. Cells were seeded at 0.5*×*10^6^ cells/mL in 2.2 L in a 5 L flask (in duplicate; Day 0). Each flask was fed with 330 mL CHO Feed 1 (MilliporeSigma, St. Louis, MO, USA) on Day 3 and 220 mL CHO Feed 1 (MilliporeSigma, St. Louis, MO, USA) on Days 6 and 8 of the production run. Starting on Day 3, glucose was measured each day and supplemented to 6 g/L if below 4 g/L. Cultures were harvested on Day 10-12.

### Protein Production and Characterization

After harvest, HCCF was purified using MabSelect PrismA (Cytiva, Marlborough, MA, USA) using 1*×* PBS (Teknova, Hollister, CA, USA) for running and wash buffer, 0.1 M Citrate pH 3.0 (Teknova, Hollister, CA, USA) for elution, and 1 M Sodium Citrate pH 6.0 (Teknova, Hollister, CA, USA) for neutralization. The protocol was 10CV of equilibration, HCCF loading, 10CV of washing, and 5-10CV of elution followed by cleaning-in-place with 1M NaOH. HCCF was loaded with a 1 minute residence time. Eluted material was neutralized to a pH of ∼4.5 and centrifuged to remove any precipitation. This material was dialyzed into 0.2 M glycine, pH 4.5 (Teknova, Hollister, CA, USA) using a 20K MWCO Dialysis Cassette (Thermo Fisher Scientific, Waltham, MA, USA) and optionally concentrated up to 30mg/mL using a 50kDa MWCO spin device (MilliporeSigma, St. Louis, MO, USA). Final material was sterilized with a 0.22 μm filter and quantified by A280 (NanoDrop; Thermo Fisher Scientific, Waltham, MA, USA). For rhATG, each of the four libraries were purified by Protein A separately and then equally pooled based on mass.

Purity of the protein was determined by SEC-HPLC. 20 μg of material at 1 mg/mL was injected over a 300 Å, 2.7 μm, 7.8*×*300 mm size exclusion column (Agilent, Santa Clara, CA, USA) using a mobile phase of 25 mM phosphate, 200 mM NaCl pH 7.0 with 10% acetonitrile at 1 mL/min. The percent monomer was determined by integrating the product peaks and reporting the percent area corresponding to ∼150 kDa. The product was further characterized by running 2 μg on a 12% SDS-PAGE gel under reduced and non-reduced buffering conditions and imaged after staining with SimplyBlue SafeStain (Thermo Fisher Scientific, Waltham, MA, USA).

### Deep Antibody Repertoire Sequencing

Deep antibody sequencing libraries were prepared as described previously,^18^ quantified using a KAPA quantitative PCR Illumina Library Quantification Kit (Roche, Mannheim, Germany), and diluted to 17.5 pM. Libraries were sequenced on a MiSeq (Illumina, San Diego, CA, USA) using a 500 cycle MiSeq Reagent Kit v2, according to the manufacturer’s instructions. To make sequencing libraries, we used tailed-end PCR to add Illumina sequencing adapters to the 5’ and 3’ ends of the constructs of interest. For scFv libraries (after droplet emulsion breaking or yeast plasmid isolation), a forward read of 340 cycles was used to capture the light chain CDR3 sequence, and a reverse read of 162 cycles was used to capture the linked heavy chain CDR3 sequence. For CHO libraries, the full-length heavy chain sequence was obtained using overlapping forward and reverse reads of 251 cycles. To determine the Fc isotype of a library, the heavy chain was amplified with a primer that binds further into the constant domain to add the Illumina sequencing adapter to the 3’ end; the first 60 bp of the constant domain was sequenced to determine the isotype, which was linked to the corresponding CDR3H that was simultaneously sequenced. Each library was sequenced one time.

Sequence analysis, including error correction, reading frame identification, and FR/CDR junction calls was performed using our previously reported bioinformatics pipeline.^18^ Reads with E > 1 (E is the expected number of errors) were discarded, such that we retained sequences for which the most probable number of base call errors is zero. Clones are defined as sequences with unique CDR3 amino acid sequences (CDR3K + CDR3H for scFv clones, CDR3H only for CHO clones). For the clonal cluster analysis, we used USEARCH^48^ to compute the total amino acid differences between each pairwise alignment of heavy chain sequences with abundance ≥0.01% in each CHO cell bank. We then used the R package igraph^49^ (version 1.2.4.1) to generate clustering plots for the pairwise alignments. The sequences were represented as “nodes”, with the color (and sometimes shape) defined in the respective figures. The size of the nodes reflects the frequency of the clone (small, <0.1%; medium, 0.1-1%; large, >1%). “Edges” are the links between nodes, which indicate pairwise alignments with ≤5 amino acid differences. The layout_with_graphopt (niter = 3000, charge = 0.03) option was used to format the output. To assess antibody repertoire overlap between libraries, we computed Jaccard and Morisita indices using the R package tcR (version 2.3.2).^50^

Sequencing data are available in the Short Read Archive (SRA) under project identifier PRJNA649279.

### *In vitro* Efficacy Studies

#### rCIG

Anti-SARS CoV-2 antibody reactivities were measured using a protocol based on published ELISA methods.^51^ In brief, SARS CoV-2 Spike and RBD (wild type and variant proteins; Sino Biological, Wayne, PA, USA) were used to coat ELISA plates at 2 μg/mL. Serial dilutions of antibody preparations including test plasma and recombinant products, positive control monoclonal antibodies (CR3022; Absolute Antibody, San Diego, CA, USA, and SAD-S35; Acro Biosystems, Newark, DE, USA) and negative control IVIG (Gamunex; Grifols, S.A., Sant Cugat, Spain) were performed in dilution buffer (1*×* PBS + 0.05% Tween + 0.3% dry milk) in singlet. Quantitative measurements were performed on a plate reader (Molecular Devices, Fremont, CA, USA) and analyzed using Softmax Pro (Version 7.1; Molecular Devices, Fremont, CA, USA) to calculate the EC50 concentrations of samples. The concentration of total IgG was calculated by Cedex Bioanalyzer Human IgG assay (Roche, Mannheim, Germany).

Blocking of binding between Spike RBD and ACE2 was demonstrated by ELISA (BPS Bioscience, San Diego, CA, USA). In brief, SARS CoV-2 Spike RBD protein was coated onto an ELISA plate, serial dilutions of test plasma and recombinant products, positive control monoclonal antibodies (CR3022; Absolute Antibody, San Diego, CA, USA, and SAD-S35; Acro Biosystems, Newark, DE, USA) and negative control IVIG (Gamunex; Grifols, S.A., Sant Cugat, Spain) were performed in singlet in dilution buffer (1*×* PBS + 0.05% Tween + 0.3% dry milk). After incubation, ACE2-His was added at 2.5 ng/mL. After further incubation, anti-His-horseradish peroxidase was added. The plate was developed for a chemiluminescent readout. Quantitative measurements were performed on a plate reader (Molecular Devices, Fremont, CA, USA) and analyzed using Softmax Pro (Version 7.1; Molecular Devices, Fremont, CA, USA) to calculate the EC50 concentrations of samples.

The SARS CoV-2 pseudotype virus neutralization assay was performed in a 96-well plate using ACE2 expressing HEK-293T target cells (CRL-11268; ATCC, Manassas, VA, USA) transiently transfected with TMPRSS-2 expression plasmid. The GFP reporter pseudotype virus expressing SARS-CoV-2 spike (Integral Molecular, Philadelphia, PA, USA) was mixed with test plasma, test rCIG, positive control monoclonal antibodies (CR3022; Absolute Antibody, San Diego, CA, USA, and SAD-S35; Acro Biosystems, Newark, DE, USA) and negative control IVIG (Gamunex; Grifols, S.A., Sant Cugat, Spain) at a five-fold dilution series in singlet. After one-hour incubation, 4*×*10^4^ cells target cells were added to each well and incubated at 37°C for 48 hours. After incubation, the media was removed from all wells without disturbing the adherent cells. TrypLE (Thermo Fisher Scientific, Waltham, MA, USA) was added to each well and incubated for 3 minutes at 37°C. Media was added to stop trypsinization and cells were stained with DAPI and passed through a 30-40 μm filter (Pall Corporation, Port Washington, NY, USA) before quantifying GFP+ cells using a Cytoflex LX (Beckman Coulter, Indianapolis, IN, USA). Flow cytometry data were analyzed by FlowJo (BD Biosciences, San Jose, CA, USA).

SARS CoV-2 microneutralization assays were performed at the Regional Biocontainment Laboratory at Duke University Medical Center (Durham, NC, USA) in a 96-well plate format using Vero E6 cells (CRL-1586; ATCC, Manassas, VA, USA) infected with 100 TCID_50_ dose of the 2019-nCoV/USA-WA1/2020 strain. Test and control samples were initially diluted to 1:50, then a 12-step, two-fold serial dilution of test antibodies was performed before infection of the cells; every test or control was run in duplicate. IVIG (Gamunex; Grifols, S.A., Sant Cugat, Spain) was used as negative control. Cell-only control wells were included alongside virus-only treated wells. Following 4 days of infection, culture media was removed, and cell monolayer was fixed with 10% neutral buffered formalin (NBF) and stained with 0.1% Crystal Violet. Absorbance at 590 nm or visual inspection was used to measure the monolayer condition/level of infection. Neutralization was reported as the lowest concentration of sample that prevents cytopathic effect in the monolayer of cells.

#### rZIG

Zika- and Dengue-specific antibodies were measured by ELISA. A 96-well microtiter plate was coated with either 2 µg/ml Zika or Dengue Serotype 1, 2, 3, or 4 recombinant envelope proteins (ProSpec Bio, East Brunswick, NJ, USA) in 1*×* carbonate coating buffer (BioLegend, San Diego, CA, USA) and incubated overnight at 4°C. After blocking the coated plate with ultrablock buffer (Bio-Rad, Hercules, CA, USA) and washing with PBS + 0.05% Tween-20 (Teknova, Hollister, CA, USA), eight-step three-fold serial dilutions in assay buffer (1*×* PBS + 0.05% Tween + 0.3% dry milk) were performed on rZIG-IgG1, rZIG-LALA, Zika/Dengue+ serum positive control (Seracare, Milford, MA, USA), and a negative control IVIG (Gamunex; Grifols, S.A., Sant Cugat, Spain). Dilutions were added in duplicate and incubated at 37°C for 1 hour. Next, a secondary rabbit anti-human IgG horseradish peroxidase conjugate (Southern Biotech, Birmingham, AL, USA) was added, and the plate was washed and developed using 3,3’,5,5’-tetramethylbenzidine (TMB) substrate solution (Thermo Fisher Scientific, Waltham, MA, USA). The reaction was halted after 8 minutes using sulfuric acid stopping solution (Southern Biotech, Birmingham, AL, USA). Quantitative absorbance measurements were performed on a SpectraMax i3x plate reader (Molecular Devices, Fremont, CA, USA) at 450 nm and 620nm. Standard curves (OD_450-620_) were artificially set to max out at 2.97 absorbance value. EC50 values were calculated by non-linear regression analysis using GraphPad Prism (San Diego, CA, USA).

Zika and Dengue *in vitro* pseudotype neutralization assays were performed at Vitalant Research Institute (VRI, San Francisco, CA, USA). rZIG, a Zika/Dengue-specific immune sera (UWIS; de-identified sample screened positive for Zika and Dengue 1-4 by University of the West Indies), monoclonal antibody positive control (UWI-mAb1; IgG1 isotype cloned from de-identified donor by University of the West Indies, found to be cross-reactive to Zika and Dengue 1-4), and IVIG negative control (Gamunex; Grifols, S.A., Sant Cugat, Spain) were co-incubated with reporter virus particles (RVPs; Integral Molecular, Philadelphia, PA, USA) expressing both luciferase and flavivirus-specific glycoproteins as previously described.^52^ Briefly, BHK/DC-SIGN cells (CRL-325; ATCC, Manassas, VA, USA) were seeded in black 96-well plates and then incubated with a 7-step, 3-fold serial dilution of antibodies pre-incubated for one hour at 37°C with RVPs and tested in duplicate. After 72 hours cells were lysed and luciferase activity measured using lysis buffer and firefly luciferase substrate following manufacturer’s guidelines (Promega, Madison, WI, USA). Infection-induced relative light units (RLU) in the presence of test articles were calculated as the RLU of the test article divided by the RLU of a no-serum control infection. The amount of protein required to inhibit 50% of the maximum untreated Zika or Dengue RLUs (IC50) was calculated by non-linear regression analysis using GraphPad Prism (San Diego, CA, USA).

Zika pseudotype assays for *in vitro* antibody-dependent enhancement (ADE) were performed at VRI. rZIG, the Zika/Dengue-specific immune sera UWIS, the monoclonal antibody positive control UWI-mAb1, and IVIG negative control (Gamunex; Grifols, S.A., Sant Cugat, Spain) were serially diluted and co-incubated with Zika pseudotype RVPs at 37°C for 1 hour before addition to K562 chronic myelogenous leukemia cells (CCL-243; Manassas, VA, USA) in U-bottom 96-well plates in triplicate. After a 72 hour incubation at 37°C, cells were harvested, lysed, and infection-induced relative light units (RLU) in the presence of test articles were calculated as the RLU of the test article divided by the RLU of a no antibody control infection (to determine the reported fold-increase in infection).

#### rHIG

The Human Anti-Hib-PRP IgG ELISA kit (#980-100-PHG, Alpha Diagnostics, San Antonio, TX, USA) was used for anti-Hib ELISA titers. Serial dilutions of test articles were performed in Low NSB (non-specific binding) sample diluent in singlet. IVIG (Gamunex; Grifols, S.A., Sant Cugat, Spain) was used as a reference control. Quantitative measurements were performed on a plate reader (Molecular Devices, Fremont, CA, USA) at 450 nm. EC50 values were calculated using SoftMax Pro (Molecular Devices, Fremont, CA, USA).

*In vitro* serum bactericidal assay neutralization studies for Hib were performed at ImQuest (Frederick, MD, USA). The *Haemophilus influenzae* strain ATCC 10211 was obtained from ATCC (Manassas, VA, USA) as a lyophilized stock and was propagated as recommended by the supplier. The Eagan strain was obtained from Zeptometrix (Buffalo, NY, USA). Colonies from an overnight incubation on chocolate agar plates were inoculated into growth media (Brain Heart Infusion, or BHI broth; BD Biosciences, San Jose, CA, USA, with 2% Fildes enrichment; Remel, San Diego, CA, USA) and allowed to achieve an optical density of 625 nm (OD_625_) of approximately 0.4. The culture was adjusted to an OD_625_ of 0.15, which is equivalent to approximately 5*×*10^8^ colony forming units (CFU)/mL. The culture was further diluted to 5*×*10^4^ CFU/mL in dilution buffer (Hanks Balanced Salt Solution; Gibco, Waltham, MA, USA, with 2% Fildes enrichment; Remel, San Diego, CA, USA). The density of the bacterial culture used in the assay was confirmed by plating 50 μL of the 5*×*10^3^ and 5*×*10^2^ dilutions in duplicate on chocolate agar and enumerating the colonies following incubation at 37°C/5% CO_2_ for 24 hours. rHIG and IVIG (Gamunex; Grifols, S.A., Sant Cugat, Spain) reference control were diluted three-fold in buffer, starting at 200 μg/mL such that a total of ten total dilutions were evaluated in singlet. 10 μL of each dilution of test article was added in duplicate to a 96-well microtiter plate. ATCC 10211 bacteria at a concentration of approximately 5*×*10^4^ CFU/mL were then added to the plate in a volume of 20 μL, such that the total in-well bacterial density would be 1*×*10^4^ CFU/20 μL. Following an incubation of 15 minutes at 37°C/5% CO_2_, 25 μL of baby rabbit complement (Pel-Freez; Rogers, AR, USA) and 25 μL of dilution buffer was added to each well. The plate was incubated at 37°C/5% CO_2_ for 60 minutes. Following the incubation, 5 μL of each reaction mixture was diluted in 45 μL of dilution buffer and the entire 50 μL was plated on chocolate agar plates. The plates were incubated for approximately 16 hours at 37°C/5% CO_2_. Following incubation, bacterial colonies were enumerated. The fold-dilution of the test article that killed >50% of the bacteria is the serum bactericidal index (SBI).

#### rPIG

The Human Anti-S. Pneumococcal vaccine (Pneumovax/CPS23) IgG ELISA kit (Alpha Diagnostics #560-190-23G, San Antonio, TX, USA) was used in parallel with the human anti-S. pneumoniae CWPS/22F IgG ELISA kit (#560-410-C22, Alpha Diagnostics, San Antonio, TX, USA) for initial assessment of anti-pneumococcal titers against a pool of all 23 polysaccharides included in the vaccine.^18^

Serotype-specific antibodies were measured by ELISA and opsonophagocytosis. The concentrations of serotype-specific IgG antibody were calculated using the standard reference serum, lot 007SP (National Institute for Biological Standards and Control; Hertfordshire, UK), using the standardized pneumococcal reference ELISA as previously described.^53^ Briefly, 96-well flat-bottomed microtiter plates were coated with capsular polysaccharide antigens (LGC Standards, Teddington, UK) from pneumococcal serotypes 1, 3, 4, 5, 6A, 6B, 7F, 9V, 12F, 14, 18C, 19A, 19F, 22F, 23F, and 33F. All samples were tested in duplicate and double absorbed with CWPS and with purified serotype 22F polysaccharide to neutralize the anti-cell wall polysaccharide and nonspecific homologous antibodies to serotype 22F, except for the 22F assay which was absorbed with CWPS Multi, as described in the WHO reference ELISA protocol.^54^ Plates were washed, and a titration of rPIG and reference control IVIG (Gamunex; Grifols, S.A., Sant Cugat, Spain) was performed. Plates were incubated and washed again, and prediluted alkaline phosphatase-conjugated goat anti-human IgG (Thermo Fisher Scientific, Waltham, MA, USA) was added to each well. After another incubation, the plates were washed a final time and *p*-nitrophenyl phosphate substrate (MilliporeSigma, St. Louis, MO, USA) was added. Following a final incubation, the reaction was stopped by adding 3 M NaOH (Thermo Fisher Scientific, Waltham, MA, USA) to each well. Plates were read using a microtiter plate reader (SPECTROstar Omega; BMG Labtech, Buckinghamshire, UK) at 405 and 620 nm.

The opsonophagocytic indices (OI) to the same pneumococcal serotypes were evaluated by multiplexed opsonophagocytic assay, as previously described.^55^ In brief, frozen aliquots of target pneumococci were thawed, washed twice with opsonization buffer B (HBSS with Ca and Mg, 0.1% gelatin, and 10% fetal bovine serum), and diluted to the proper bacterial density (approximately 2*×*10^5^ CFUs/mL each serotype). Equal volumes of four bacterial suspensions chosen for simultaneous analysis were pooled. Duplicate serially diluted test articles (20 μL/well) were mixed with 10 μL of bacterial suspension in each well of a microplate. After 30 minutes of incubation at room temperature with shaking at 700 rpm, 10 μL of 3- to 4-week-old rabbit complement (Pel-Freeze, Rogers, AR, USA) and 40 μL of differentiated HL60 cells (10^7^ cells; CCL-240, ATCC, Manassas, VA, USA) were added. Plates were incubated in a 37°C/5% CO_2_ incubator with shaking at 700 rpm. After being incubated for 45 minutes, plates were placed on ice for 20 minutes, and an aliquot of the final reaction mixture (10 μL) was spotted onto four different Todd-Hewitt broth with 0.5% yeast extract and 0.75% agar (THY) plate. When the fluid was absorbed into the agar, an equal volume of overlay agar containing one of four antibiotics (optochin, spectinomycin, streptomycin, or trimethoprim) was applied to each THY agar plate. After overnight incubation at 37°C, the number of bacterial colonies in the agar plates was enumerated. IVIG was used as reference control (Gamunex; Grifols, S.A., Sant Cugat, Spain). The OI was defined as the test product dilution that kills 50% of bacteria and was determined by linear interpolation.

#### rhATG

To assess relative amount and specificity of rhATG, we performed an ELISA on antigens known to be expressed on thymocytes and previously described as having rabbit-ATG reactivity.^41^ Rabbit-ATG positive control was from Sanofi Genzyme (Thymogobulin; Cambridge, MA, USA). T cell antigens (CD3, CD4, CD5, CD7, CD8, CD16a, CD32a, CD45, CD81, CD85 CD95) were purchased from Sino Biological (Wayne, PA, USA) and individually coated onto 96-well ELISA plates (Thermo Fisher Scientific, Waltham, MA, USA) at 1μg/mL in 1*×* carbonate coating buffer (BioLegend, San Diego, CA, USA). After an overnight incubation at 4°C, coated plates were washed and blocked (Bio-Rad, Hercules, CA, USA) for 1 hour. Polyclonal products were diluted to 200 μg/mL of total IgG and an 8-step 1:3 titration in assay buffer (1*×* PBS + 0.05% Tween + 0.3% dry milk) was performed. The antibody titrations were added to each antigen and incubated for 1 hour at 37°C. Polyclonal HRP goat anti-rabbit IgG (E28002; Novodiax, Hayward, CA, USA) or mouse anti-human IgG HRP (109-035-088; Jackson ImmunoResearch, West Grove, PA, USA) were diluted 1:2500 and incubated on the plate for 1 hour. Plates were washed, developed using 1-step ultra TMB substrate (Thermo Fisher Scientific, Waltham, MA, USA), and stopped with 1 N HCl. Plates were read by a spectrophotometer (Molecular Devices, Fremont, CA, USA) at 450 nm and analyzed with Softmax Pro (v7; Molecular Devices, Fremont, CA, USA).

To assess off-target antibody binding we performed a red blood cell antigen binding assay, using the Capture-R kit (Immucor, Norcross, GA, USA). Using a dilution of test article (Thymoglobulin or rhATG) or positive control from the Immucor kit, samples were added to the plate and incubated for 1hr at 37°C in singlets. ELISA plates were washed and incubated Polyclonal HRP goat anti-rabbit IgG (E28002; Novodiax, Hayward, CA, USA) or mouse anti-human IgG HRP (109-035-088; Jackson ImmunoResearch, West Grove, PA, USA). Subsequently, plates were washed and developed with ultra-TMB substrate (Thermo Fisher Scientific, Waltham, MA) and the reaction was stopped with 3 M NaOH (Thermo Fisher Scientific, Waltham, MA, USA) and read the plate on a spectrophotometer at 450 nm.

To determine *in vitro* function of rhATG, peripheral blood mononuclear cells were isolated from whole blood acquired from a CRO (StemCell Technologies, Vancouver, Canada) and frozen. PBMCs were thawed, washed, and plated at 1.5*×*10^5^ cells/well in singlets. Thymoglobulin or rhATG were diluted five-fold starting at 40 μg/mL total IgG and co-incubated with each donor PBMC. Cells were co-incubated overnight at 37°C. After incubation, cells were washed, FcR blocked, and stained for CD45 (clone H130; BioLegend, San Diego, CA, USA), CD3 (clone UCHT1; BioLegend, San Diego, CA, USA), CD8 (clone BW135/80; Miltenyi, Bergisch Gladbach, Germany), CD20 (clone 2H7; BioLegend, San Diego, CA), CD56 (clone 5.1H11; BioLegend, San Diego, CA, USA), and CD16 (clone 3G8; BioLegend, San Diego, CA, USA). Flow cytometry was performed using a Cytoflex LX (Beckman Coulter, Indianapolis, IN, USA) and a consistent collection volume of 150 seconds per well was implemented for every sample. The data were analyzed by FlowJo (BD Biosciences, San Jose, CA, USA). Cell counts after antibody co-incubation relative to no-antibody control (% cells) were calculated. Results were graphed in GraphPad Prism (San Diego, CA, USA). The gating strategy is outlined in **Supplementary Figure S27a**.

### *In Vivo* Mouse Efficacy Studies

Ethical approval was obtained by Institutional Animal Care and Use Committees (IACUCs) at either SSI (Copenhagen, Denmark) for the Hib challenge model or Jackson Laboratory (Sacramento, CA, USA) for the GVH model.

#### IVIG + rHIG/rPIG

For IVIG + rHIG/rPIG *in vivo* challenge studies, the *Haemophilus influenza* strain ATCC 10211 was grown on chocolate agar plates overnight at 35°C and 5% CO_2_. Single overnight colonies were resuspended in sterile saline to 1.5*×*10^8^ CFU/mL. This suspension was diluted in BHI broth to 1.5*×*10^7^ CFU/mL and further diluted in BHI broth with 5% mucin and 2% hemoglobin to 1.5*×*10^4^ CFU/mL. In an IACUC-approved protocol (SSI, Copenhagen, Denmark), Balb/cJ mice (Taconic, Rensselaer, NY, USA; n=6 per group) were inoculated with single 0.5 mL intraperitoneal doses of 10^5^ CFU/mL ATCC 10211. Approximately 1 hour before inoculation, mice were treated orally with 45 μL Nurofen (30 mg/kg) as pain relief. Twenty-four hours prior to Hib inoculation, mice were intravenously administered 200 mg/kg IVIG + rHIG/rPIG mixture, 500 mg/kg IVIG + rHIG/rPIG mixture, 500 mg/kg IVIG (Gamunex; Grifols, S.A., Sant Cugat, Spain), or saline (no treatment). For the ciprofloxacin positive control, one hour after Hib inoculation, mice were dosed with 20 mg/kg ciprofloxacin. Mice were scored for clinical signs of infection, then after 6 hours all animals were sacrificed and blood and peritoneal fluid was collected for CFU determination by serial dilution and plating of 0.02 ml spots on chocolate agar plates.

#### rhATG

We contracted with Jackson Laboratory (Sacramento, CA, USA) under an IACUC-approved protocol to Jackson Laboratory to test rhATG for ability to delay GVHD in immunodeficient NOD *scid* gamma (NSG) mice (genotype: NOD.*Cg-Prkdc^scid^ Il2rg*^tm1Wjl^/SzJ), compared against rabbit-ATG (Thymoglobulin; Sanofi Genzyme, Cambridge, MA) and a vehicle control. Each animal was grafted with approximately 1*×*10^7^ PBMC of a single human donor. On Day 5 after PBMC engraftment, animals were randomized by weight and dosed intravenously every other day for two weeks with 5.5 mg/kg rhATG (n=8), 6.5 mg/kg Thymoglobulin (n=8), or a vehicle control (n=8), or Days 5, 6, and 7 post-engraftment with 5.5 mg/kg rhATG (n=8), 6.5 mg/kg Thymoglobulin (n=8), or a vehicle control (n=8). Two PBMC donors were tested for each dosing regimen (2 PBMC donors *×* 2 dosing regimens *×* 3 treatment groups *×* 8 animals per group = 96 animals). Animals were assessed for clinical signs of mortality daily. Mice were euthanized by CO_2_ asphyxiation before final study take down if they showed >20% weight loss from their starting weight or a combination of the following clinical signs: >10-20% weight loss from their starting weight, cold to touch, lethargic, pale, hunched posture and scruffy coat. 50 μL of blood was drawn from all alive animals on Days 9,16, 23, and 30 post-engraftment via retro-orbital bleed, and flow cytometry stained for Human (hu)CD45-PE (clone HI30; BioLegend, San Diego, CA, USA) and 7AAD (BioLegend, San Diego, CA, USA); 50 μL of CountBright beads (Thermo Fisher Scientific, Waltham, MA, USA) were added to each sample prior to acquisition. Flow cytometric data acquisition was performed using the BD Biosciences FACSCanto flow cytometer (San Jose, CA, USA), and data were acquired and analyzed using BD Biosciences FACSDiva™ software (version 8.0 or higher; San Jose, CA, USA); lymphocytes, singlet, live cells, and CD45+ cells were gated and cell numbers quantified relative to CountBright beads. The gating strategy is outlined in **Supplementary Figure S27b**.

### Statistical Analysis

All statistical tests were performed on non-normalized data, two-sided without adjustments to type I error rates. We used a significance threshold of *α*=0.05 for all statistical tests. All statistical analyses were conducted using R version 3.6.2.

#### Figure 2f

A Wilcoxon rank sum test was used to compare the minimum concentration to achieve SARS CoV-2 live virus neutralization between convalescent plasma samples and rCIG. Duplicate live virus neutralization measurements were pooled.

#### Figure 3b

Simple linear regression was used to calculate the coefficient of determination (R^2^) between Zika and Dengue ELISA EC50 values. EC50 values for all Dengue serotypes were pooled for the analysis. Significance of the regression model was determined using an F-statistic with 1 and 10 degrees of freedom.

#### Figure 3c

Simple linear regression was used to calculate the coefficient of determination (R^2^) between Zika and Dengue pseudotype neutralization IC50 values. IC50 values for all Dengue serotypes were pooled for the analysis. Significance of the regression model was determined using an F-statistic with 1 and 10 degrees of freedom.

#### Figure 4d

Fold improvement over IVIG, by assay (ELISA or opsonophagocytosis) was tested using a one-sample Wilcoxon signed rank test, with the null hypothesis that the median equals 1, i.e., H_0_ =1. For each assay, all individual serotypes were pooled a single Wilcoxon signed rank test. Values for each individual serotype were generated by dividing the mean of duplicate rPIG measurements by the mean of duplicate IVIG measurements.

#### Figure 4e

Welch’s t-tests were used to compare CFU Hib per mL between test groups. Degrees of freedom were 7.87 for IVIG + rHIG/rPIG (500 mg/kg) and 7.13 for IVIG + rHIG/rPIG (200 mg/kg) in peritoneal fluid. Degrees of freedom were 10.87 for IVIG + rHIG/rPIG (500 mg/kg) and 8.03 for IVIG + rHIG/rPIG (200 mg/kg) in blood.

#### Figure 5b

Linear mixed effects models were used to compute p-values for each of the four cell types, with group and concentration as fixed effects and PBMC donor as a random effect to account for the dependence of repeated measures. Degrees of freedom were 31 for each of the four models.

#### Figure 5c, Supplementary Figure S25

Kaplan-Meier survival models were fit on time to mortality and pairwise log rank tests were performed to compare median survival between treatment groups.

#### Figure 5d, Supplementary Figure S26

Linear mixed effects models were used to compute p-values for trends in CD45+ cell counts in each of the four GVH experiments (2 PBMC donors *×* 2 drug dosing regimens = 4 experiments) with day as a fixed effect and PBMC donor as a random effect to account for the dependence of repeated measures. A Wilcoxon rank sum test was used to compare CD45+ cell counts on Day 9 for saline negative control vs. rhATG and saline negative control vs. rabbit-ATG, in each of the four GVH experiments (2 PBMC donors *×* 2 drug dosing regimens = 4 experiments).

---

**Supplementary Figure S1.**
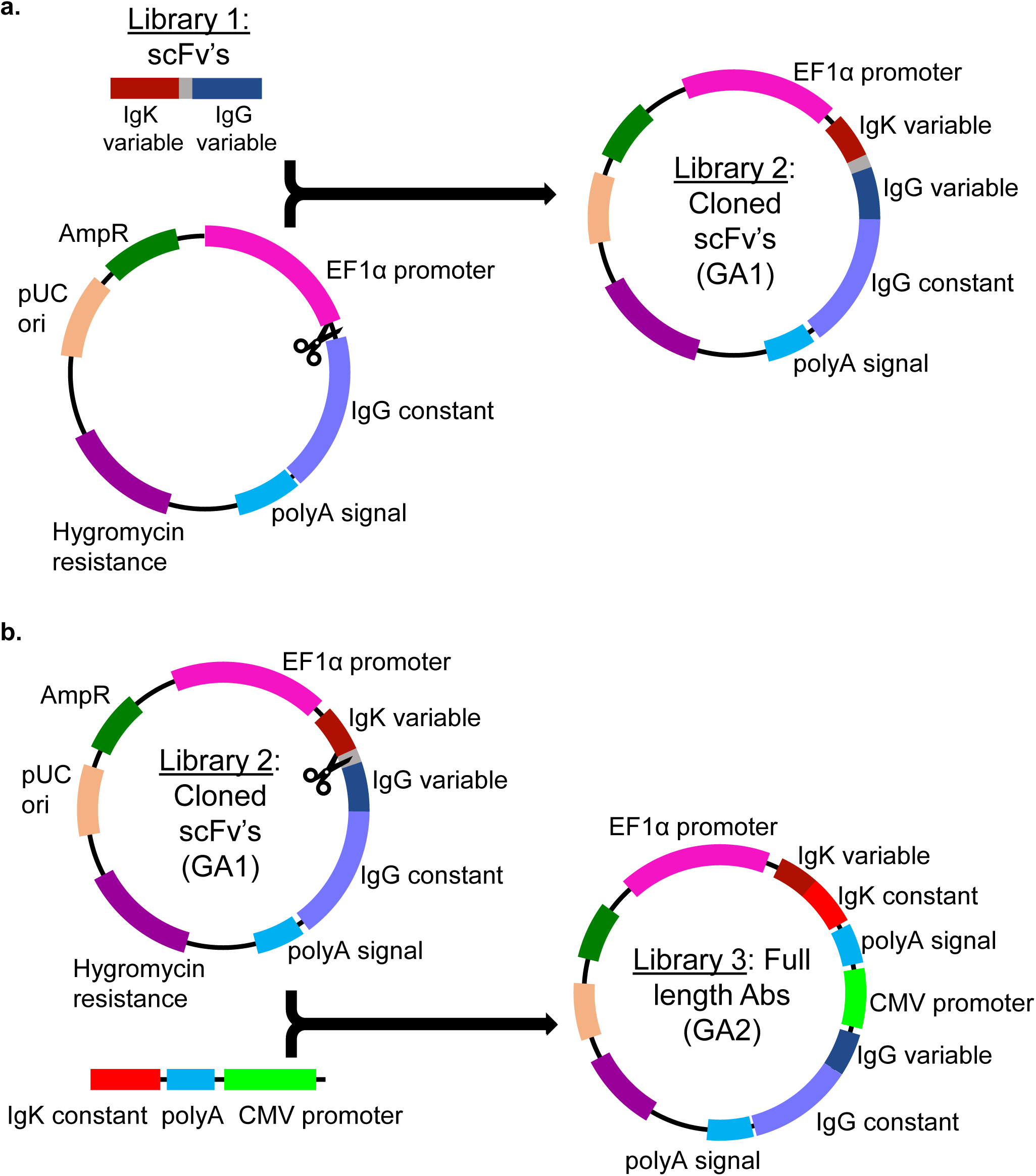
Schematic of Gibson assembly processes. (**a**) Product of Gibson Assembly 1 (GA1), comprising the based plasmid vector with an scFv (from Library 1) cloned between the EF1α promoter and the IgG constant domain sequence. (**b**) Product of Gibson Assembly 2 (GA2), with the scFv linker excised and replaced with a sequence that comprises an IgK constant domain sequence, a poly(A) signal sequence, and a CMV promoter.

**Supplementary Figure S2.**
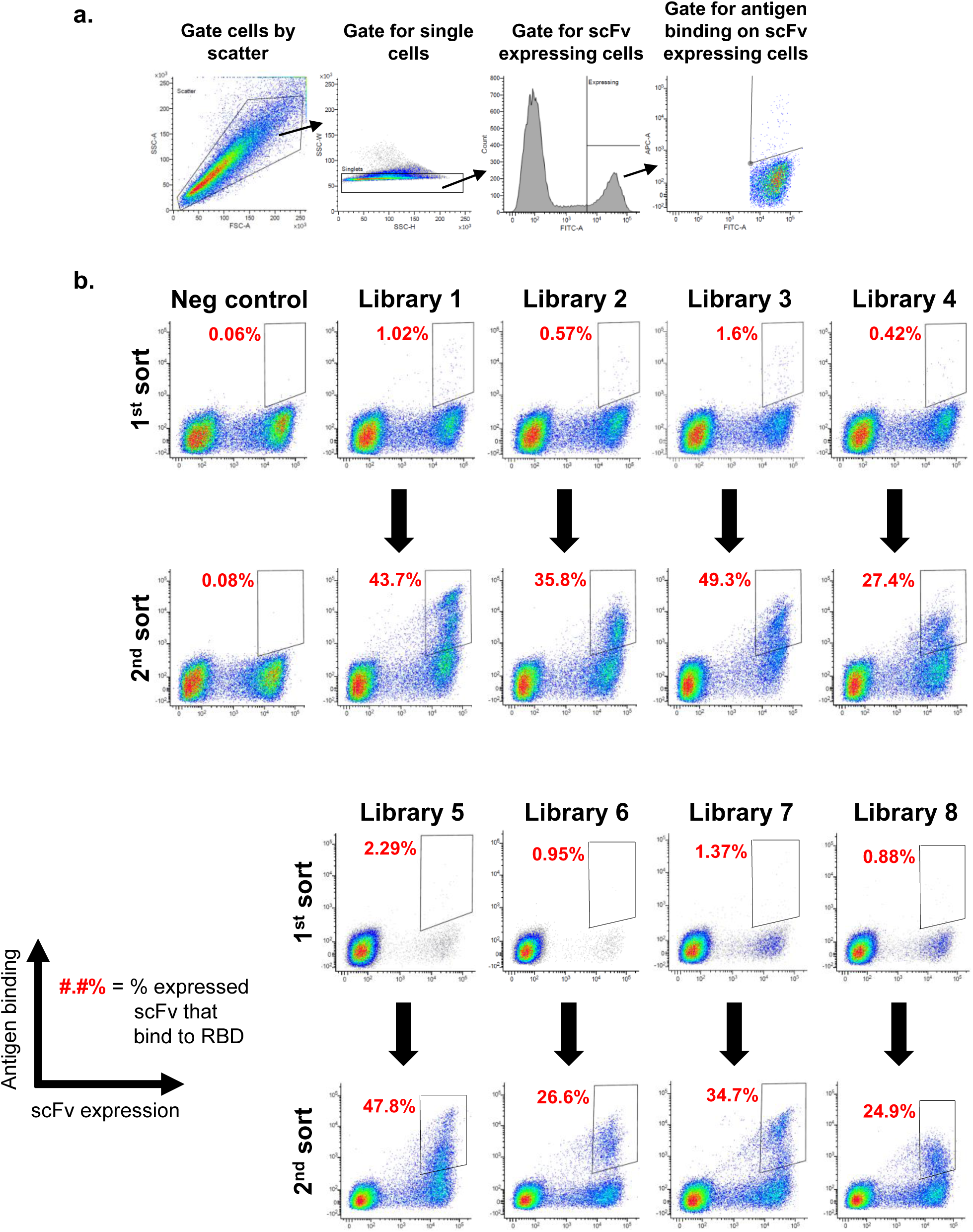
Flow sorting for SARS-CoV-2 specific antibodies using yeast scFv display. (**a**) Gating strategy used for each yeast sort. (**b**) The x-axis measures presence of a C-terminal c-Myc tag, indicating expression of an scFv on the surface of the cell. The y-axis measures binding of antigen to the scFv-expressing cells. The gates used for yeast selection (double positive) are indicated, with the percentage of scFv-expressed RBD binders in red. An irrelevant negative control and the eight rCIG convalescent donor libraries were stained with 1200 nM biotinylated CoV-2 RBD-His. An average of 1.1% of the expressed antibodies were RBD-specific on the first sort. After the second sort, the RBD-specific scFv were amplified and then cloned into full-length antibody expression plasmids. The data for the negative control and Library 1 are the same as that used in Figure 1b.

**Supplementary Figure S3.**
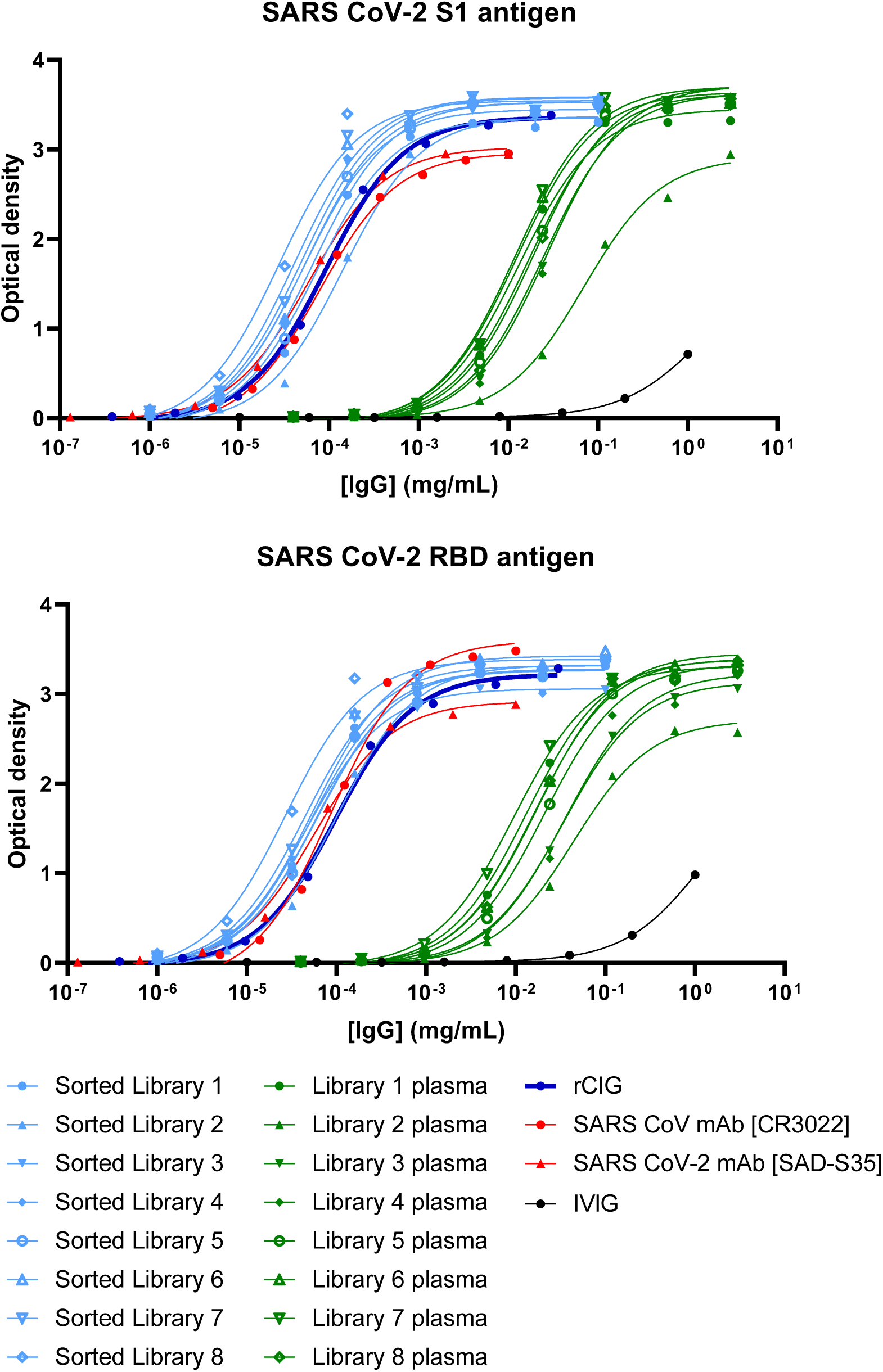
CoV-2 S1- and RBD-specific antibody binding by ELISA. rCIG (dark blue), the 8 recombinant libraries (light blue), the 8 plasma pools from donors that made up rCIG (green), neutralizing SARS-CoV-2 mAb (red triangle), non-neutralizing SARS-CoV mAb (red circle), and IVIG (black) were titrated relative to total IgG concentration.

**Supplementary Figure S4.**
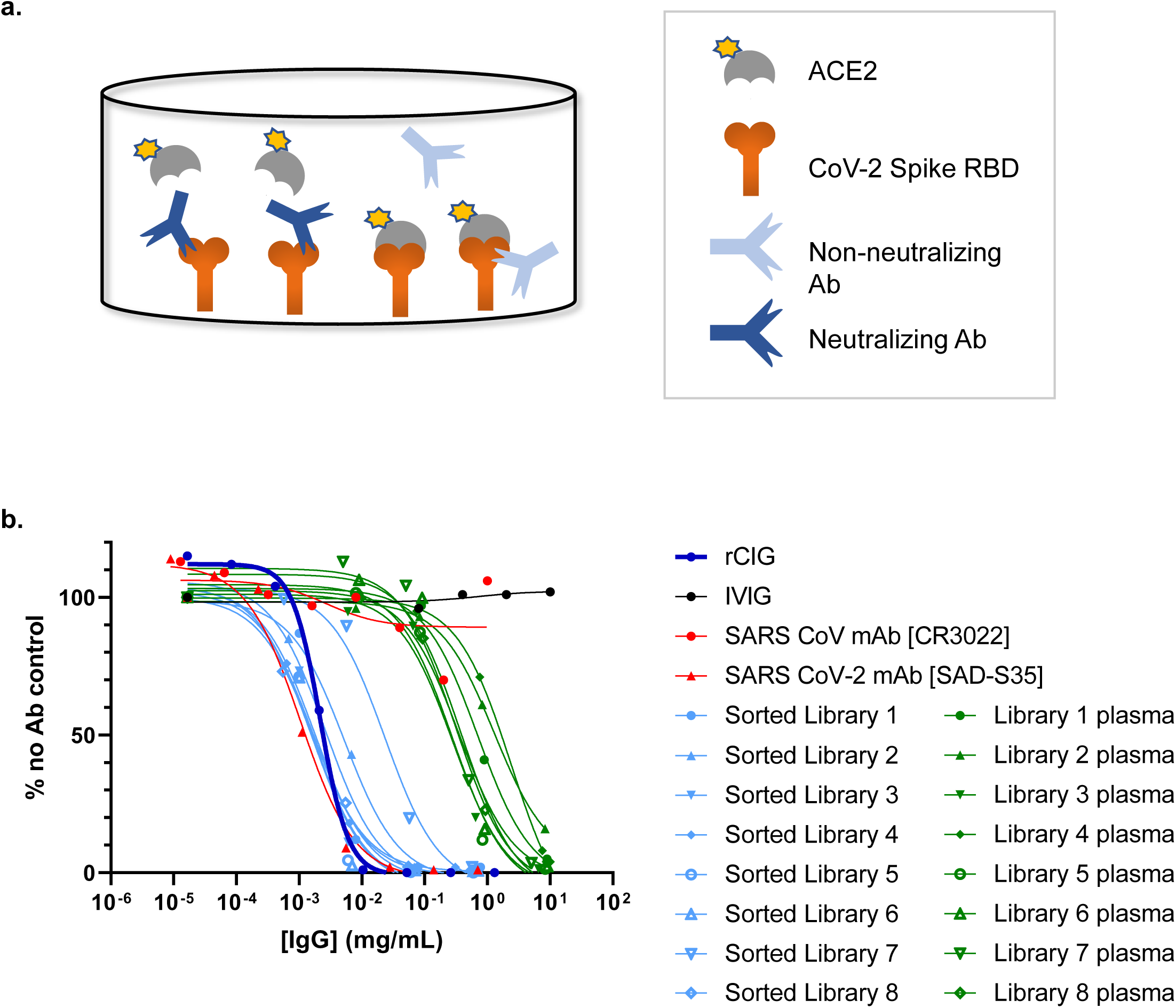
CoV-2 Spike:ACE2 inhibition assay. (**a**) The blocking ability of SARS-CoV-2 specific antibodies were measured using a plate-based ELISA method. Spike RBD was used to coat the plate and after the antibody samples are co-incubated, ACE2 was added and measured for binding to RBD; antibodies that block the interaction demonstrate low or no binding of ACE2 and are considered neutralizing. (**b**) The recombinant polyclonal rCIG (dark blue), the 8 recombinant antibody libraries (light blue), the 8 plasma pools from donors that make up rCIG (green), neutralizing SARS-CoV-2 mAb (red triangle), non-neutralizing SARS-CoV mAb (red circle), and IVIG (black) were titrated and added to the RBD coated plate. The data are reported as “% no Ab control”, i.e., dividing the signal of the test article by the signal of a no Ab control.

**Supplementary Figure S5.**
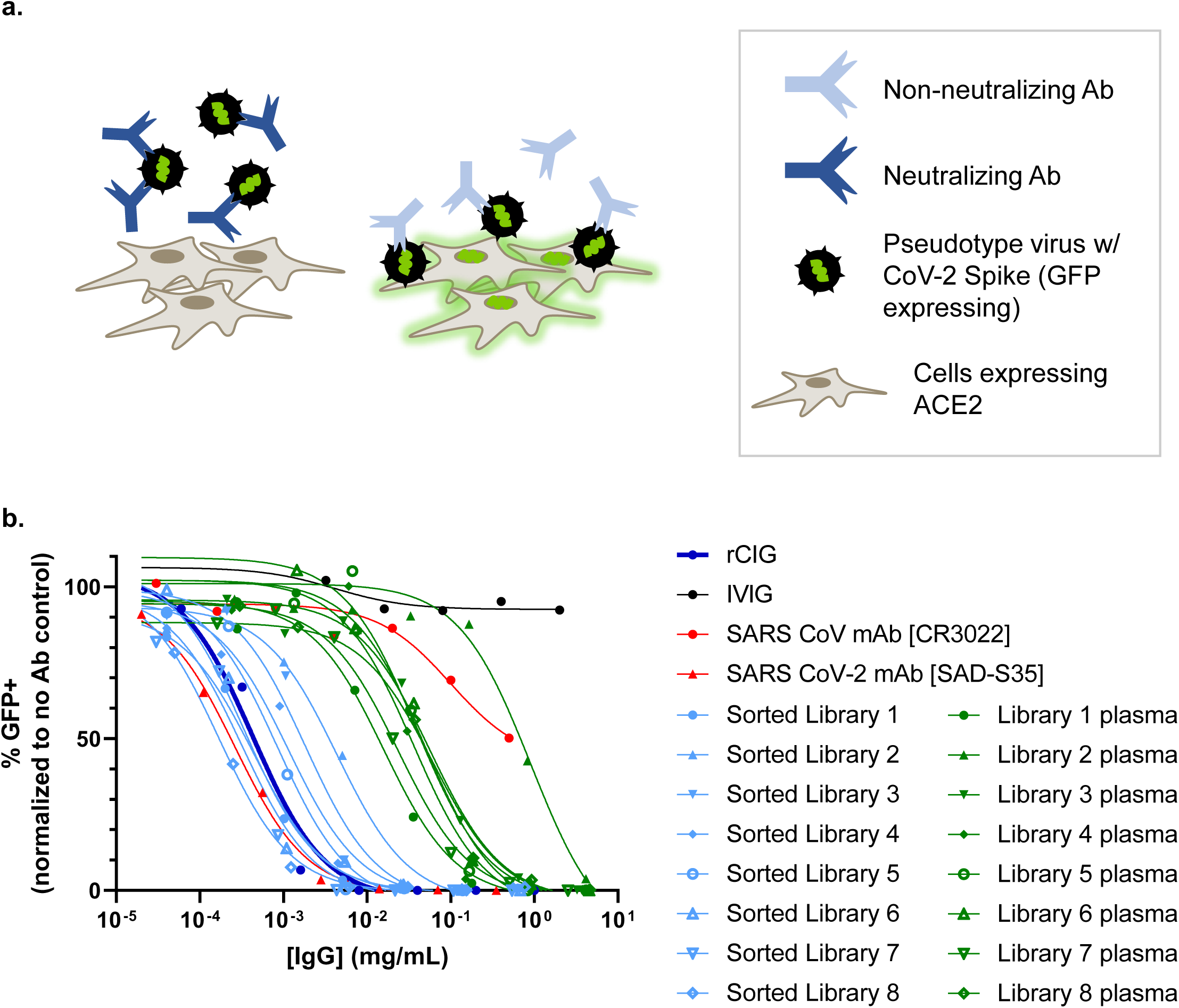
CoV-2 pseudotype virus neutralization assay. (**a**) A pseudotype virus expressing the SARS-CoV-2 spike proteins can infect ACE2-expressing target cells (which then turn green due to GFP expression from the pseudotype virus), which is used to demonstrate whether antibodies specific to SARS-CoV-2 can neutralize infection. (**b**) The recombinant polyclonal rCIG (dark blue), the 8 recombinant antibody libraries (light blue), the eight plasma pools from donors that make up rCIG (green), neutralizing SARS-CoV-2 mAb (red triangle), non-neutralizing SARS-CoV mAb (red circle), and IVIG (black) were titrated and added to ACE2-expressing cells in the presence of CoV-2 pseudotype virus. The percent of infected cells (GFP+) was quantified by flow cytometry and was normalized by dividing by the GFP+ signal in the negative control wells, which lacked test article.

**Supplementary Figure S6.**
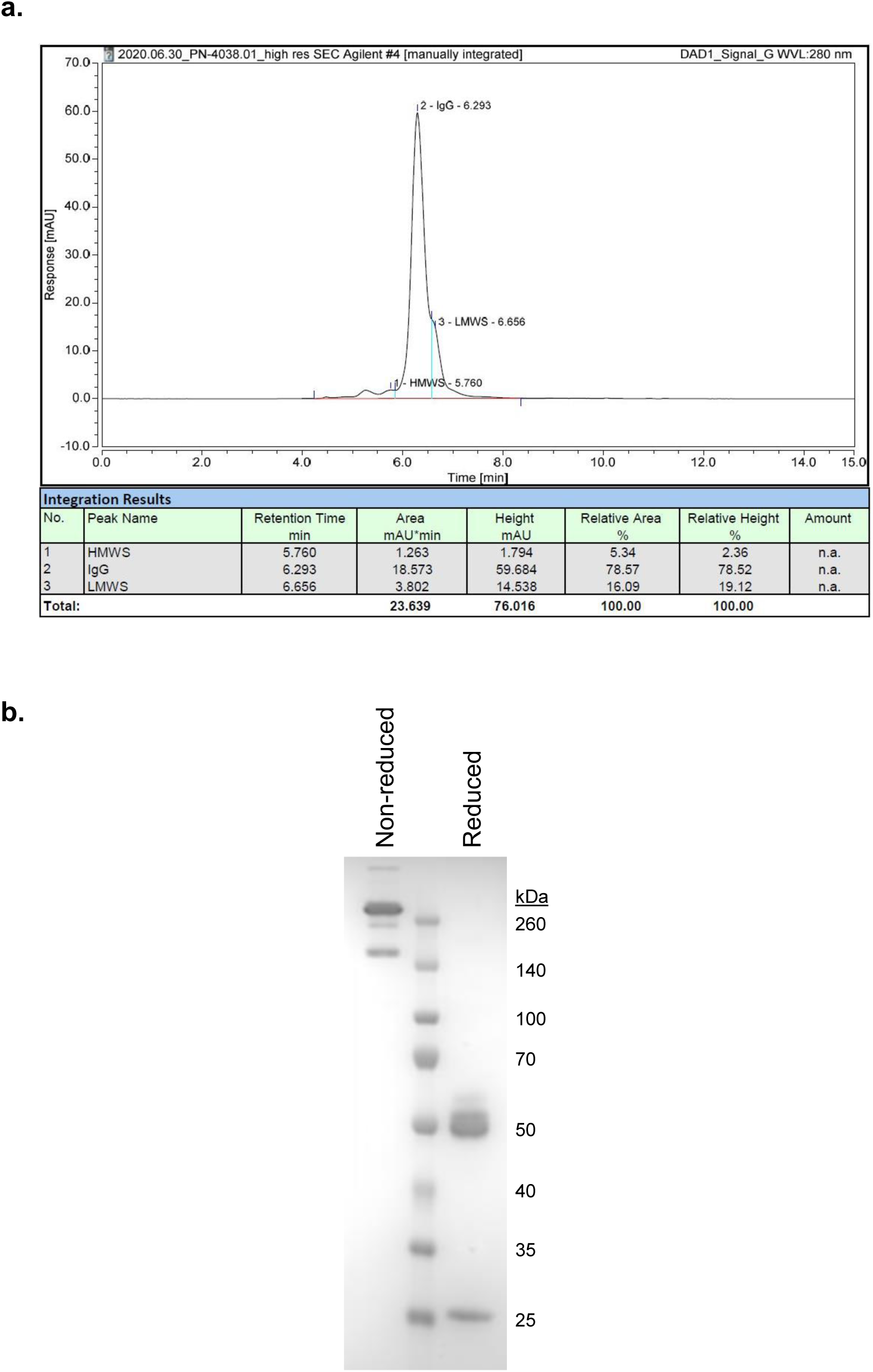
Quality control analysis of purified rCIG protein. (**a**) SEC-HPLC and (**b**) SDS-PAGE analysis were used to assess the purity of the Protein A-purified protein.

**Supplementary Figure S7.**
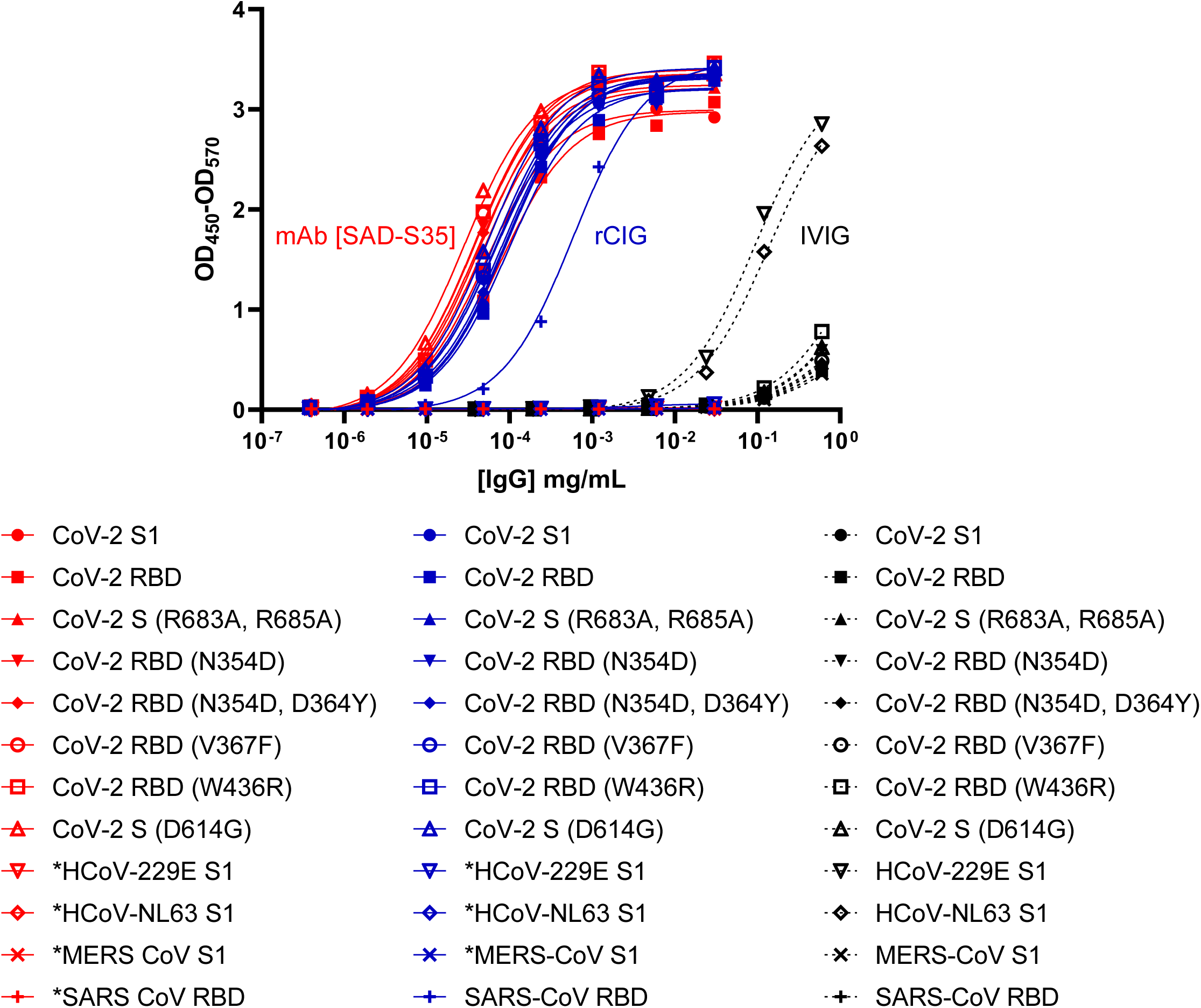
Antibody reactivity to SARS-CoV-2 variants and other coronaviruses. ELISA plates were coated with 2 µg/mL of spike or RBD proteins from known circulating variants of SARS-CoV-2, SARS-CoV, MERS, and other human coronaviruses (HCoV). The binding ability of rCIG (blue), SARS-CoV-2 neutralizing mAb (SAD-S35, red), and IVIG (black) was determined for each antigen. *, No binding was observed against the indicated antigen. While rCIG and the mAb had poly-variant specific responses to all SARS-CoV-2 variants, only rCIG bound SARS-CoV RBD while the mAb did not. IVIG had no specific responses to SARS-CoV-2, SARS-CoV or MERS but did have a weak response to HCoV-229E and HCoV-NL63.

**Supplementary Figure S8.**
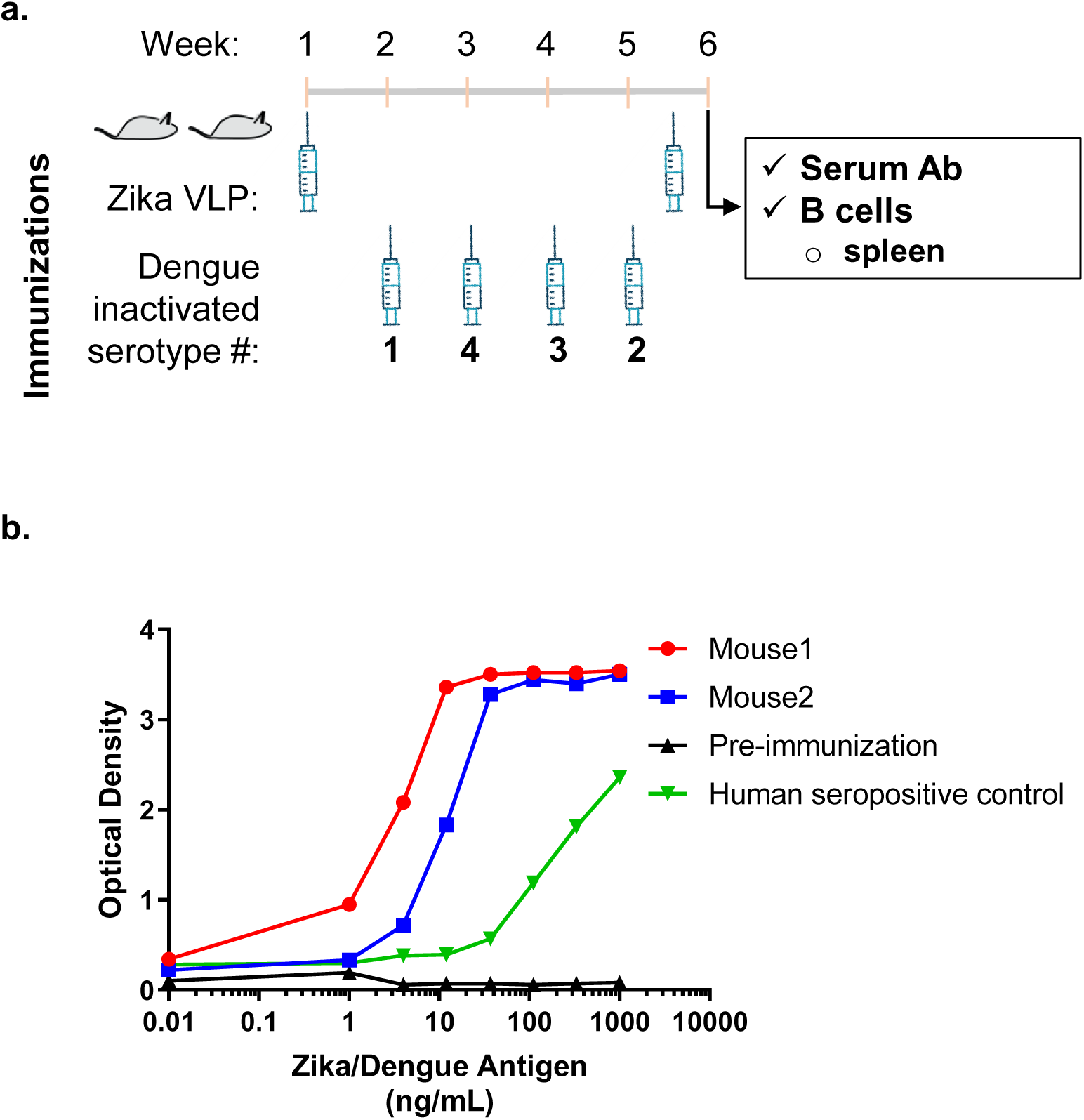
Immunization induced antibody responses to Zika and Dengue in humanized mice. (**a**) Two Trianni mice were immunized weekly with Zika virus like particles (VLP), inactivated Dengue 1, Dengue 4, Dengue 3, or Dengue 2 as indicated in the figure. (**b**) After week 5, serum from the mice was tested for antibody response against a mixture of Zika and Dengue antigens and compared to pre-immunization or human seropositive controls to confirm antigen-specific antibody responses.

**Supplementary Figure S9.**
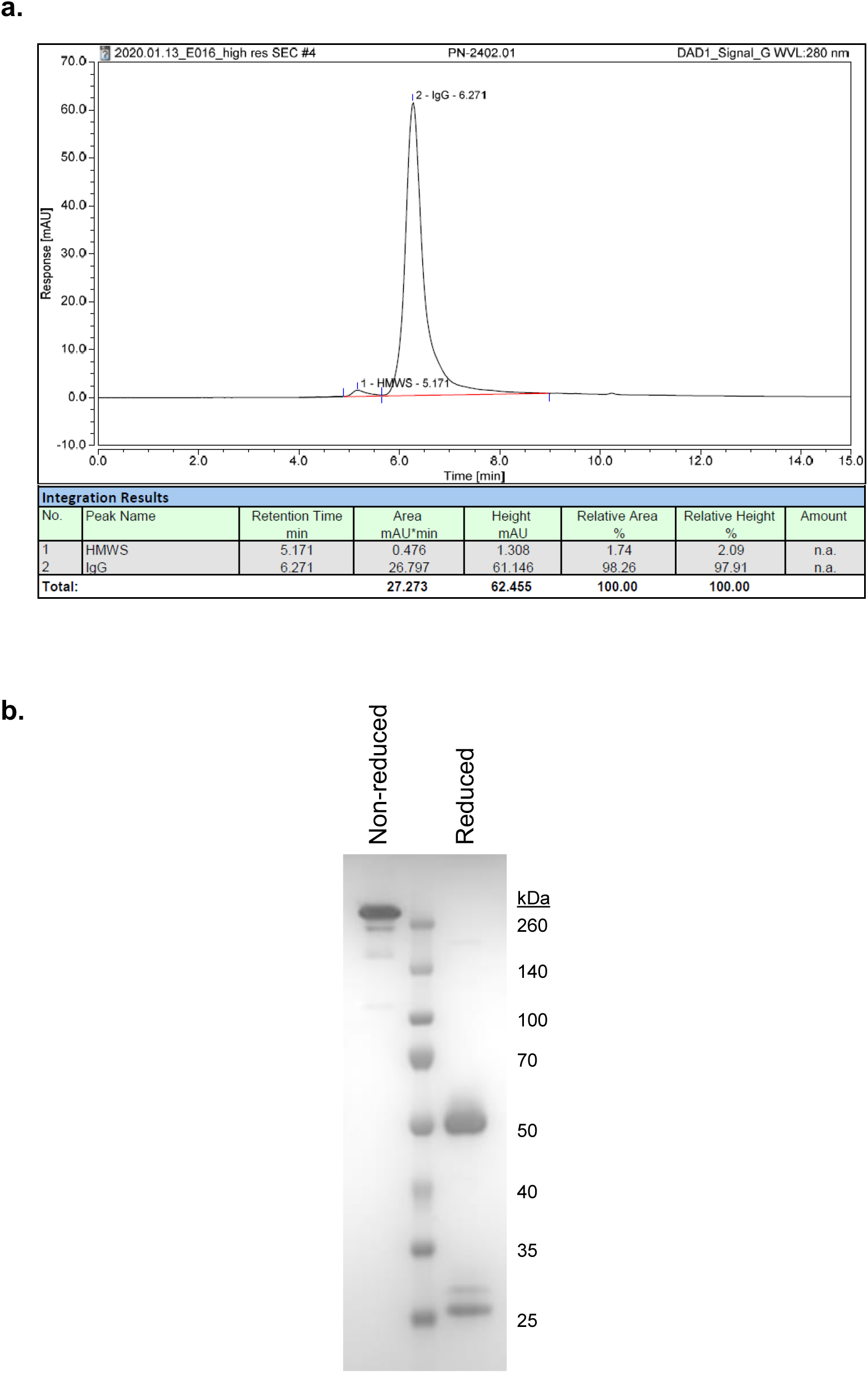
Quality control analysis of purified rZIG-IgG1 protein. (**a**) SEC-HPLC and (**b**) SDS-PAGE analysis were used to assess the purity of the Protein A-purified protein.

**Supplementary Figure S10.**
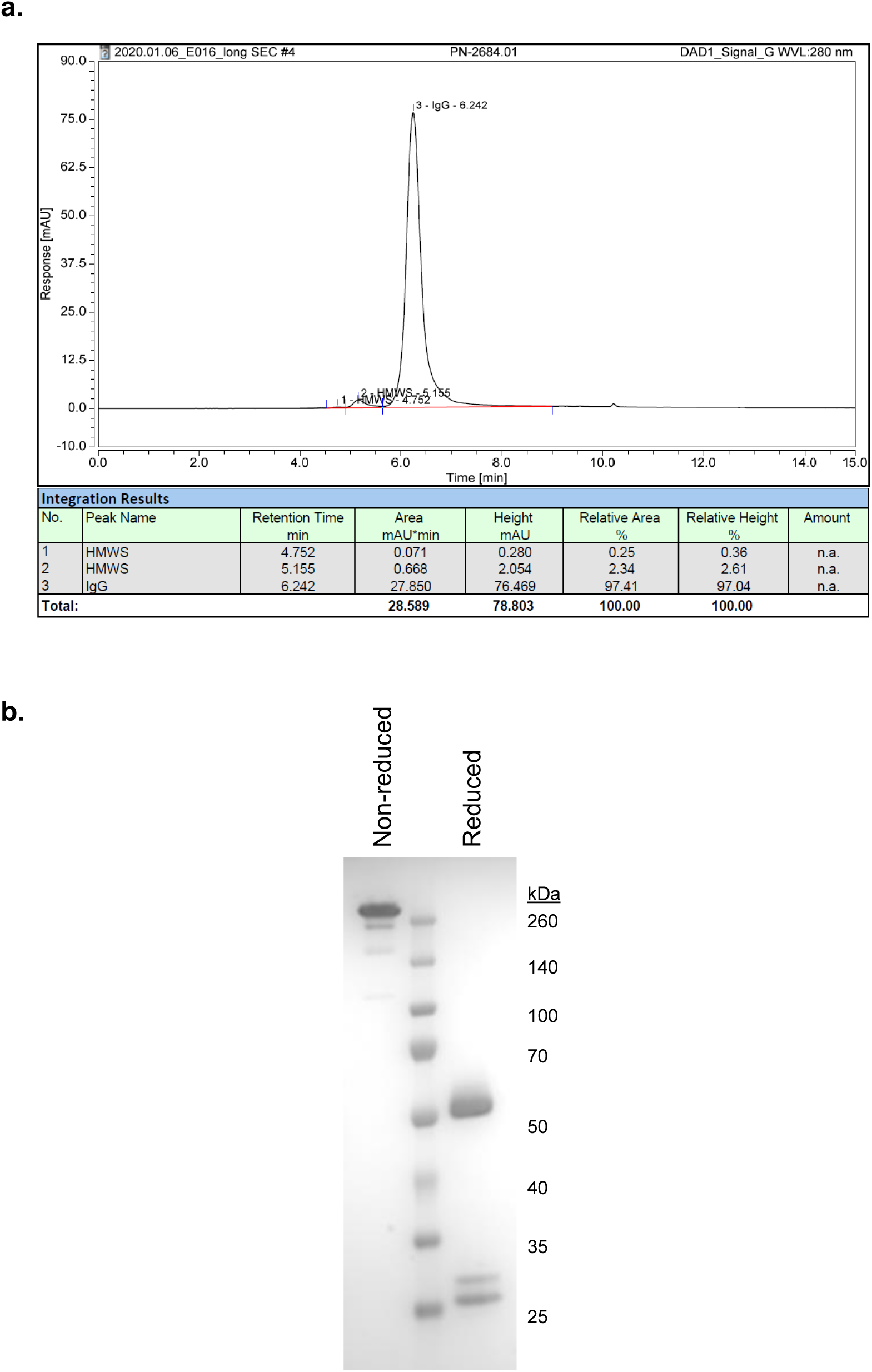
Quality control analysis of purified rZIG-LALA protein. (**a**) SEC-HPLC and (**b**) SDS-PAGE analysis were used to assess the purity of the Protein A-purified protein.

**Supplementary Figure S11.**
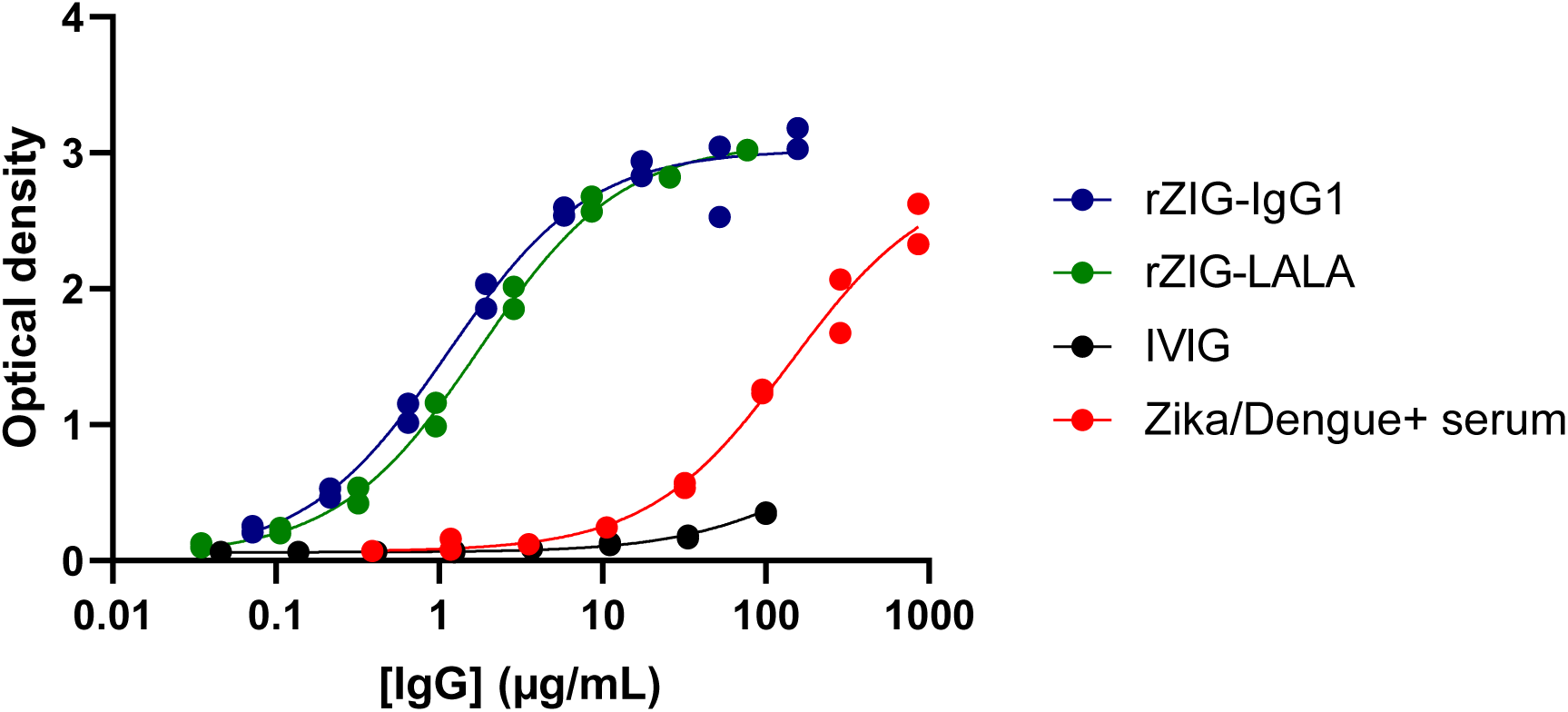
rZIG binding to Zika by ELISA. rZIG-IgG1 (blue), rZIG-LALA (green), negative control IVIG (black), and positive control Zika/Dengue positive serum (red) were serially diluted and added to a Zika envelope-coated plate. Antigen-specific responses were quantified by an anti-human-HRP secondary antibody.

**Supplementary Figure S12.**
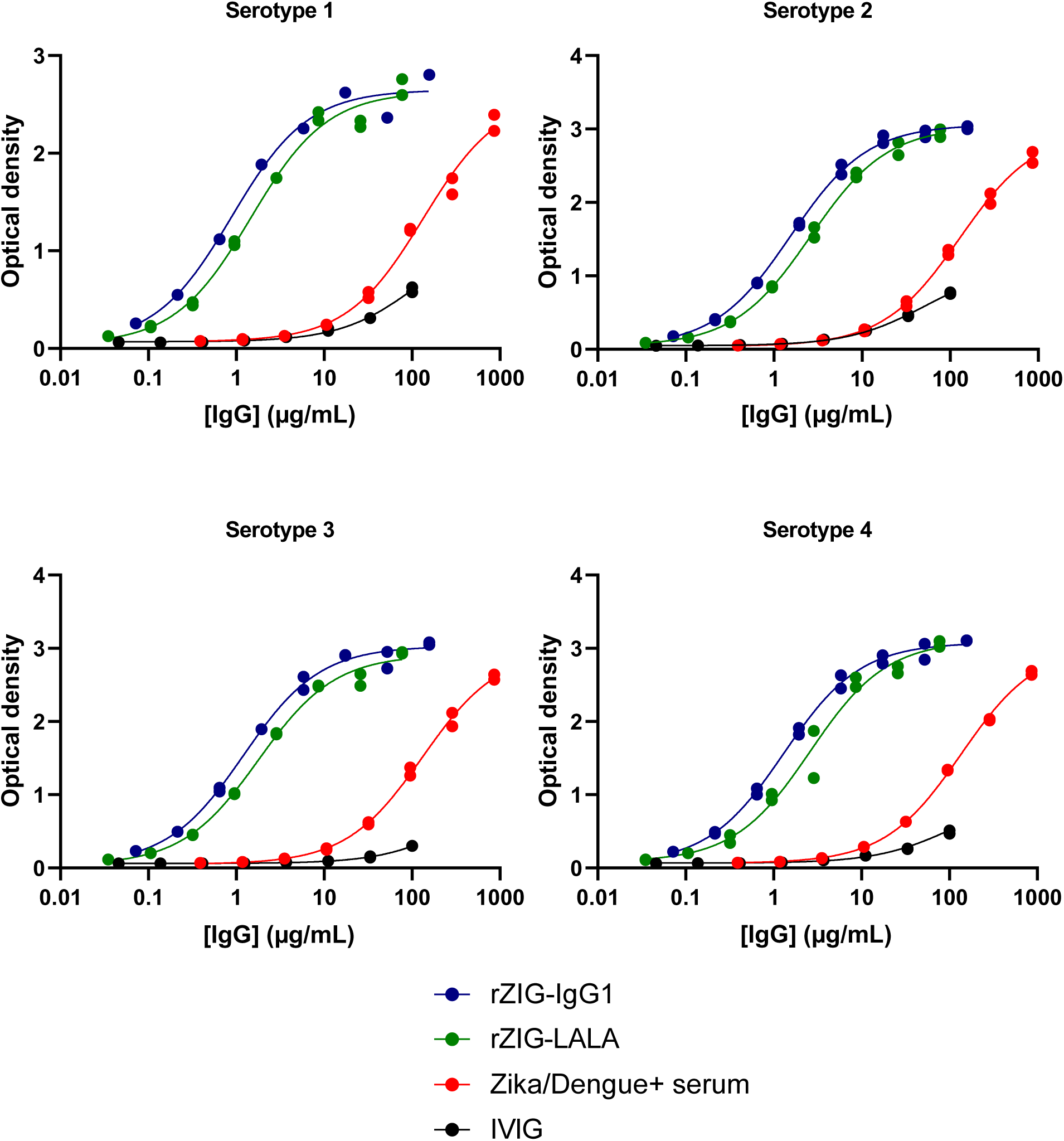
rZIG binding to Dengue serotypes by ELISA. rZIG-IgG1 (blue), rZIG-LALA (green), negative control IVIG (black), and positive control Zika/Dengue positive serum (red) were serially diluted and added to a Dengue serotypes 1, 2, 3, or 4 envelope-coated plates. Antigen-specific responses were quantified by an anti-human-HRP secondary antibody.

**Supplementary Figure S13.**
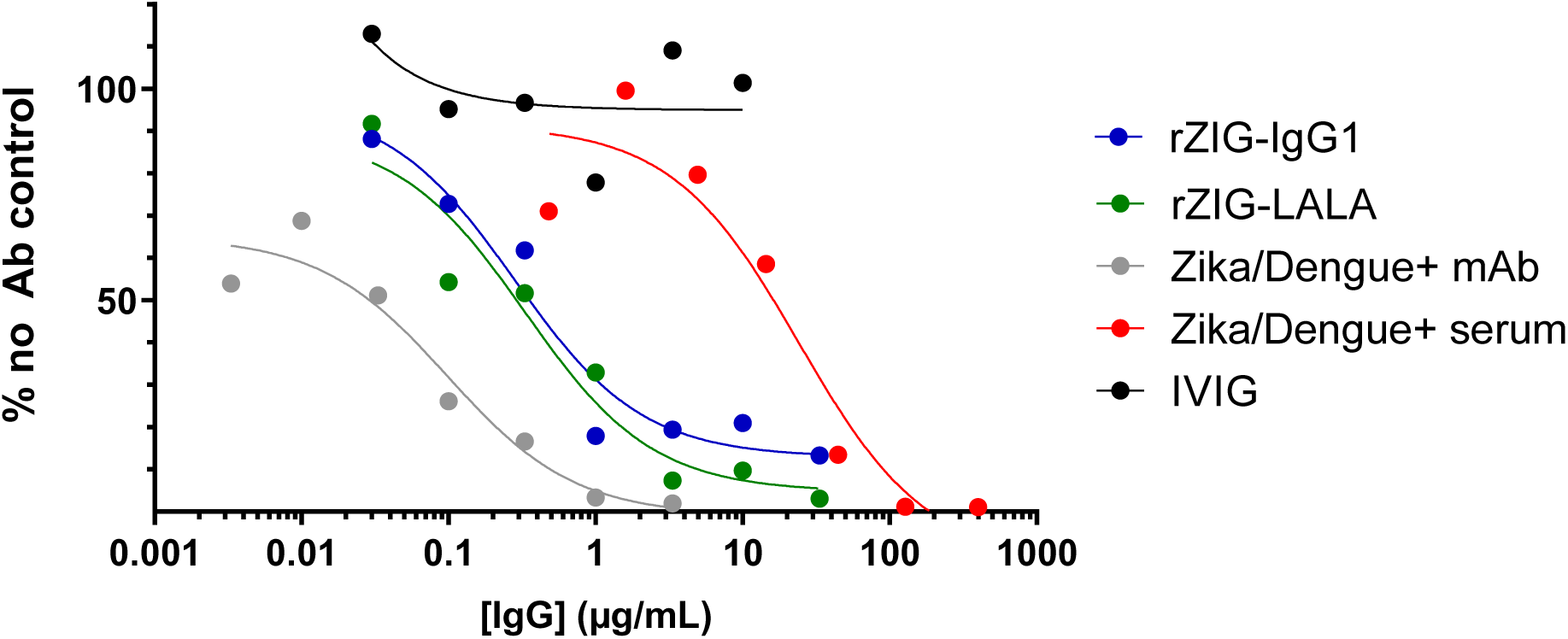
Zika pseudotype virus neutralization assay. rZIG-IgG1 (blue), rZIG-LALA (green), negative control IVIG (black), positive control Zika/Dengue mAb (gray), and positive control Zika/Dengue positive serum (red) were serially diluted, co-incubated with Zika pseudotype virus, and added to BHK/DC-SIGN target cells. Antibody induced neutralization was quantified by the infection-induced luciferase expression divided by luciferase expression in the no Ab control.

**Supplementary Figure S14.**
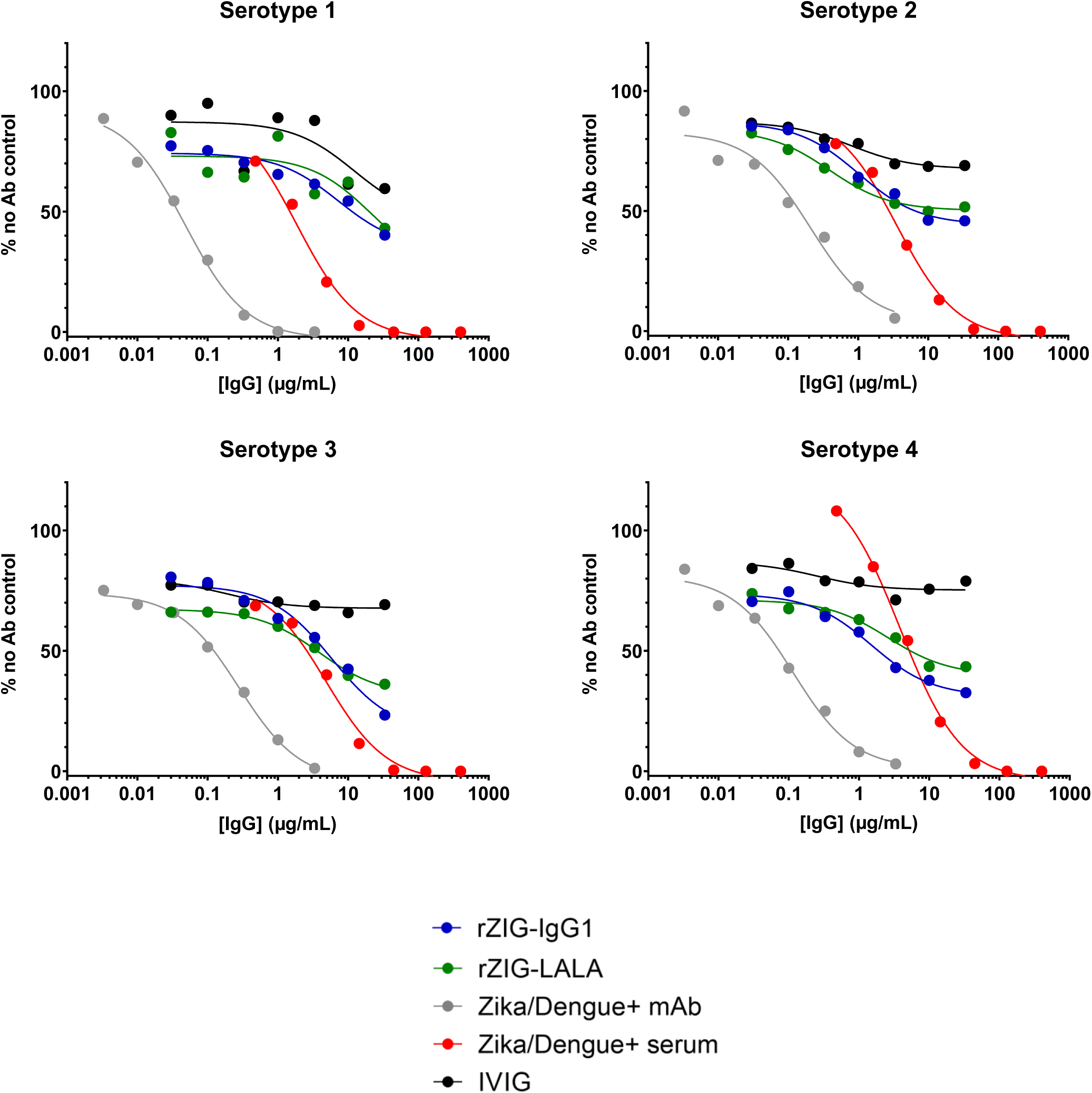
Dengue serotype pseudotype virus neutralization assay. rZIG-IgG1 (blue), rZIG-LALA (green), negative control IVIG (black), positive control Zika/Dengue mAb (gray), and positive control Zika/Dengue positive serum (red) were serially diluted, co-incubated with pseudotype virus expressing the indicated Dengue serotype envelope antigens, and added to BHK/DC-SIGN target cells. Antibody induced neutralization was quantified as “% no Ab control”, or infection-induced luciferase expression divided by the luciferase expression in the no Ab control.

**Supplementary Figure S15.**
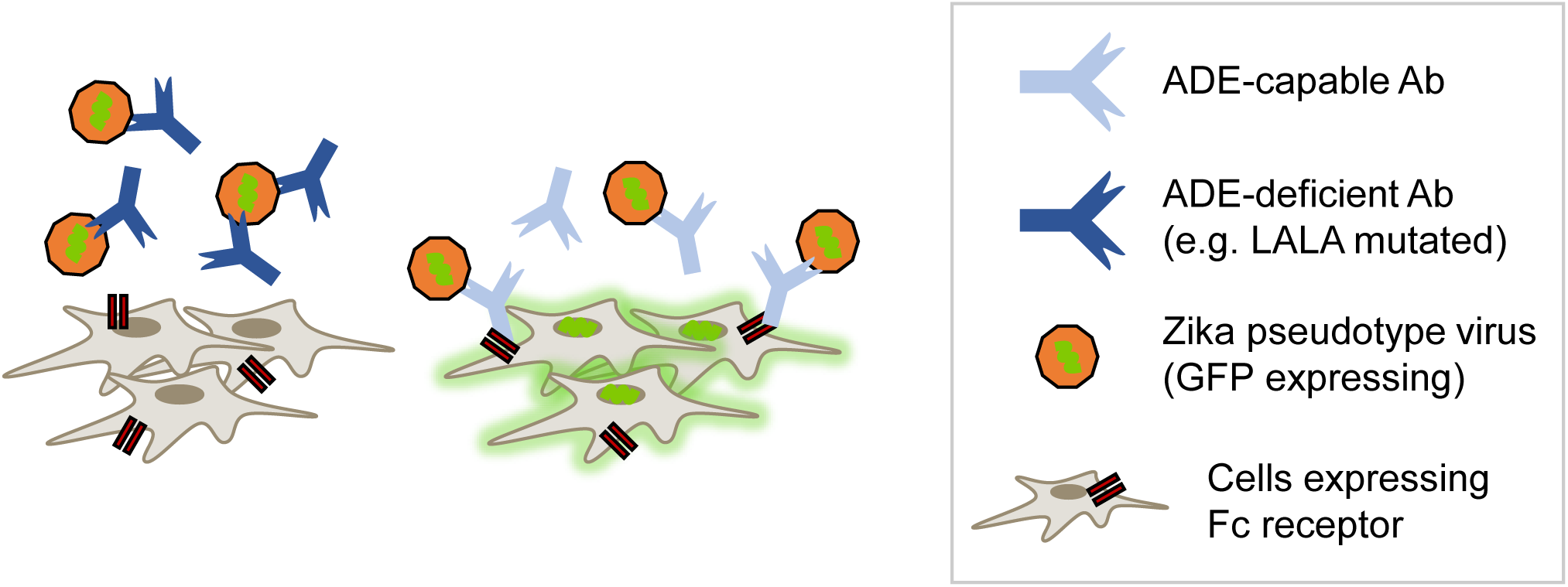
Schematic of antibody dependent enhancement (ADE) assay. Antibody bound to pseudotype virus can infect cells through the Fc receptor expressed on target cells. By introducing an Fc mutation that prevents FcR binding (e.g. LALA), antibody-induced viral infection is abrogated.

**Supplementary Figure S16.**
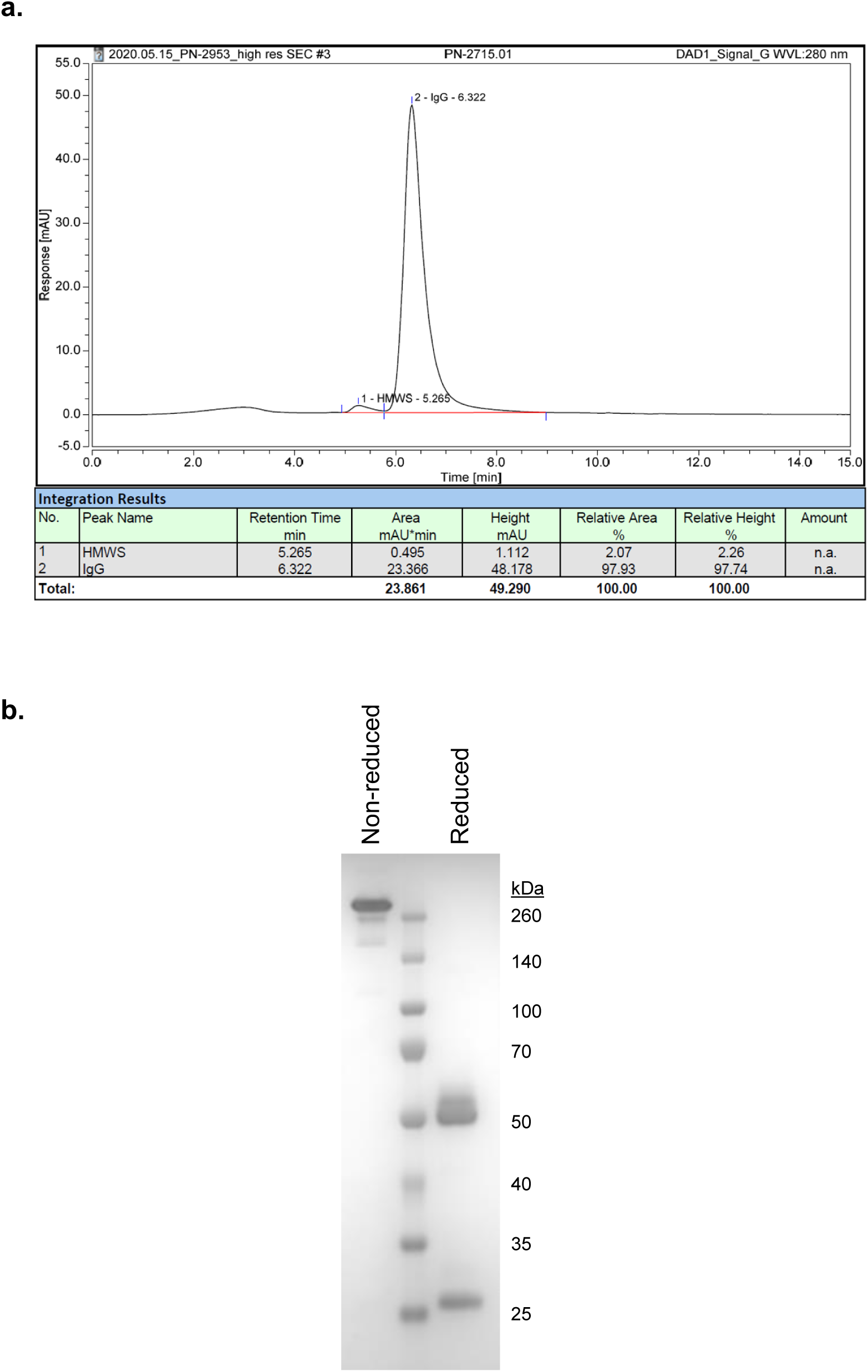
Quality control analysis of purified rHIG protein. (**a**) SEC-HPLC and (**b**) SDS-PAGE analysis were used to assess the purity of the Protein A-purified protein.

**Supplementary Figure S17.**
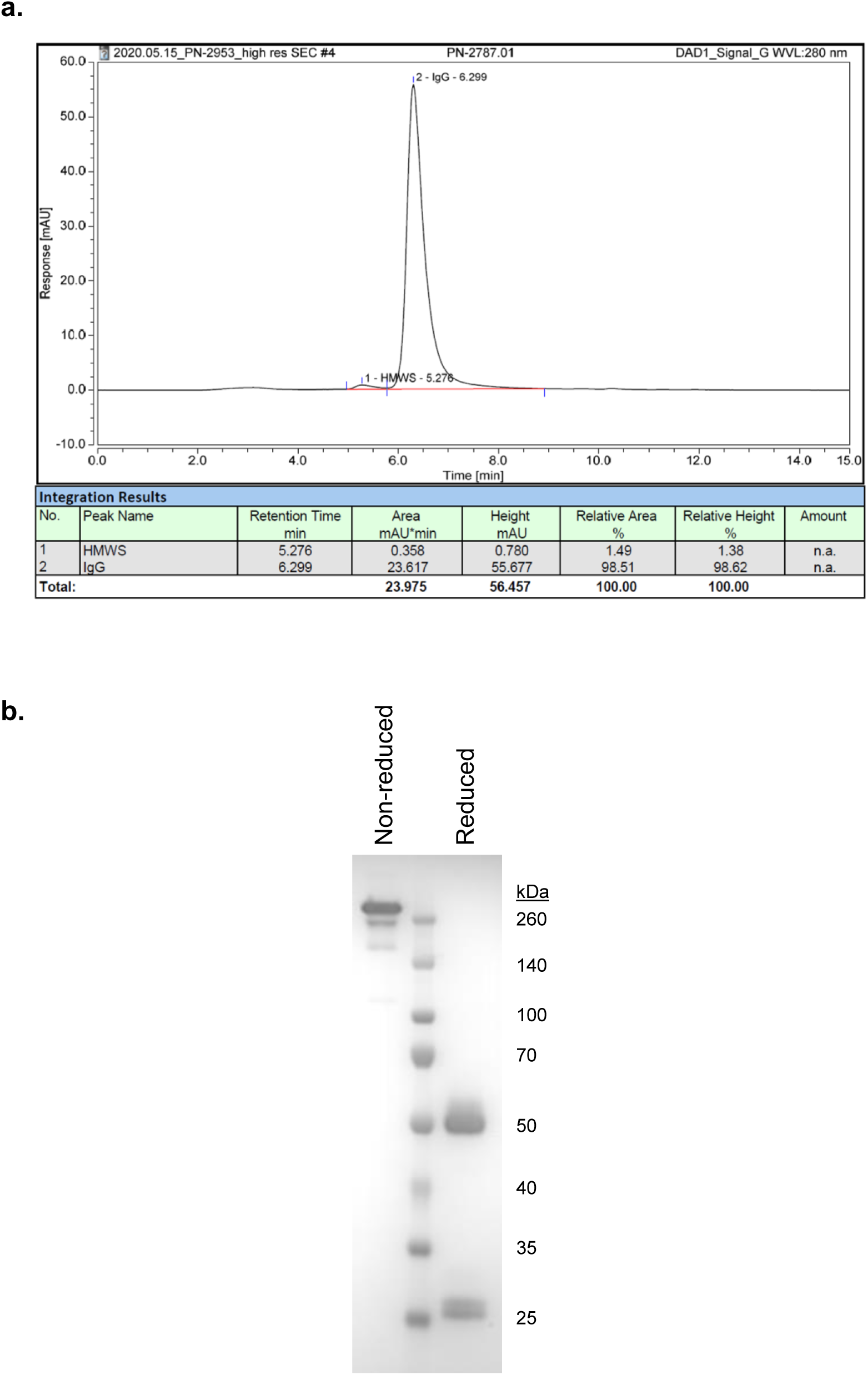
Quality control analysis of purified rPIG protein. (**a**) SEC-HPLC and (**b**) SDS-PAGE analysis were used to assess the purity of the Protein A-purified protein.

**Supplementary Figure S18.**
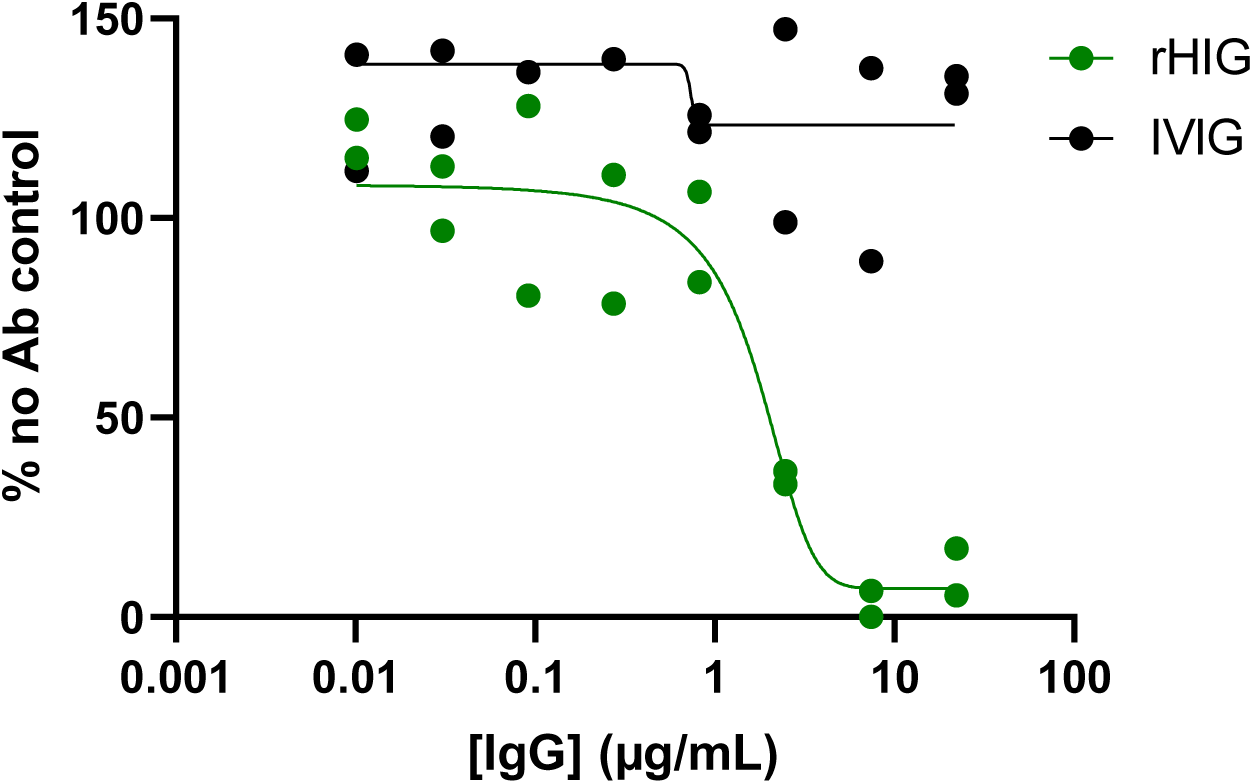
Haemophilus influenzae serum bactericidal assay (SBA). rHIG (green) and IVIG (black) were serially diluted and co-incubated with 5×10^4^ CFU/mL *Haemophilus influenzae* Eagan strain. After incubation, complement was added, incubated, and test samples were plated on chocolate agar. After 16 hours incubation, bacteria colony counts for each serial dilution were quantified and divided by the bacteria colony counts in the no Ab control.

**Supplementary Figure S19.**
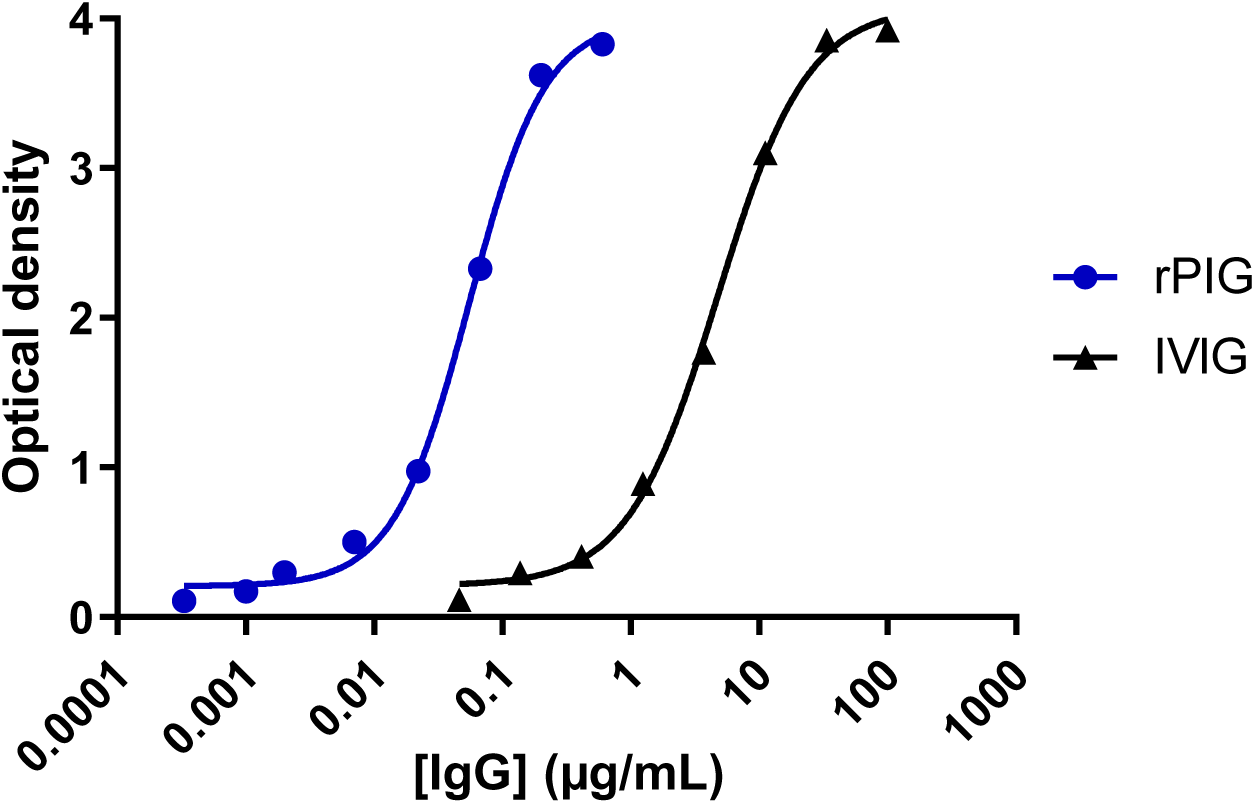
Pneumococcal antibody binding by ELISA. Binding of serially diluted rPIG (blue) and IVIG (black) to a pool of 23 pneumococcal polysaccharides was measured by ELISA.

**Supplementary Figure S20.**
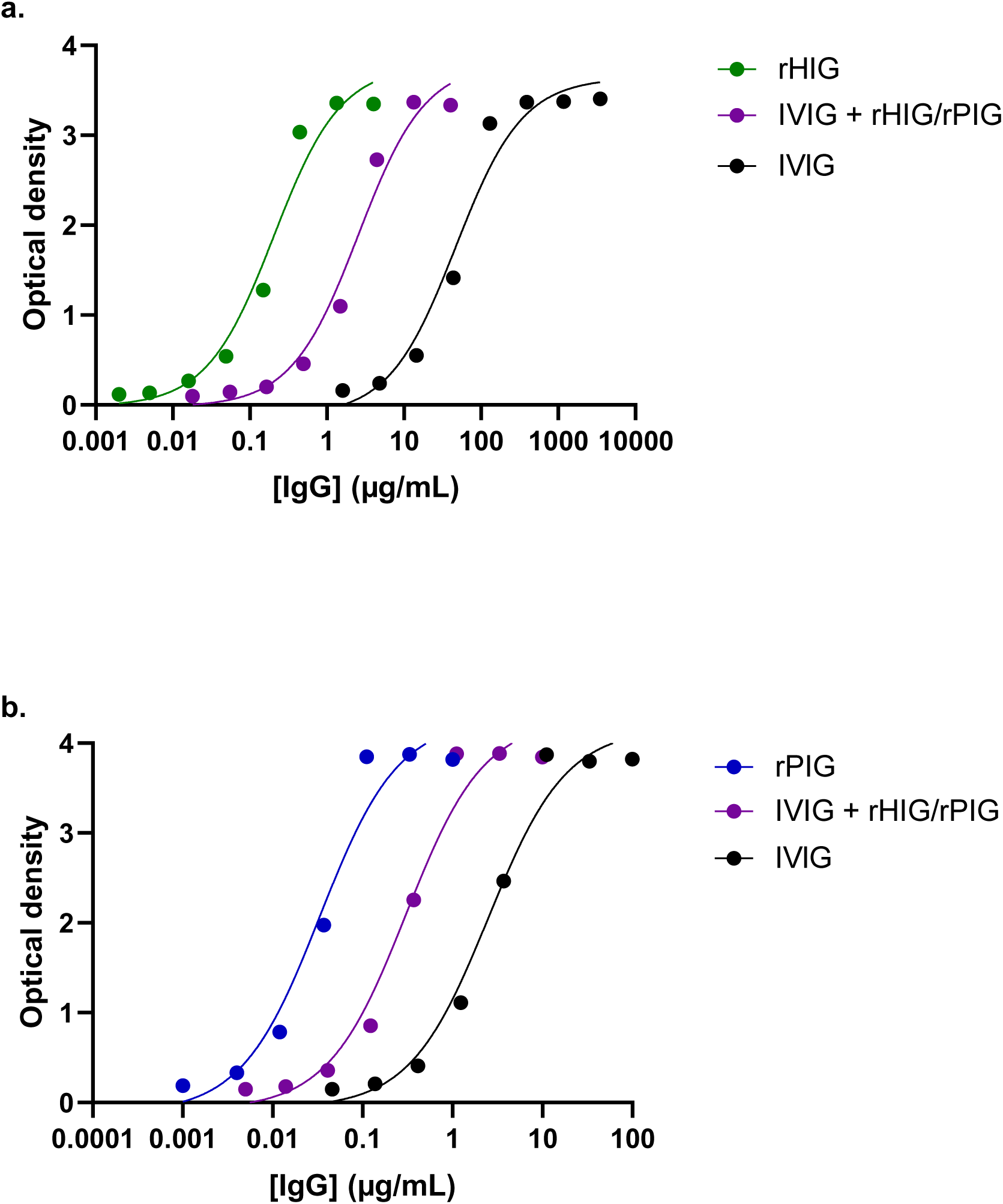
Pneumococcal or Hib antibody binding of IVIG + rHIG/rPIG by ELISA. (**a**) Binding of serially diluted rHIG (green), IVIG + rHIG/rPIG (purple), and IVIG (black) to Hib was measured by ELISA. (**b**) Binding of serially diluted rPIG (blue), IVIG + rHIG/rPIG (purple), and IVIG (black) to a pool of 23 pneumococcal polysaccharides was measured by ELISA.

**Supplementary Figure S21.**
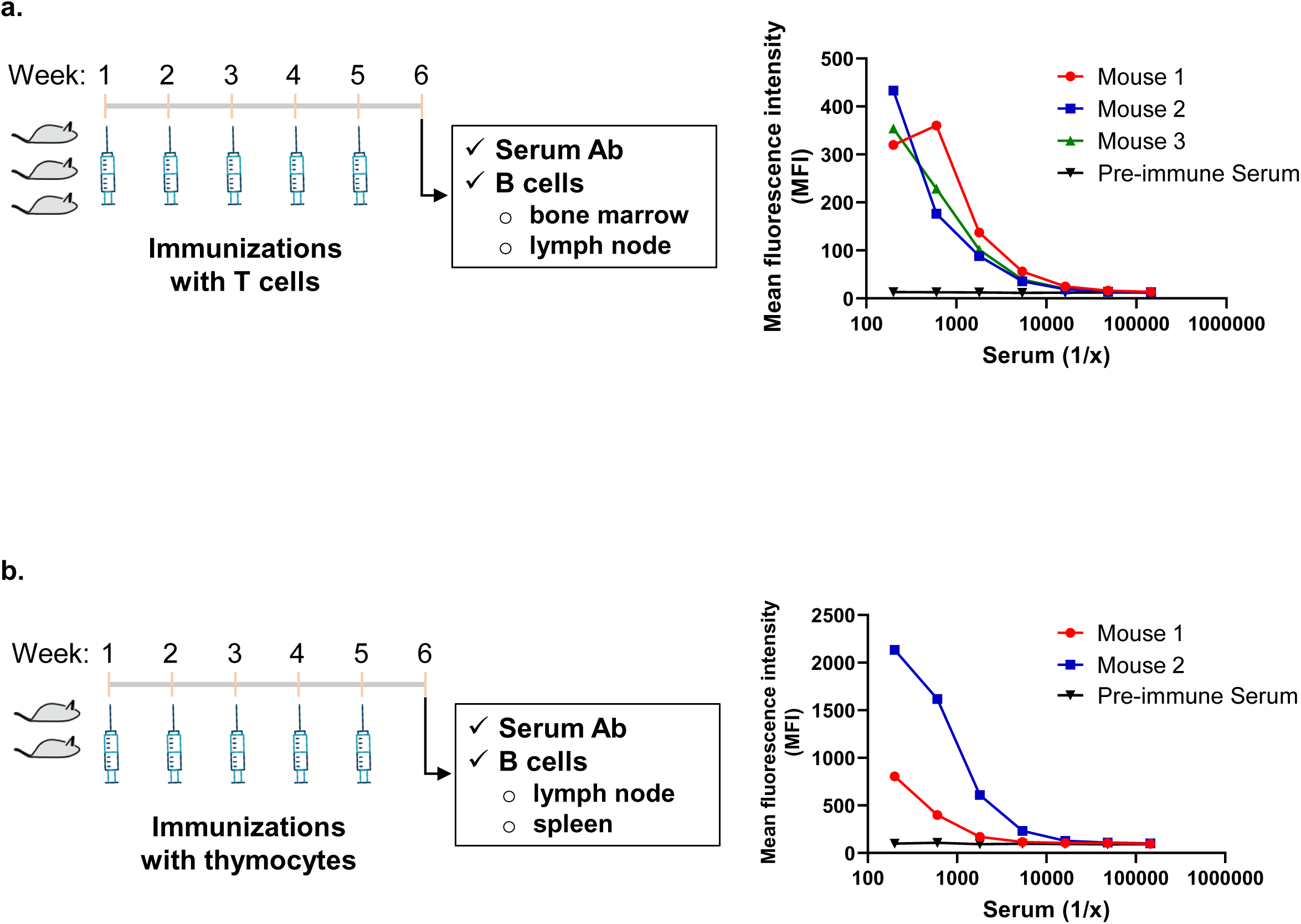
Immunization induced antibody responses to human T cells and thymocytes. (**a**) Three Trianni mice were immunized weekly with T cells isolated from one human donor with ALD/MDP adjuvant. After week 5, serum from the mice was tested to confirm binding to T cells before a final boost without adjuvant 5 days prior to harvesting the organ B cells. (**b**) Two Trianni mice were immunized weekly with thymocytes isolated from five separate human donors with ALD/MDP adjuvant. After week 5, serum from the mice was tested to confirm binding to T cells before a final boost without adjuvant 5 days prior to harvesting the organ B cells.

**Supplementary Figure S22.**
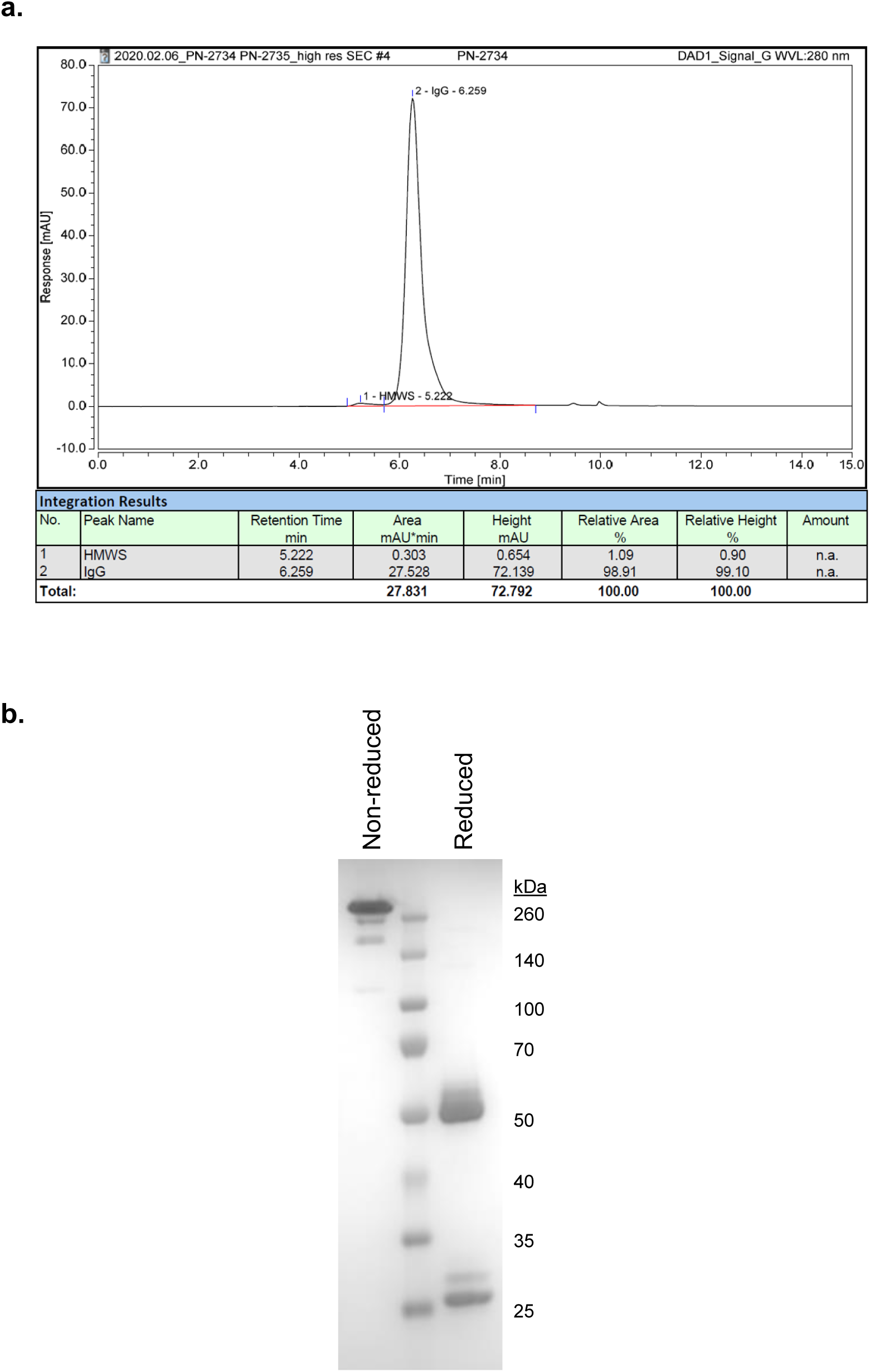
Quality control analysis of purified rhATG protein. (**a**) SEC-HPLC and (**b**) SDS-PAGE analysis were used to assess the purity of the Protein A-purified protein.

**Supplementary Figure S23.**
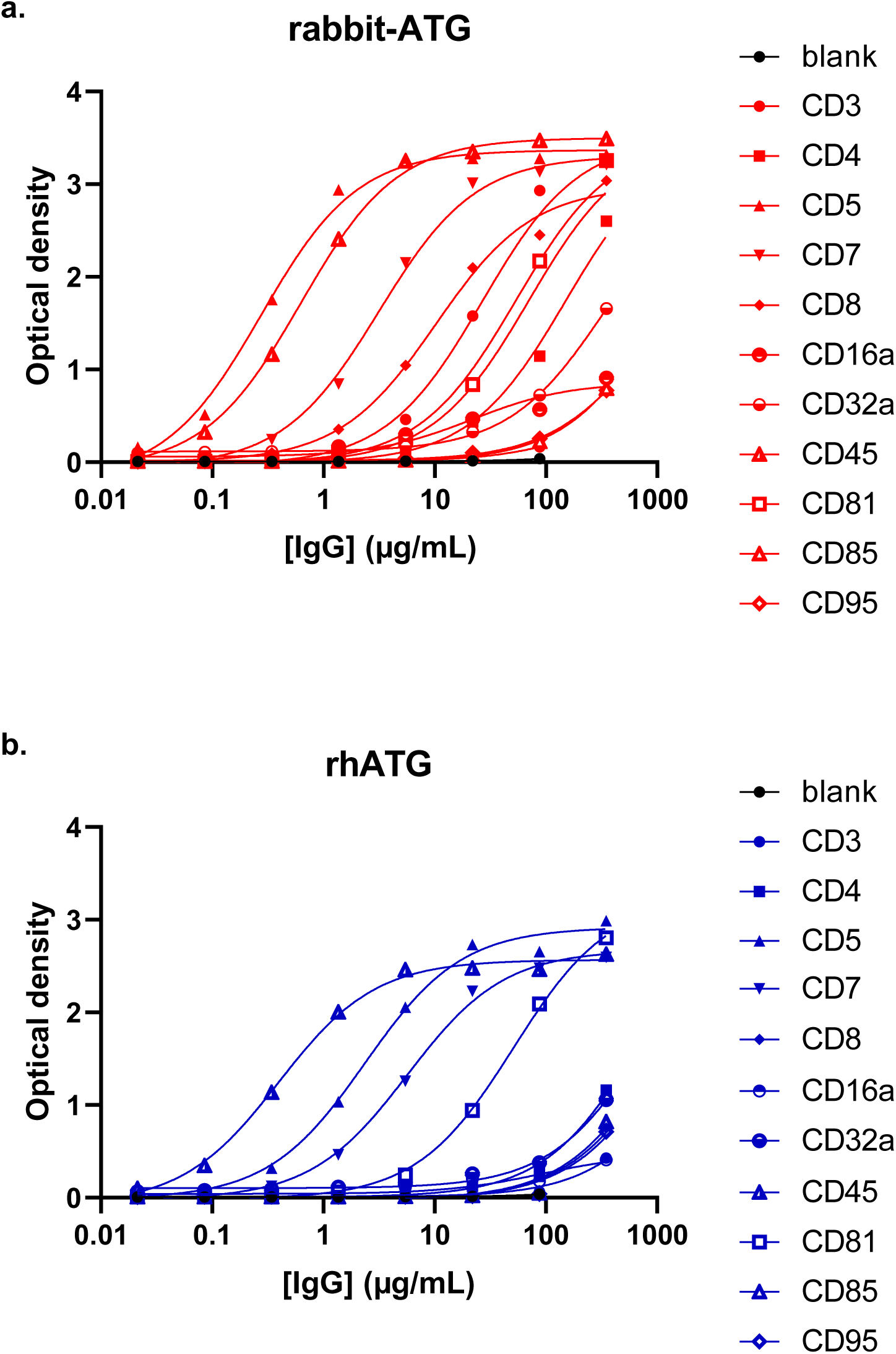
ATG immune cell-specific antibody responses measure by ELISA. The indicated Immune cell antigens were coated onto ELISA plates. **(a)** rabbit-ATG (red) and **(b)** rh-ATG (blue) were serially diluted and added to the plate. Antibody bound to antigens were quantified by anti-rabbit-HRP or anti-human-HRP, respectively.

**Supplementary Figure S24.**
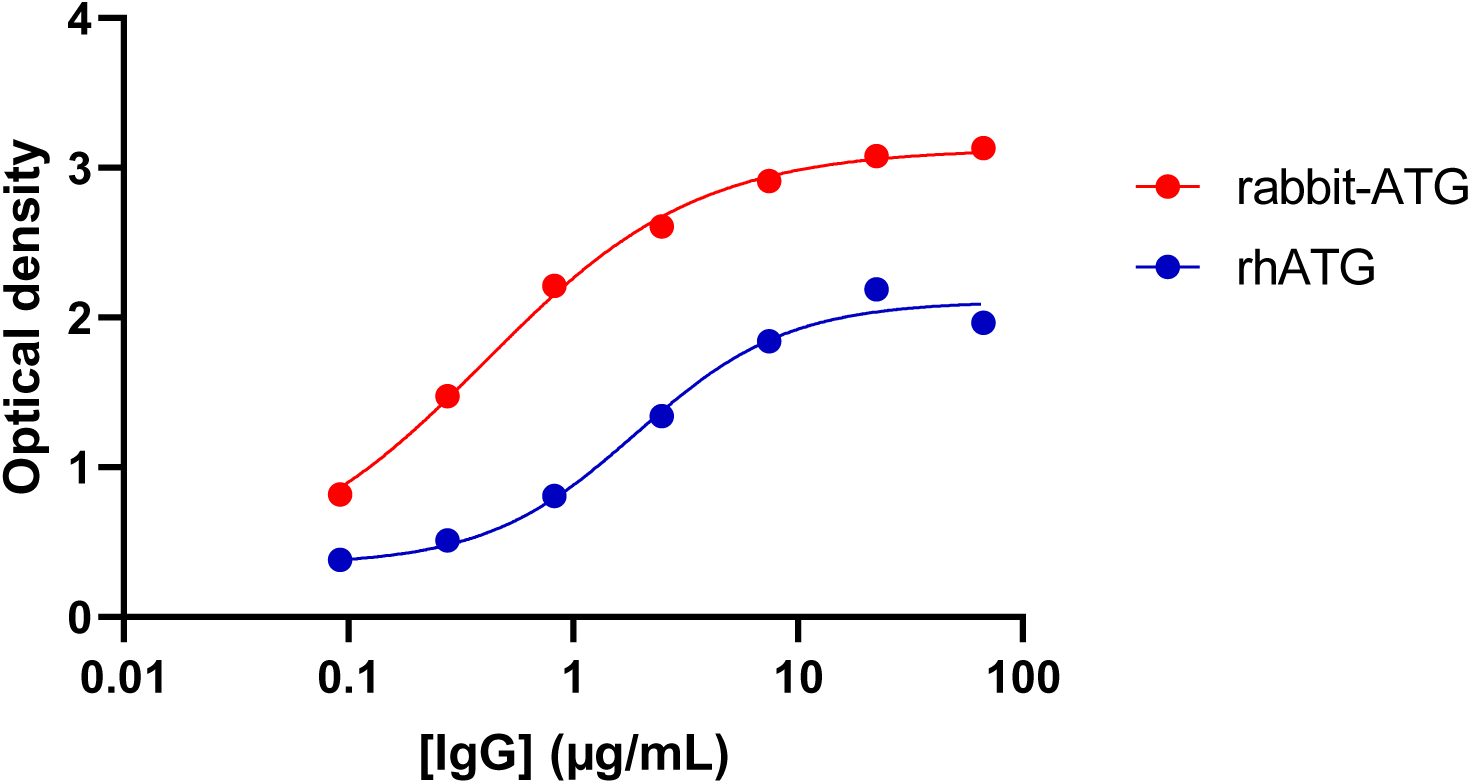
ATG binding to red blood cells by ELISA. RBC-specific antibody response was measured by Immucor Capture-R ELISA. rabbit-ATG (red) and rhATG (blue) were serially diluted and added to the Immucor Capture-R plate. RBC-bound antibodies were quantified by anti-rabbit-HRP or anti-human-HRP, respectively.

**Supplementary Figure S25.**
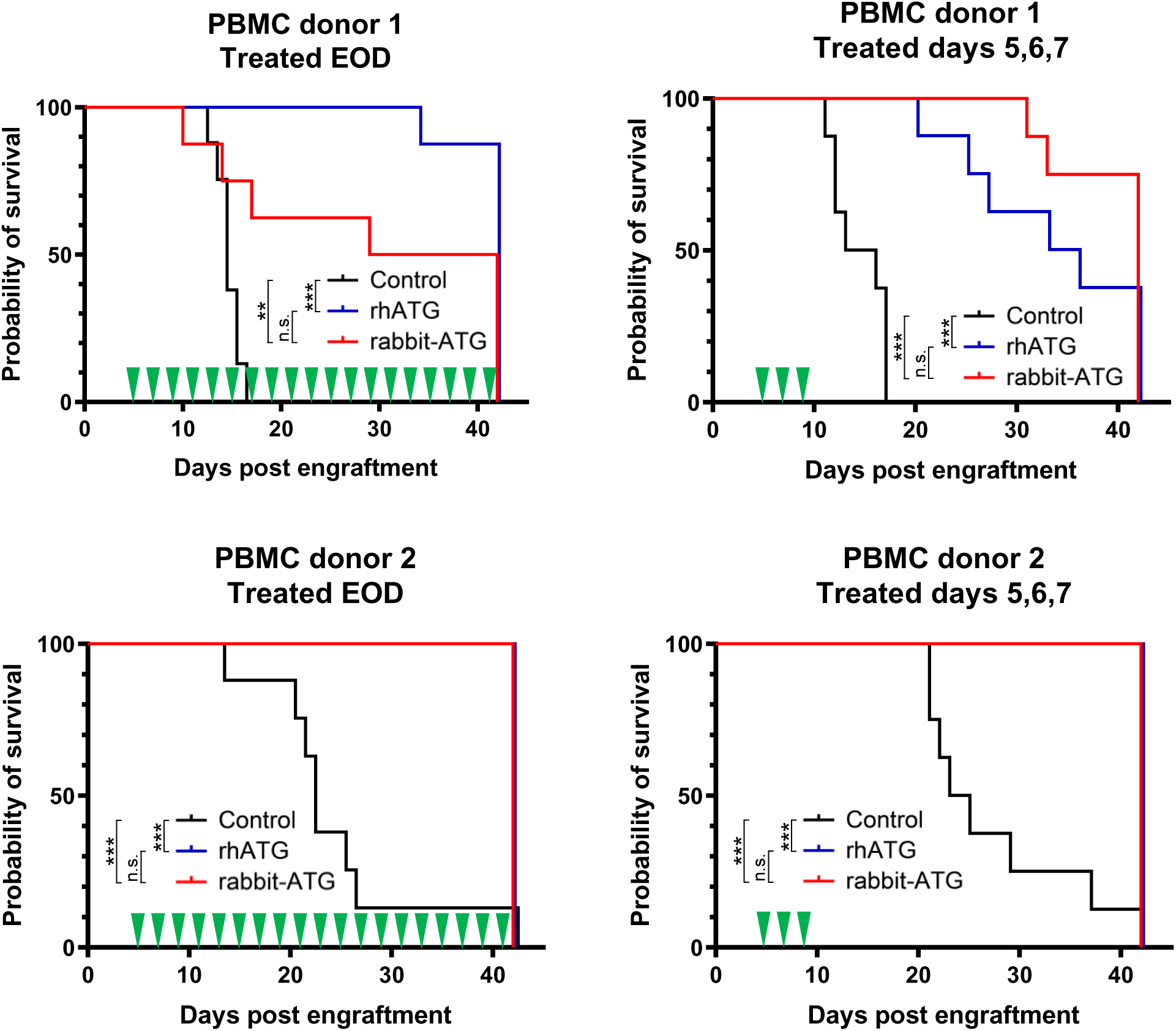
Survival of mice in the GVHD study after ATG treatment. Eight animals per treatment group were engrafted with 10^7^ PBMC from one of two donors. Animals were treated with rhATG (blue), rabbit-ATG (red), or vehicle control (black) either every other day beginning at day 5 or on days 5, 6, and 7 (treatment days are indicated by green triangles), then monitored for progression to GVHD and death. ** p<0.01, *** p<0.001, n.s. not significant.

**Supplementary Figure S26.**
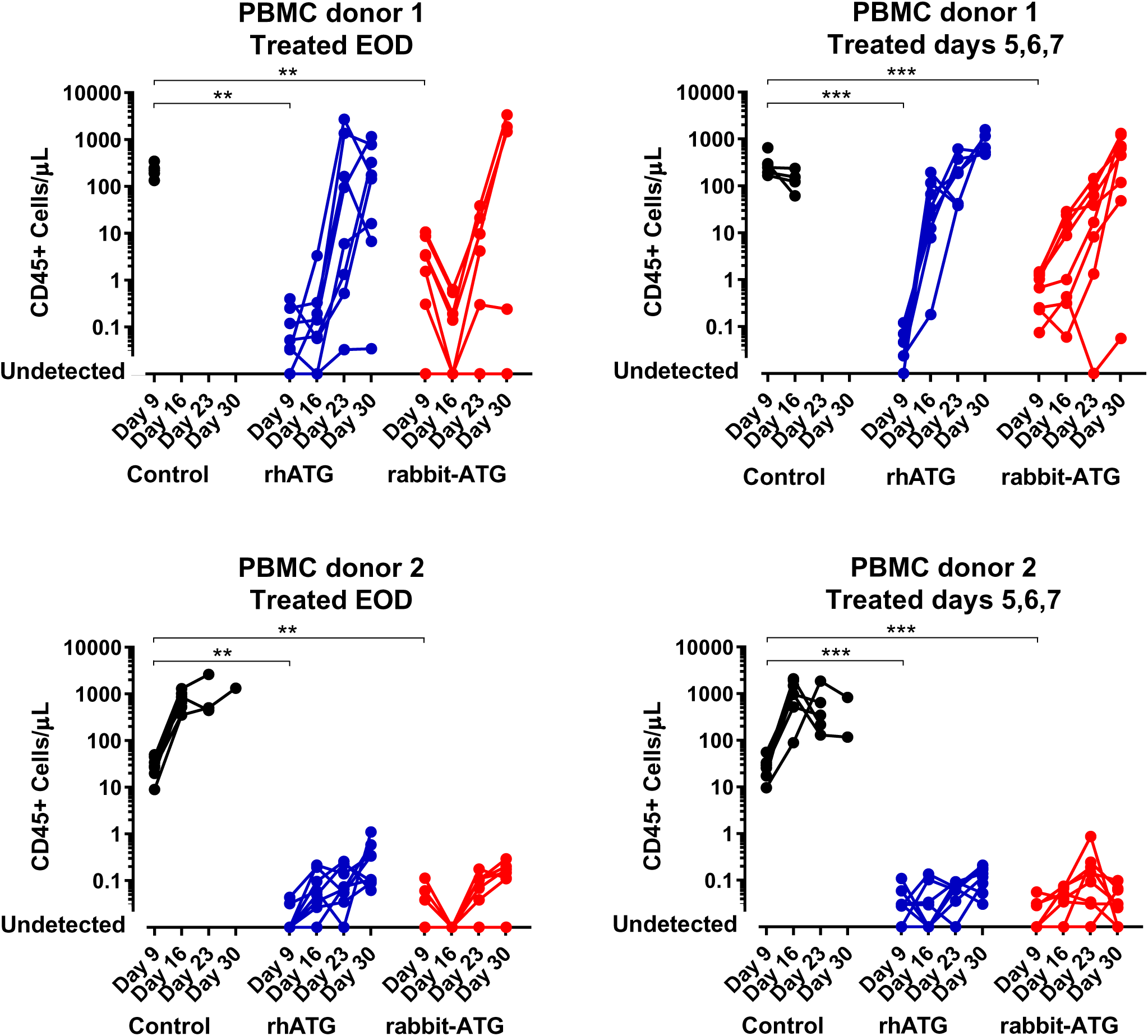
Flow cytometry of CD45+ cells from the ATG GVHD study. Eight animals per treatment group were engrafted with 10^7^ PBMC from one of two donors. Animals were treated with rhATG (blue), rabbit-ATG (red), or vehicle control (black) either every other day beginning at day 5 or on days 5, 6, and 7. Flow cytometry was used to determine the concentration of CD45+ cells from each alive mouse on Days 9, 16, 23, and 30. Lines connect measurements from each mouse. No CD45+ cells were observed where circles intercept the x-axis. ** p<0.01, *** p<0.001.

**Supplementary Figure S27.**
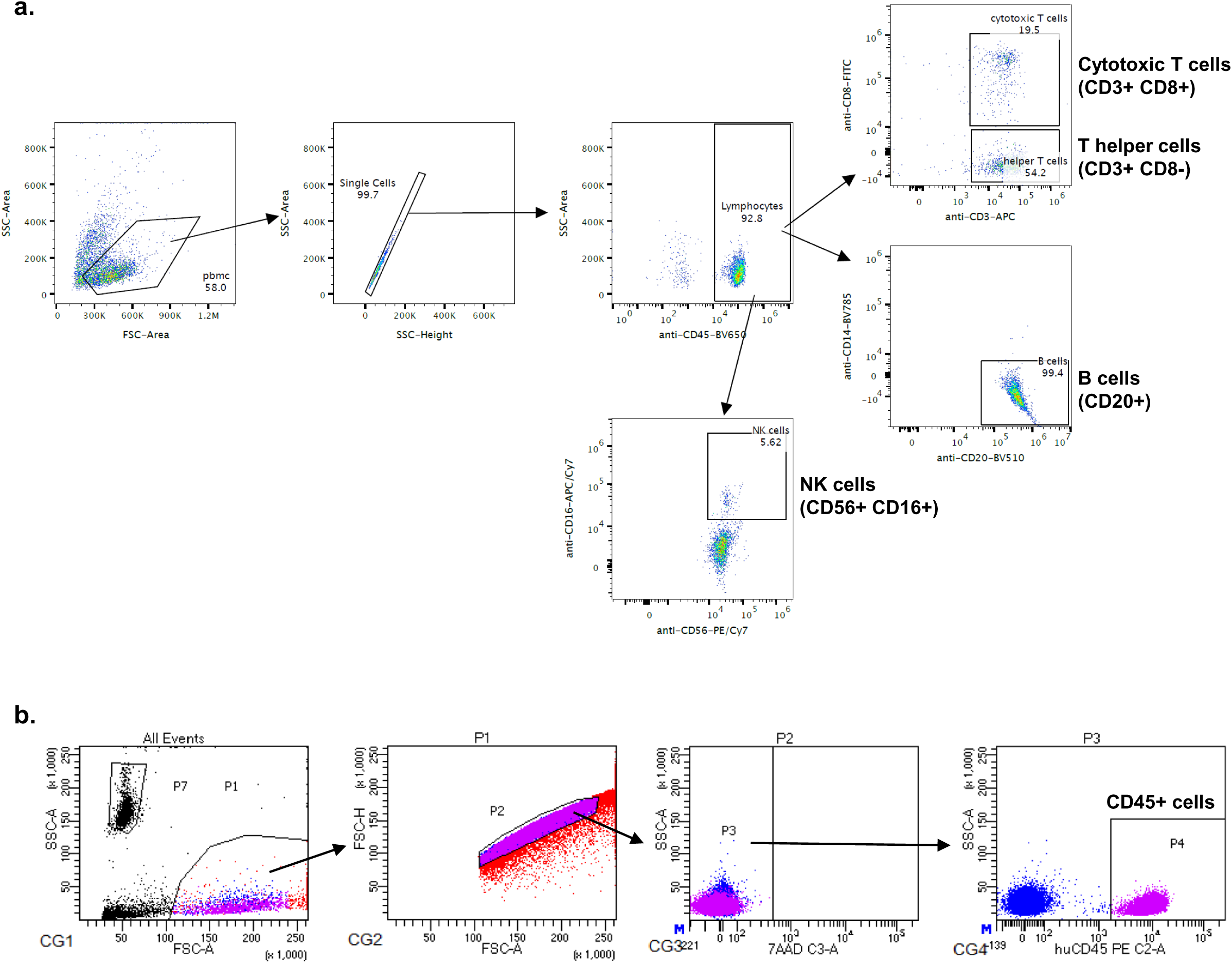
ATG assay flow gating strategies. (**a**) Flow gating strategy for the ATG PBMC killing assay to quantify cytotoxic T cells, T helper cells, B cells, and NK cells. (b) Flow gating strategy of the GVH study to quantify CD45+ cells.

**Table S1.** Characteristics of the 50 donors screened to create rCIG libraries. SOB = shortness of breath; ST = sore throat.

| GigaGen donor |  |  |  |  | Date of | Days between |  | CoV-2 S1 | CoV-2 | CoV-2 RBD | CoV-2 |  |
| --- | --- | --- | --- | --- | --- | --- | --- | --- | --- | --- | --- | --- |
| ID | Race | Sex | Age | Symptoms | symptoms onset | symptoms onset and blood draw | Presumed/Confirmed | binding EC50 (mg/ml) | S1 peak OD signal | binding EC50 (mg/ml) | RBD peak OD signal | Library ID |
| CSS-1902 | AA | Male | 67 | Fever, SOB | 2/28/20 | 32 | Confirmed | 0.0056 | 3.15 | 0.039 | 2.46 | Library 1 |
| CSS-1905 | White | Male | 71 | Fever, Cough, SOB | 3/16/20 | 15 | Presumed | 0.019 | 3.22 | 0.12 | 2.34 | Library 1 |
| CSS-1907 | AA | Female | 66 | Fever, Cough | 3/14/20 | 17 | Presumed | 0.063 | 2.98 | 0.091 | 1.41 | Library 2 |
| CSS-1911 | White | Female | 61 | Fever, ST, SOB | 3/9/20 | 22 | Presumed | 0.10 | 2.80 | 0.13 | 0.75 | Library 2 |
| CSS-1921 | White | Female | 30 | Fever, ST, Cough, SOB | 3/17/20 | 15 | Confirmed | 0.021 | 2.95 | 0.11 | 1.81 | Library 3 |
| CSS-1920 | White | Female | 57 | Fever, Cough, SOB | 3/16/20 | 16 | Confirmed | 0.022 | 2.81 | 0.066 | 1.52 | Library 3 |
| CSS-1928 | White | Female | 63 | Fever, Cough | 3/14/20 | 19 | Presumed | 0.025 | 2.95 | 0.076 | 1.96 | Library 4 |
| CSS-1924 | White | Female | 31 | Fever, Cough, SOB | 3/21/20 | 12 | Confirmed | 0.16 | 2.63 | 0.23 | 2.30 | Library 4 |
| CSS-1944 | White | Female | 49 | Fever, ST, Cough, Pneumonia | 3/15/20 | 24 | Confirmed | 0.014 | 3.21 | 0.053 | 2.17 | Library 5 |
| CSS-1943 | White | Female | 52 | ST, Cough, SOB | 3/16/20 | 23 | Confirmed | 0.014 | 3.15 | 0.067 | 1.85 | Library 5 |
| CSS-1937 | White | Female | 35 | Fever, ST, Cough, SOB | 3/17/20 | 21 | Confirmed | 0.009 | 3.17 | 0.027 | 1.71 | Library 6 |
| CSS-1936 | White | Female | 23 | Fever, Cough, SOB | 3/7/20 | 31 | Confirmed | 0.011 | 3.16 | 0.088 | 2.01 | Library 6 |
| CSS-1901 | White | Male | 67 | Fever, Cough | 3/15/20 | 16 | Confirmed | 0.010 | 3.09 | 0.085 | 2.37 | Library 7 |
| CSS-1940 | White | Female | 40 | Fever, Cough | 3/18/20 | 21 | Confirmed | 0.012 | 3.07 | 0.057 | 1.99 | Library 7 |
| CSS-1939 | White | Female | 24 | Fever, ST, Cough, SOB | 3/8/20 | 30 | Confirmed | 0.037 | 3.10 | 0.073 | 1.37 | Library 8 |
| CSS-1949 | White | Male | 47 | Fever, Cough | 3/21/20 | 19 | Confirmed | 0.13 | 2.79 | 0.32 | 2.32 | Library 8 |
| CSS-1945 | White | Female | 31 | Fever, ST, Cough, SOB | 3/20/20 | 19 | Confirmed | 0.051 | 3.09 | 0.15 | 1.03 |  |
| CSS-1931 | White | Female | 35 | Fever, ST, Cough, SOB | 3/24/20 | 13 | Confirmed | 0.11 | 2.45 | 0.27 | 1.22 |  |
| CSS-1933 | White | Female | 43 | Fever, Cough | 2/25/20 | 42 | Confirmed | 0.13 | 3.23 | 0.27 | 1.16 |  |
| CSS-1932 | White | Female | 33 | Fever, SOB | 3/18/20 | 20 | Confirmed | 0.13 | 3.09 | 0.11 | 1.05 |  |
| CSS-1935 | White | Female | 45 | ST, Cough, SOB | 3/22/20 | 16 | Confirmed | 0.16 | 3.07 | 0.14 | 1.22 |  |
| CSS-1948 | White | Female | 45 | Fever, Cough, SOB | 3/25/20 | 15 | Confirmed | 0.17 | 2.64 | 0.091 | 0.97 |  |
| CSS-1938 | White | Female | 37 | Fever, ST, Cough, SOB | 3/22/20 | 16 | Confirmed | 0.20 | 3.20 | 0.25 | 0.82 |  |
| CSS-1929 | AA | Male | 49 | Fever, Cough, SOB | 2/28/20 | 34 | Presumed | 0.23 | 0.92 | 0.041 | 1.31 |  |
| CSS-1922 | White | Male | 54 | Fever, Cough, SOB | 2/28/20 | 34 | Presumed | 0.24 | 0.34 | 0.14 | 0.45 |  |
| CSS-1914 | White | Female | 65 | Fever, Cough, SOB | 2/26/20 | 35 | Presumed | 0.37 | 1.54 | 0.39 | 2.76 |  |
| CSS-1925 | White | Male | 63 | Fever, Cough, SOB, ST | 3/10/20 | 23 | Presumed | 0.65 | 0.68 | 0.17 | 0.75 |  |
| CSS-1946 | White | Female | 50 | ST, Cough | 3/19/20 | 21 | Confirmed | 0.67 | 2.50 | 0.69 | 1.66 |  |
| CSS-1930 | White | Female | 47 | Fever, Cough | 3/10/20 | 23 | Presumed | 0.69 | 0.45 | 1.29 | 0.94 |  |
| CSS-1934 | White | Female | 42 | Fever, ST, Cough | 3/21/20 | 17 | Confirmed | 0.75 | 3.17 | 0.74 | 1.11 |  |
| CSS-1926 | White | Female | 59 | Fever, Cough, SOB, ST | 3/8/20 | 25 | Presumed | 0.77 | 0.55 | 0.70 | 0.80 |  |
| CSS-1915 | White | Male | 59 | Fever, ST, SOB | 3/11/20 | 21 | Presumed | 0.83 | 0.81 | 0.61 | 0.85 |  |
| CSS-1941 | White | Female | 69 | ST, Cough | 3/11/20 | 28 | Confirmed | 0.91 | 2.93 | 0.55 | 0.94 |  |
| CSS-1917 | White | Male | 67 | Fever, Cough | 3/10/20 | 22 | Presumed | 0.91 | 0.64 | 0.69 | 0.77 |  |
| CSS-1904 | White | Female | 20 | Fever, ST, Cough | 3/11/20 | 20 | Confirmed | 0.96 | 0.64 | 0.63 | 0.86 |  |
| CSS-1916 | White | Male | 70 | Fever, SOB | 2/27/20 | 34 | Presumed | 1.00 | 0.69 | 0.15 | 1.00 |  |
| CSS-1947 | AA | Female | 52 | Fever, Cough | 3/21/20 | 19 | Confirmed | 1.25 | 2.68 | 0.81 | 0.99 |  |
| CSS-1918 | White | Female | 59 | Fever, Cough, SOB | 3/2/20 | 30 | Presumed | 1.32 | 1.55 | 0.38 | 0.91 |  |
| CSS-1923 | White | Female | 57 | Fever, Cough | 2/25/20 | 37 | Presumed | 1.33 | 0.62 | 0.92 | 1.35 |  |
| CSS-1919 | White | Female | 53 | Fever, ST, SOB | 3/13/20 | 19 | Presumed | 1.34 | 0.80 | 0.30 | 0.76 |  |
| CSS-1913 | White | Female | 57 | Fever, Cough | 3/5/20 | 26 | Presumed | 1.47 | 1.69 | 0.23 | 0.88 |  |
| CSS-1927 | AA | Female | 55 | Cough, ST | 3/2/20 | 31 | Presumed | 1.60 | 0.92 | 0.72 | 1.18 |  |
| CSS-1909 | White | Female | 70 | Fever, Cough, SOB | 2/25/20 | 35 | Presumed | 1.88 | 0.92 | 0.13 | 0.75 |  |
| CSS-1912 | White | Female | 46 | Fever, Cough, SOB | 3/8/20 | 23 | Confirmed | 2.20 | 0.76 | 0.10 | 0.67 |  |
| CSS-1906 | White | Male | 58 | Fever, ST, SOB, Cough | 2/25/20 | 35 | Presumed | 2.59 | 1.28 | 1.46 | 1.31 |  |
| CSS-1908 | White | Female | 57 | Fever, Cough, SOB | 2/24/20 | 36 | Presumed | 2.60 | 0.82 | 0.46 | 0.72 |  |
| CSS-1910 | White | Female | 25 | Fever, ST, Cough, SOB | 3/4/20 | 27 | Presumed | 2.84 | 1.27 | 1.12 | 1.23 |  |
| CSS-1942 | White | Female | 39 | Fever, Cough | 3/24/20 | 15 | Confirmed | 2.89 | 2.03 | 1.88 | 2.31 |  |
| CSS-1903 | White | Female | 52 | Fever, Cough, SOB, Fatigue | 3/1/20 | 30 | Presumed | 3.47 | 0.84 | 1.25 | 0.99 |  |
| CSS-1900 | White | Female | 69 | Fever, Cough | 3/6/20 | 25 | Presumed | 9.94 | 0.73 | 3.11 | 1.00 |  |
| IVIG (Gamunex) | --- | --- | --- | --- | --- | --- | --- | 1.41 | 1.99 | 1.27 | 2.26 |  |

**Table S2.** Characterization of the rCIG scFv libraries. Library IDs are from Table S1. Each library plasma is a 1:1 mixture of plasma from the two donors who were used to generate each respective rCIG library. The % binders is determined by yeast scFv flow cytometry. The number of post-sort antibodies was determined by antibody RNA-seq using Illumina sequencing after the 2nd yeast sort.

| Library / Sample | # Input cells | # Ab clones (pre-sort) | # Ab clones (post-sort) | # sequencing reads (pre-sort) | # sequencing reads (post-sort) | % binders (1st sort) | % binders (2nd sort) | % IgG isotype (pre-sort) |  |  |  | % IgG isotype (post-sort) |  |  |  | CoV-2 S1 binding EC50 (mg/ml) | CoV-2 S1 binding fold over plasma avg | CoV-2 RBD binding EC50 (mg/ml) | CoV-2 RBD binding fold over plasma avg | Spike:ACE2 plate-based neutralization IC50 (mg/ml) | Spike:ACE2 plate-based neutralization over plasma avg | CoV-2 pseudotype neutralization IC50 (mg/ml) | CoV-2 pseudotype neutralization over plasma avg | Live CoV-2 virus neutralization concentration (mg/ml) | Live CoV-2 virus neutralization over plasma avg |
| --- | --- | --- | --- | --- | --- | --- | --- | --- | --- | --- | --- | --- | --- | --- | --- | --- | --- | --- | --- | --- | --- | --- | --- | --- | --- |
|  |  |  |  |  |  |  |  | IgG1 | IgG2 | IgG3 | IgG4 | IgG1 | IgG2 | IgG3 | IgG4 |  |  |  |  |  |  |  |  |  |  |
| Sorted Library 1 | 3,400,000 | 87,902 | 3,971 | 674,234 | 58,556 | 1.02 | 43.7 | 83.4 | 8.9 | 7.4 | 0.3 | 88.9 | 8.6 | 2.6 | 0.0 | 0.000072 | 335 | 0.000053 | 447 | 0.0026 | 288 | 0.00049 | 264 | 0.0032 | 17 |
| Library 1 plasma | --- | --- | --- | --- | --- | --- | --- | --- | --- | --- | --- | --- | --- | --- | --- | 0.013 | 1.9 | 0.013 | 1.8 | 0.7 | 1.1 | 0.015 | 8.6 | 0.016 | 3.3 |
| Sorted Library 2 | 3,580,000 | 54,986 | 1,730 | 594,209 | 64,730 | 0.57 | 35.8 | 61.1 | 21.4 | 16.2 | 1.3 | 63.6 | 28.8 | 3.0 | 4.6 | 0.00014 | 172 | 0.0001 | 237 | 0.0052 | 144 | 0.0048 | 27 | 0.0032 | 17 |
| Library 2 plasma | --- | --- | --- | --- | --- | --- | --- | --- | --- | --- | --- | --- | --- | --- | --- | 0.069 | 0.3 | 0.044 | 0.5 | 1.28 | 0.6 | 0.76 | 0.2 | 0.166 | 0.3 |
| Sorted Library 3 | 3,490,000 | 67,805 | 3,042 | 612,720 | 67,669 | 1.6 | 49.3 | 68.1 | 14.1 | 17.4 | 0.4 | 83.9 | 14.4 | 1.7 | 0.0 | 0.000052 | 464 | 0.000047 | 504 | 0.0018 | 416 | 0.0018 | 72 | 0.0047 | 11 |
| Library 3 plasma | --- | --- | --- | --- | --- | --- | --- | --- | --- | --- | --- | --- | --- | --- | --- | 0.024 | 1.0 | 0.034 | 0.7 | 0.28 | 2.7 | 0.056 | 2.3 | 0.032 | 1.7 |
| Sorted Library 4 | 3,230,000 | 74,074 | 3,069 | 708,240 | 76,274 | 0.42 | 27.4 | 55.4 | 9.9 | 33.5 | 1.3 | 72.3 | 18.1 | 9.0 | 0.6 | 0.000054 | 447 | 0.000058 | 409 | 0.0017 | 440 | 0.002 | 65 | 0.002 | 26 |
| Library 4 plasma | --- | --- | --- | --- | --- | --- | --- | --- | --- | --- | --- | --- | --- | --- | --- | 0.027 | 0.9 | 0.035 | 0.7 | 2.4 | 0.3 | 0.031 | 4.2 | 0.038 | 1.4 |
| Sorted Library 5 | 3,790,000 | 156,592 | 2,122 | 717,789 | 34,397 | 2.29 | 47.8 | 54.7 | 22.0 | 22.8 | 0.6 | 80.8 | 14.1 | 5.1 | 0.0 | 0.000066 | 366 | 0.000061 | 389 | 0.0016 | 468 | 0.00082 | 158 | 0.0017 | 31 |
| Library 5 plasma | --- | --- | --- | --- | --- | --- | --- | --- | --- | --- | --- | --- | --- | --- | --- | 0.017 | 1.4 | 0.021 | 1.1 | 0.28 | 2.7 | 0.049 | 2.6 | 0.042 | 1.3 |
| Sorted Library 6 | 3,900,000 | 62,368 | 1,656 | 765,707 | 25,234 | 0.95 | 26.6 | 64.2 | 16.6 | 17.8 | 1.3 | 85.1 | 13.4 | 1.5 | 0.0 | 0.000049 | 492 | 0.000054 | 439 | 0.0016 | 468 | 0.00034 | 381 | 0.00043 | 123 |
| Library 6 plasma | --- | --- | --- | --- | --- | --- | --- | --- | --- | --- | --- | --- | --- | --- | --- | 0.012 | 2.0 | 0.017 | 1.4 | 0.38 | 2.0 | 0.039 | 3.3 | 0.045 | 1.2 |
| Sorted Library 7 | 2,950,000 | 67,212 | 1,768 | 846,270 | 34,556 | 1.37 | 34.7 | 74.2 | 15.5 | 10.0 | 0.3 | 92.2 | 5.2 | 2.6 | 0.0 | 0.000042 | 574 | 0.000044 | 539 | 0.023 | 33 | 0.00032 | 404 | 0.00095 | 56 |
| Library 7 plasma | --- | --- | --- | --- | --- | --- | --- | --- | --- | --- | --- | --- | --- | --- | --- | 0.012 | 2.0 | 0.0096 | 2.5 | 0.32 | 2.3 | 0.024 | 5.4 | 0.018 | 2.9 |
| Sorted Library 8 | 3,880,000 | 116,642 | 1,214 | 783,140 | 30,044 | 0.88 | 24.9 | 51.5 | 19.0 | 26.5 | 3.0 | 75.9 | 16.7 | 5.6 | 1.9 | 0.000028 | 862 | 0.000027 | 878 | 0.0016 | 468 | 0.00017 | 761 | 0.00048 | 110 |
| Library 8 plasma | --- | --- | --- | --- | --- | --- | --- | --- | --- | --- | --- | --- | --- | --- | --- | 0.019 | 1.3 | 0.016 | 1.5 | 0.35 | 2.1 | 0.061 | 2.1 | 0.066 | 0.8 |
| IVIG (Gamunex) | --- | --- | --- | --- | --- | --- | --- | --- | --- | --- | --- | --- | --- | --- | --- | 1.38 | --- | 1.25 | --- | None | --- | None | --- | None | --- |
| SARS CoV mAb [CR3022] | --- | --- | --- | --- | --- | --- | --- | --- | --- | --- | --- | --- | --- | --- | --- | 0.000078 | --- | 0.000088 | --- | None | --- | 0.1 | --- | Not tested | --- |
| SARS CoV-2 mAb [SAD-S35] | --- | --- | --- | --- | --- | --- | --- | --- | --- | --- | --- | --- | --- | --- | --- | 0.000058 | --- | 0.000057 | --- | 0.0011 | --- | 0.00026 | --- | Not tested | --- |

**Table S3.**
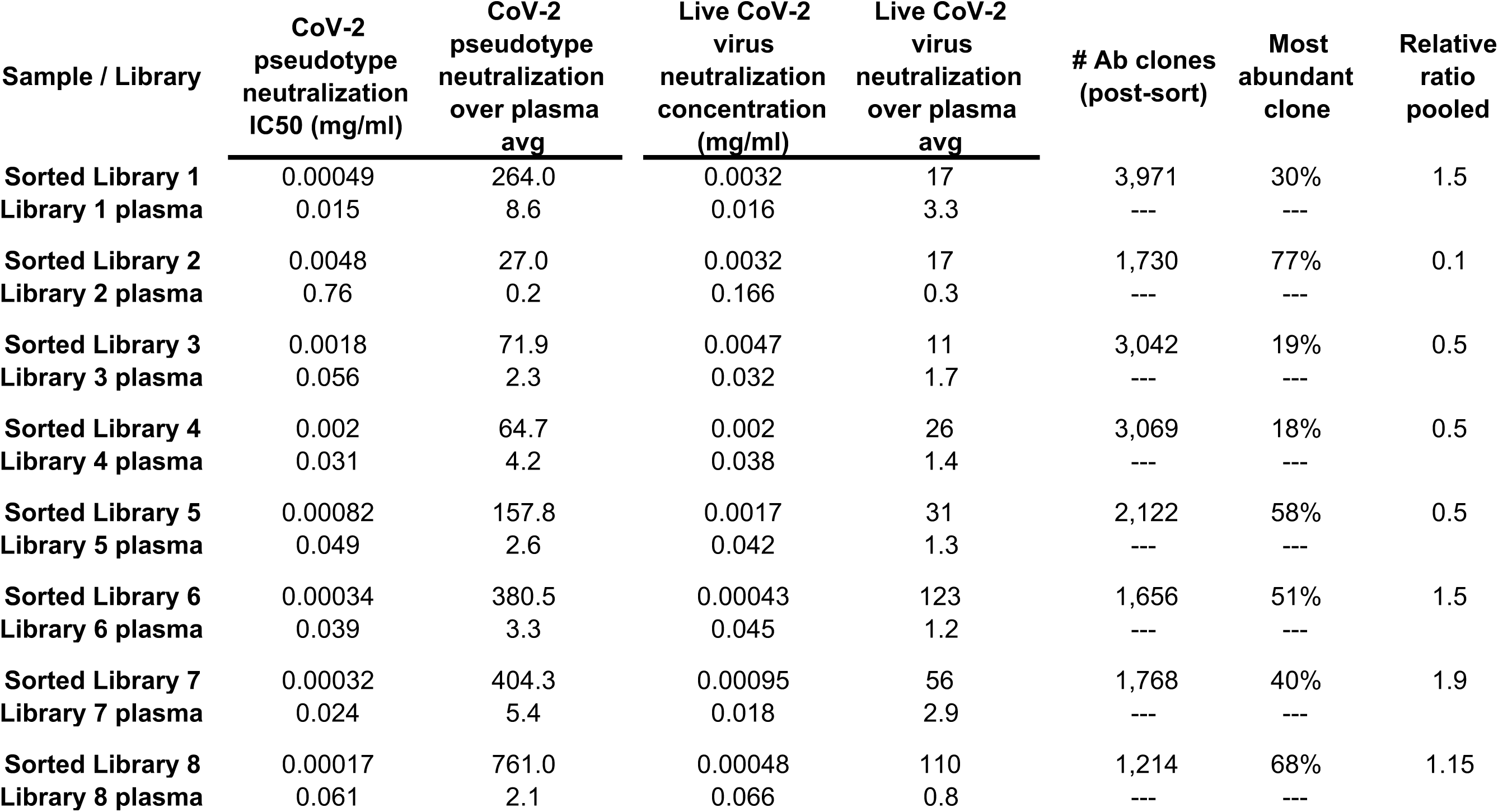
The relative ratio that each of the eight sorted rCIG libraries was mixed is indicated.

| Sample / Library | CoV-2<br>pseudotype<br>neutralization<br>IC50 (mg/ml) | CoV-2<br>pseudotype<br>neutralization<br>over plasma<br>avg | Live CoV-2<br>virus<br>neutralization<br>concentration<br>(mg/ml) | Live CoV-2<br>virus<br>neutralization<br>over plasma<br>avg | # Ab clones<br>(post-sort) | Most<br>abundant<br>clone | Relative<br>ratio<br>pooled |
| --- | --- | --- | --- | --- | --- | --- | --- |
| Sorted Library 1 | 0.00049 | 264.0 | 0.0032 | 17 | 3,971 | 30% | 1.5 |
| Library 1 plasma | 0.015 | 8.6 | 0.016 | 3.3 | --- | --- |  |
| Sorted Library 2 | 0.0048 | 27.0 | 0.0032 | 17 | 1,730 | 77% | 0.1 |
| Library 2 plasma | 0.76 | 0.2 | 0.166 | 0.3 | --- | --- |  |
| Sorted Library 3 | 0.0018 | 71.9 | 0.0047 | 11 | 3,042 | 19% | 0.5 |
| Library 3 plasma | 0.056 | 2.3 | 0.032 | 1.7 | --- | --- |  |
| Sorted Library 4 | 0.002 | 64.7 | 0.002 | 26 | 3,069 | 18% | 0.5 |
| Library 4 plasma | 0.031 | 4.2 | 0.038 | 1.4 | --- | --- |  |
| Sorted Library 5 | 0.00082 | 157.8 | 0.0017 | 31 | 2,122 | 58% | 0.5 |
| Library 5 plasma | 0.049 | 2.6 | 0.042 | 1.3 | --- | --- |  |
| Sorted Library 6 | 0.00034 | 380.5 | 0.00043 | 123 | 1,656 | 51% | 1.5 |
| Library 6 plasma | 0.039 | 3.3 | 0.045 | 1.2 | --- | --- |  |
| Sorted Library 7 | 0.00032 | 404.3 | 0.00095 | 56 | 1,768 | 40% | 1.9 |
| Library 7 plasma | 0.024 | 5.4 | 0.018 | 2.9 | --- | --- |  |
| Sorted Library 8 | 0.00017 | 761.0 | 0.00048 | 110 | 1,214 | 68% | 1.15 |
| Library 8 plasma | 0.061 | 2.1 | 0.066 | 0.8 | --- | --- |  |

**Table S4.**
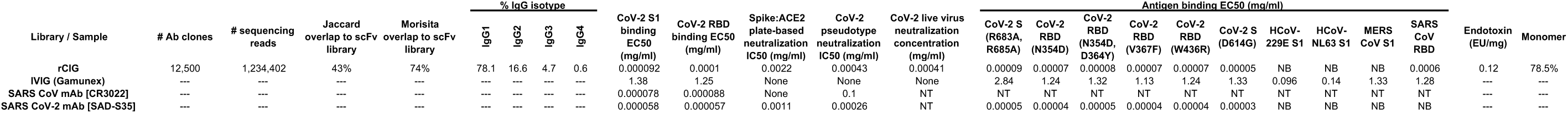
Characteristics of the rCIG CHO cell library. The IgG isotype refers to what isotype the variable region was prior to conversion of the sequence to IgG1 for expression in CHO. NB, no binding observed (expected to be similar to IVIG if a higher concentration could be tested). NT, not tested

**Table S5.** Characterization of the rZIG scFv library.

| **Library** | **# input cells** | **# Ab clones** | **# sequencing reads** |
| --- | --- | --- | --- |
| **rZIG** | 2,080,000 | 119,700 | 1,128,316 |

**Table S6.** Characteristics of the rZIG CHO cell libraries. NT, Not tested

| Library / Sample | # Ab clones | # sequencing reads | Jaccard overlap to scFv library | Morisita overlap to scFv library | Jaccard overlap between rZIG-IgG1 and rZIG-LALA | Morisita overlap between rZIG-IgG1 and rZIG-LALA | Binding EC50 (µg/ml) |  |  |  |  | Neutralization IC50 (µg/ml) |  |  |  |  | Endotoxin (EU/mg) | Monomer |
| --- | --- | --- | --- | --- | --- | --- | --- | --- | --- | --- | --- | --- | --- | --- | --- | --- | --- | --- |
|  |  |  |  |  |  |  | Zika | Dengue serotype 1 | Dengue serotype 2 | Dengue serotype 3 | Dengue serotype 4 | Zika | Dengue serotype 1 | Dengue serotype 2 | Dengue serotype 3 | Dengue serotype 4 |  |  |
| rZIG-IgG1 | 33,642 | 1,580,084 | 40% | 56% | 58% | 86% | 1.13 | 0.89 | 1.55 | 1.21 | 1.31 | 0.28 | 7.24 | 1.01 | 5.88 | 1.51 | <0.04 | 97.8% |
| rZIG-LALA | 26,708 | 1,500,424 | 35% | 37% |  |  | 1.77 | 1.46 | 2.60 | 1.79 | 2.61 | 0.33 | 21.5 | 0.39 | 3.86 | 2.58 | <0.09 | 97% |
| IVIG | --- | --- | --- | --- | --- | --- | None | None | None | None | None | None | 12.2 | None | None | None | --- | --- |
| Zika/Dengue+ serum | --- | --- | --- | --- | --- | --- | 141.8 | 133.8 | 128.1 | 131.8 | 135.6 | 34.7 | 2.5 | 4.94 | 6.79 | 5.39 | --- | --- |
| Zika/Dengue+ mAb | --- | --- | --- | --- | --- | --- | NT | NT | NT | NT | NT | 0.098 | 0.048 | 0.21 | 0.28 | 0.12 | --- | --- |

**Table S7.**
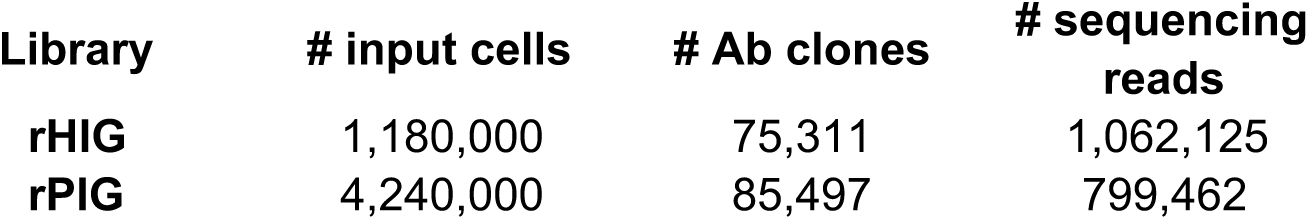
Characterization of the rHIG and rPIG scFv libraries.

| **Library** | **# input cells** | **# Ab clones** | **# sequencing reads** |
| --- | --- | --- | --- |
| **rHIG** | 1,180,000 | 75,311 | 1,062,125 |
| **rPIG** | 4,240,000 | 85,497 | 799,462 |

**Table S8.**
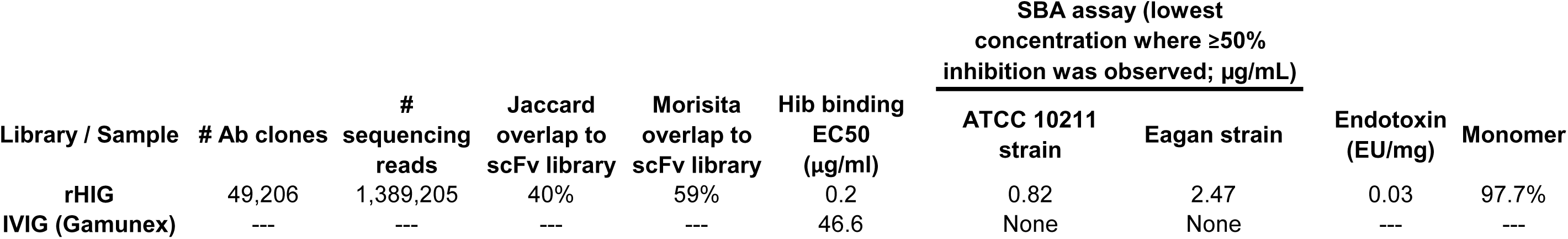
Characteristics of the rHIG CHO cell library. SBA, serum bactericidal assay

| Library / Sample | # Ab clones | # sequencing reads | Jaccard overlap to scFv library | Morisita overlap to scFv library | Hib binding EC50 (µg/ml) | SBA assay (lowest concentration where ≥50% inhibition was observed; µg/mL) |  | Endotoxin (EU/mg) | Monomer |
| --- | --- | --- | --- | --- | --- | --- | --- | --- | --- |
|  |  |  |  |  |  | ATCC 10211 strain | Eagan strain |  |  |
| rHIG | 49,206 | 1,389,205 | 40% | 59% | 0.2 | 0.82 | 2.47 | 0.03 | 97.7% |
| IVIG (Gamunex) | --- | --- | --- | --- | 46.6 | None | None | --- | --- |

**Table S9.** Characteristics of the rPIG CHO cell library.

| Library / Sample | # Ab clones | # sequencing reads | Jaccard overlap to scFv library | Morisita overlap to scFv library | Binding EC50 to pool of 23 pneumococcus serotypes | Concentration of pneumococcus serotype-specific antibodies (µg/mg) |  |  |  |  |  |  |  |  |  |  |  |  |  |  |  | Pneumococcus serotype-specific opsonophagocytosis index |  |  |  |  |  |  |  |  |  |  |  |  |  |  |  | Endotoxin (EU/mg) | Monomer |
| --- | --- | --- | --- | --- | --- | --- | --- | --- | --- | --- | --- | --- | --- | --- | --- | --- | --- | --- | --- | --- | --- | --- | --- | --- | --- | --- | --- | --- | --- | --- | --- | --- | --- | --- | --- | --- | --- | --- | --- |
|  |  |  |  |  |  | 1 | 3 | 4 | 5 | 6A | 6B | 7F | 9V | 12F | 14 | 18C | 19A | 19F | 22F | 23F | 33F | 1 | 3 | 4 | 5 | 6A | 6B | 7F | 9V | 12F | 14 | 18C | 19A | 19F | 22F | 23F | 33F |  |  |
| rPIG | 17,938 | 1,450,945 | 41% | 77% | 0.055 | 21.7 | 6.6 | 0.4 | 2.2 | none | 0.2 | none | 5.0 | 1.0 | 53.1 | 4.1 | 5.5 | 5.1 | 1.3 | 0.5 | 10.1 | 8 | 277 | 76 | 61 | 17 | 15 | 189 | 56 | 620 | 727 | 357 | 72 | 72 | 96 | 45 | 147 | 0.03 | 98.6% |
| IVIG (Gamunex) | --- | --- | --- | --- | 4.7 | 0.11 | 0.05 | 0.05 | 0.11 | 0.15 | 0.13 | 0.11 | 0.13 | 0.05 | 0.47 | 0.11 | 0.39 | 0.23 | 0.14 | 0.09 | 0.20 | 8 | 8 | 11 | 8 | 26 | 15 | 47 | 56 | 23 | 46 | 66 | 20 | 13 | 39 | 27 | 57 | --- | --- |
|  |  |  |  | Fold over IVIG: | 85 | 203 | 140 | 7 | 20 | --- | 1.7 | --- | 39 | 20 | 113 | 36 | 14 | 22 | 9.5 | 5.5 | 50 | 1 | 35 | 6.9 | 7.6 | 0.7 | 1 | 4 | 1 | 27 | 16 | 5.4 | 3.6 | 5.5 | 2.5 | 1.7 | 2.6 |  |  |

**Table S10.** Characteristics of the IVIG + rHIG/rPIG mixture.

| **Library / Sample** | **Hib binding**<br>**EC50 (µg/ml)** | **Binding EC50 to**<br>**pool of 23**<br>**pneumococcus**<br>**serotypes (µg/ml)** |
| --- | --- | --- |
| **IVIG + rHIG/rPIG** | 2.54 | 0.3 |
| **IVIG (Gamunex)** | 46.6 | 2.49 |
| ***Fold over IVIG:*** | **18.3** | **8.3** |

**Table S11.** Characterization of the rhATG scFv libraries.

| **Library** | **# input cells** | **# Ab clones** | **# sequencing reads** |
| --- | --- | --- | --- |
| rhATG (T cells; bone marrow) | 150,000 | 13,314 | 807,164 |
| rhATG (T cells; lymph node) | 1,640,000 | 34,324 | 1,189,267 |
| rhATG (Thymocytes; lymph node) | 2,060,000 | 22,030 | 632,083 |
| rhATG (Thymocytes; spleen) | 1,730,000 | 32,604 | 923,203 |

**Table S12.** Characteristics of the rhATG CHO cell library.

| Library | # Ab clones | # sequencing reads | Jaccard overlap to scFv library (average of the 4 scFv libraries) | Morisita overlap to scFv library (average of the 4 scFv libraries) | Endotoxin (EU/mg) | Monomer |
| --- | --- | --- | --- | --- | --- | --- |
| rhATG | 49,885 | 5,324,998 | 35% | 86% | <0.07 | 99.1% |

## REFERENCES

1 Bozzo J, Jorquera JI. Use of human immunoglobulins as an anti-infective treatment: the experience so far and their possible re-emerging role. Expert Rev Anti Infect Ther. 2017;15(6):585–604. doi:10.1080/14787210.2017.1328278

2 Beasley RP, Hwang LY, Stevens CE, et al. Efficacy of hepatitis B immune globulin for prevention of perinatal transmission of the hepatitis B virus carrier state: final report of a randomized double-blind, placebo-controlled trial. Hepatology. 1983;3(2):135–141. doi:10.1002/hep.1840030201

3 Payne JR, Khouri JM, Jewell NP, Arnon SS. Efficacy of Human Botulism Immune Globulin for the Treatment of Infant Botulism: The First 12 Years Post Licensure. J Pediatr. 2018;193:172–177. doi:10.1016/j.jpeds.2017.10.035

4 Gaber AO, First MR, Tesi RJ, et al. Results of the double-blind, randomized, multicenter, phase III clinical trial of Thymoglobulin versus Atgam in the treatment of acute graft rejection episodes after renal transplantation. Transplantation. 1998;66(1):29–37. doi:10.1097/00007890-199807150-00005

5 Branche E, Simon AY, Sheets N, et al. Human Polyclonal Antibodies Prevent Lethal Zika Virus Infection in Mice. Sci Rep. 2019;9(1):9857. Published 2019 Jul 8. doi:10.1038/s41598-019-46291-9

6 Bloch EM, Shoham S, Casadevall A, et al. Deployment of convalescent plasma for the prevention and treatment of COVID-19. J Clin Invest. 2020;130(6):2757–2765. doi:10.1172/JCI138745

7 Shen C, Wang Z, Zhao F, et al. Treatment of 5 Critically Ill Patients With COVID-19 With Convalescent Plasma [published online ahead of print, 2020 Mar 27]. JAMA. 2020;323(16):1582–1589. doi:10.1001/jama.2020.4783

8 Boulis A, Goold S, Ubel PA. Responding to the immunoglobulin shortage: a case study. J Health Polit Policy Law. 2002;27(6):977–999. doi:10.1215/03616878-27-6-977

9 Bjøro K, Frøland SS, Yun Z, Samdal HH, Haaland T. Hepatitis C infection in patients with primary hypogammaglobulinemia after treatment with contaminated immune globulin. N Engl J Med. 1994;331(24):1607–1611. doi:10.1056/NEJM199412153312402

10 Etscheid M, Breitner-Ruddock S, Gross S, Hunfeld A, Seitz R, Dodt J. Identification of kallikrein and FXIa as impurities in therapeutic immunoglobulins: implications for the safety and control of intravenous blood products. Vox Sang. 2012;102(1):40–46. doi:10.1111/j.1423-0410.2011.01502.x

11 Brabant S, Facon A, Provôt F, Labalette M, Wallaert B, Chenivesse C. An avoidable cause of thymoglobulin anaphylaxis. Allergy Asthma Clin Immunol. 2017;13:13. Published 2017 Feb 23. doi:10.1186/s13223-017-0186-9

12 Ober RJ, Radu CG, Ghetie V, Ward ES. Differences in promiscuity for antibody-FcRn interactions across species: implications for therapeutic antibodies. Int Immunol. 2001;13(12):1551–1559. doi:10.1093/intimm/13.12.1551

13 Simon HU, Späth PJ. IVIG--mechanisms of action. Allergy. 2003;58(7):543–552. doi:10.1034/j.1398-9995.2003.00239.x

14 Lejtenyi D, Mazer B. Consistency of protective antibody levels across lots of intravenous immunoglobulin preparations. J Allergy Clin Immunol. 2008;121(1):254–255. doi:10.1016/j.jaci.2007.11.001

15 Popow I, Leitner J, Majdic O, et al. Assessment of batch to batch variation in polyclonal antithymocyte globulin preparations. Transplantation. 2012;93(1):32–40. doi:10.1097/TP.0b013e31823bb664

16 Rita Costa A, Elisa Rodrigues M, Henriques M, Azeredo J, Oliveira R. Guidelines to cell engineering for monoclonal antibody production. Eur J Pharm Biopharm. 2010;74(2):127–138. doi:10.1016/j.ejpb.2009.10.002

17 Frandsen TP, Naested H, Rasmussen SK, et al. Consistent manufacturing and quality control of a highly complex recombinant polyclonal antibody product for human therapeutic use. Biotechnol Bioeng. 2011;108(9):2171–2181. doi:10.1002/bit.23166

18 Adler AS, Mizrahi RA, Spindler MJ, et al. Rare, high-affinity anti-pathogen antibodies from human repertoires, discovered using microfluidics and molecular genomics. MAbs. 2017;9(8):1282–1296. doi:10.1080/19420862.2017.1371383

19 Soto C, Bombardi RG, Branchizio A, et al. High frequency of shared clonotypes in human B cell receptor repertoires. Nature. 2019;566(7744):398–402. doi:10.1038/s41586-019-0934-8

20 Gibson DG, Young L, Chuang RY, Venter JC, Hutchison CA 3rd, Smith HO. Enzymatic assembly of DNA molecules up to several hundred kilobases. Nat Methods. 2009;6(5):343–345. doi:10.1038/nmeth.1318

21 Kito M, Itami S, Fukano Y, Yamana K, Shibui T. Construction of engineered CHO strains for high-level production of recombinant proteins. Appl Microbiol Biotechnol. 2002;60(4):442–448. doi:10.1007/s00253-002-1134-1

22 Iversen PL, Kane CD, Zeng X, et al. Recent successes in therapeutics for Ebola virus disease: no time for complacency [published online ahead of print, 2020 Jun 18]. Lancet Infect Dis. 2020;S1473-3099(20)30282–6. doi:10.1016/S1473-3099(20)30282-6

23 Guan Y, Zheng BJ, He YQ, et al. Isolation and characterization of viruses related to the SARS coronavirus from animals in southern China. Science. 2003;302(5643):276–278. doi:10.1126/science.1087139

24 Zaki AM, van Boheemen S, Bestebroer TM, Osterhaus AD, Fouchier RA. Isolation of a novel coronavirus from a man with pneumonia in Saudi Arabia [published correction appears in N Engl J Med. 2013 Jul 25;369(4):394]. N Engl J Med. 2012;367(19):1814–1820. doi:10.1056/NEJMoa1211721

25 Itoh Y, Shinya K, Kiso M, et al. In vitro and in vivo characterization of new swine-origin H1N1 influenza viruses. Nature. 2009;460(7258):1021–1025. doi:10.1038/nature08260

26 Wikan N, Smith DR. Zika virus: history of a newly emerging arbovirus. Lancet Infect Dis. 2016;16(7):e119–e126. doi:10.1016/S1473-3099(16)30010-X

27 Zhou P, Yang XL, Wang XG, et al. A pneumonia outbreak associated with a new coronavirus of probable bat origin. Nature. 2020;579(7798):270–273. doi:10.1038/s41586-020-2012-7

28 Henao-Restrepo AM, Camacho A, Longini IM, et al. Efficacy and effectiveness of an rVSV-vectored vaccine in preventing Ebola virus disease: final results from the Guinea ring vaccination, open-label, cluster-randomised trial (Ebola Ça Suffit!) [published correction appears in Lancet. 2017 Feb 4;389(10068):504] [published correction appears in Lancet. 2017 Feb 4;389(10068):504]. Lancet. 2017;389(10068):505–518. doi:10.1016/S0140-6736(16)32621-6

29 Mlakar J, Korva M, Tul N, et al. Zika Virus Associated with Microcephaly. N Engl J Med. 2016;374(10):951–958. doi:10.1056/NEJMoa1600651

30 Katzelnick LC, Gresh L, Halloran ME, et al. Antibody-dependent enhancement of severe dengue disease in humans. Science. 2017;358(6365):929–932. doi:10.1126/science.aan6836

31 Priyamvada L, Quicke KM, Hudson WH, et al. Human antibody responses after dengue virus infection are highly cross-reactive to Zika virus. Proc Natl Acad Sci U S A. 2016;113(28):7852–7857. doi:10.1073/pnas.1607931113

32 Stettler K, Beltramello M, Espinosa DA, et al. Specificity, cross-reactivity, and function of antibodies elicited by Zika virus infection. Science. 2016;353(6301):823–826. doi:10.1126/science.aaf8505

33 Langerak T, Mumtaz N, Tolk VI, et al. The possible role of cross-reactive dengue virus antibodies in Zika virus pathogenesis. PLoS Pathog. 2019;15(4):e1007640. doi:10.1371/journal.ppat.1007640

34 Hessell AJ, Hangartner L, Hunter M, et al. Fc receptor but not complement binding is important in antibody protection against HIV. Nature. 2007;449(7158):101–104. doi:10.1038/nature06106

35 Gathmann B, Mahlaoui N; CEREDIH, et al. Clinical picture and treatment of 2212 patients with common variable immunodeficiency. J Allergy Clin Immunol. 2014;134(1):116–126. doi:10.1016/j.jaci.2013.12.1077

36 Sperlich JM, Grimbacher B, Workman S, et al. Respiratory Infections and Antibiotic Usage in Common Variable Immunodeficiency. J Allergy Clin Immunol Pract. 2018;6(1):159–168.e3. doi:10.1016/j.jaip.2017.05.024

37 Orange JS, Grossman WJ, Navickis RJ, Wilkes MM. Impact of trough IgG on pneumonia incidence in primary immunodeficiency: A meta-analysis of clinical studies. Clin Immunol. 2010;137(1):21–30. doi:10.1016/j.clim.2010.06.012

38 Geno KA, Gilbert GL, Song JY, et al. Pneumococcal Capsules and Their Types: Past, Present, and Future. Clin Microbiol Rev. 2015;28(3):871–899. doi:10.1128/CMR.00024-15

39 Hart A, Smith JM, Skeans MA, Gustafson SK, Wilk AR, Robinson A, Wainright JL, Haynes CR, Snyder JJ, Kasiske BL, Israni AK. OPTN/SRTR 2016 Annual Data Report: Kidney. Am J Transplant. 2018;18 Suppl 1:18–113. doi: 10.1111/ajt.14557.

40 Brennan DC, Daller JA, Lake KD, Cibrik D, Del Castillo D, Thymoglobulin Induction Study G. Rabbit antithymocyte globulin versus basiliximab in renal transplantation. N Engl J Med. 2006;355(19):1967–77. doi: 10.1056/NEJMoa060068.

41 Popow I, Leitner J, Grabmeier-Pfistershammer K, Majdic O, Zlabinger GJ, Kundi M, Steinberger P. A comprehensive and quantitative analysis of the major specificities in rabbit antithymocyte globulin preparations. Am J Transplant. 2013;13(12):3103–13. doi: 10.1111/ajt.12514.

42 Norelli M, Camisa B, Bondanza A. Modeling Human Graft-Versus-Host Disease in Immunocompromised Mice. Methods Mol Biol. 2016;1393:127–132. doi:10.1007/978-1-4939-3338-9_12

43 Spindler MJ, Nelson AL, Wagner EK, et al. Massively parallel interrogation and mining of natively paired human TCRαβ repertoires. Nat Biotechnol. 2020;38(5):609–619. doi:10.1038/s41587-020-0438-y

44 Lee J, Boutz DR, Chromikova V, et al. Molecular-level analysis of the serum antibody repertoire in young adults before and after seasonal influenza vaccination. Nat Med. 2016;22(12):1456–1464. doi:10.1038/nm.4224

45 Vollmers C, Sit RV, Weinstein JA, Dekker CL, Quake SR. Genetic measurement of memory B-cell recall using antibody repertoire sequencing. Proc Natl Acad Sci U S A. 2013;110(33):13463–13468. doi:10.1073/pnas.1312146110

46 Plesa C, Sidore AM, Lubock NB, Zhang D, Kosuri S. Multiplexed gene synthesis in emulsions for exploring protein functional landscapes. Science. 2018;359(6373):343–347. doi:10.1126/science.aao5167

47 Medina-Cucurella AV, Mizrahi RA, Asensio MA, et al. Preferential Identification of Agonistic OX40 Antibodies by Using Cell Lysate to Pan Natively Paired, Humanized Mouse-Derived Yeast Surface Display Libraries. Antibodies (Basel). 2019;8(1):17. doi:10.3390/antib8010017

48 Edgar RC. Search and clustering orders of magnitude faster than BLAST. Bioinformatics. 2010 Oct 1;26(19):2460–1. doi: 10.1093/bioinformatics/btq461.

49 Csardi G, Nepusz T, The igraph software package for complex network research. InterJournal 2006; Complex Systems:1695.

50 Nazarov VI, Pogorelyy MV, Komech EA, et al. tcR: an R package for T cell receptor repertoire advanced data analysis. BMC Bioinformatics. 2015;16(1):175. doi:10.1186/s12859-015-0613-1

51 Amanat F, Stadlbauer D, Strohmeier S, et al. A serological assay to detect SARS-CoV-2 seroconversion in humans [published online ahead of print, 2020 May 12]. Nat Med. 2020;10.1038/s41591-020-0913-5. doi:10.1038/s41591-020-0913-5

52 Williamson PC, Linnen JM, Kessler DA, et al. First cases of Zika virus-infected US blood donors outside states with areas of active transmission. Transfusion. 2017;57(3pt2):770–778. doi:10.1111/trf.14041

53 Plikaytis BD, Goldblatt D, Frasch CE, et al. An analytical model applied to a multicenter pneumococcal enzyme-linked immunosorbent assay study. J Clin Microbiol. 2000;38(6):2043–2050.

54 Wernette CM, Frasch CE, Madore D, et al. Enzyme-linked immunosorbent assay for quantitation of human antibodies to pneumococcal polysaccharides. Clin Diagn Lab Immunol. 2003;10(4):514–519. doi:10.1128/cdli.10.4.514-519.2003

55 Rose CE, Romero-Steiner S, Burton RL, et al. Multilaboratory comparison of Streptococcus pneumoniae opsonophagocytic killing assays and their level of agreement for the determination of functional antibody activity in human reference sera. Clin Vaccine Immunol. 2011;18(1):135–142. doi:10.1128/CVI.00370-10

